# Novel graph theoretic biological pathway network analytics methods for analyzing and discovering Alzheimer’s disease related genes

**DOI:** 10.1101/2021.10.19.465019

**Authors:** Subrata Saha, Ahmed Soliman, Sanguthevar Rajasekaran

## Abstract

**Background:** Alzheimer’s disease (AD) is the most common form of dementia among older people. It is a complex disease and the genetics and environmental factors behind it are not conclusive yet. Traditional statistical analyses are inadequate to identify variants, genes, or pathways capable of explaining AD as a unit. In this context, pathway network analysis based on a set of curated AD-specific genes identified in the literature can elucidate biological mechanisms underneath AD. Through the network, we can infer influential pathways that can together explain AD. Consequently, we can target those pathways and corresponding genes for further analysis to develop new drugs, discover novel AD-related genes, combine multiple hypotheses, and so forth.

**Methods:** We have developed a novel graph theoretic algorithm that can elucidate complex biology from a given set of disease-related genes. It constructs a weighted network of enriched pathways where similarity score between a pair of pathways is defined in a context-specific manner. To make the network robust, we employ topological overlap techniques on top of the raw similarity measure. We then provide the importance of each pathway with respect to the entire network, functional modules and importance of each pathway in a specific module, gene clusters, and so forth. We also provide a method to identify a set of novel genes that can further explain the disease-related genes and the disease itself.

**Results:** We have employed our algorithms onto a set of AD-specific genes. It identified three distinct functional modules that are related to metabolism, cancer, and infectious disease related pathways. These findings are matched with three recognized hypotheses in Alzheimer’s disease, e.g. “metabolism hypothesis,” “cell cycle hypothesis,” and “infectious disease hypothesis.” By analyzing the curated genes common among those functional modules, we can attain more understanding about this fateful disease. We have also identified 24 novel AD-related genes of which at least 14 genes are known to be involved in AD.

**Conclusions:** We developed a computational framework for analyzing biological pathways in a context-specific manner. It can be used in any sets of disease-related genes. We manifest its efficacy, reliability, and accuracy by employing a set of AD-specific genes.

## Background

Alzheimer’s disease (AD) is the most common type of dementia leading to 60-70% of total cases of dementia [1]. There are two basic types of AD, e.g. early-onset (EOAD) and late-onset (LOAD). LOAD is the most common type commonly starting to appear after age 65. EOAD is a relatively rare and mostly diagnosed in individuals under the age of 65 (more frequent in their 40s and 50s). It is also known as young-onset AD and less than 5% of all AD patients have this type. Irrespective of the types, Alzheimer’s disease is a progressive neurodegenerative disease that gradually impairs memory and reasoning skills. Finally, it diminishes the ability of an individual to execute the simplest possible tasks. As bodily functions are lost due to the death of neurons, it ultimately leads to death [2]. Alzheimer’s Association enlists 10 early signs and symptoms of Alzheimer’s (https://www.alz.org/alzheimers-dementia/10_signs), such as memory loss that disrupts daily life, challenges in planning or solving problems, difficulty completing familiar tasks, confusion with time or place, trouble understanding visual images and spatial relationships, etc. Reportedly, around 5.8 million Americans were living with Alzheimer’s disease in 2020 and the number of individuals living with AD doubles every 5 years beyond age 65 [3]. According to this study, this number is estimated to approximately triple to 14 million people by the year of 2060. The total cost of care in the US for the treatment of AD in 2020 was estimated to be $305 billion and it is estimated to increase to more than $1 trillion by 2050 [4].

The cause of Alzheimer’s disease is not clearly explainable yet. There are plenty of environmental as well as genetic risk factors correlated with its advancement. The strongest genetic risk factor known is stemming from mutating an allele of APOE gene [5, 6]. Moreover, a history of head injury, clinical depression, and high blood pressure are also recognized as substantial risk factors associated with AD. The disease process is generally associated with amyloid plaques, neurofibrillary tangles, and loss of neuronal connections in the brain [2]. Among the stated factors above, it is likely that about 70% of the risk for AD is owed to genetics [7, 8]. But the genetics are different in AD patients based on its type. Scientists have identified mutations in three genes, namely Amyloid precursor protein (APP), Presenilin 1 (PSEN1), and Presenilin 2 (PSEN2) that are grossly responsible for early-onset Alzheimer’s disease [9]. If an individual inherits any one of these mutated genes from either of their parents, he will most likely have Alzheimer’s symptoms before the age of 65. On the contrary, the most common gene associated with late-onset Alzheimer’s disease is a risk gene called apolipoprotein E (APOE). Other notable genes include ATP Binding Cassette Subfamily A Member 7 (ABCA7), Clusterin (CLU), Complement C3b/C4b Receptor 1 (CR1), Phosphatidylinositol Binding Clathrin Assembly Protein (PICALM), Phospholipase D Family Member 3 (PLD3), Triggering Receptor Expressed On Myeloid Cells 2 (TREM2), Sortilin Related Receptor 1 (SORL1), etc. [10, 11, 12, 13]. The ongoing research suggests that a large number of genes that haven’t been identified yet might increase the risk of Alzheimer’s disease. It is a complex disorder and genes with various biological activities may together explain this fateful disease. But current form of genome-wide association studies (GWAS), which is being extensively applied to genetic studies of AD, is inadequate to accurately explain the genetics of complex traits due to the lack of sufficient statistical power. It explores each variant individually, but much of the heritability of some complex diseases like AD is missing because a person’s susceptibility to a particular disease may well depend on the combined effect of a set of variants. In this context, we propose a novel graph theoretic pathway network analytics algorithm that not only can decode the complex biology of AD but also can cater the opportunity to obtain critical understanding beyond the classical single-gene based analyses.

In graph theory, a graph/network is an abstraction of a structure containing a set of objects where the relation among the objects are well defined (i.e. they are “related” with respect to a specific context). The objects correspond to mathematical abstractions labelled as vertices (also known as nodes or points) and each of the related pairs of vertices is termed an edge (also dubbed as link or line) [14]. Complex biological systems may be represented and analyzed by constructing well-defined networks. Some notable examples of network in biology include protein-protein interaction networks, gene regulatory networks, metabolic networks, gene co-expression networks, among others. By analyzing the network, we can identify influential nodes, clusters having varying and distinct activity, outliers, and so on. Although network abstractions are prevalent in biology as stated above, there is a lack of pathway network analytics methods in the current literature. Moreover, existing methods are not only sketchy to decipher complex biology in the network but are also lacking automation [15, 16, 17]. A weak variation of the proposed algorithm can be found in [18]. It lacks robustness as it does not consider the neighboring pathways while estimating the weight of an edge. The centrality measure also does not account for a node’s influential neighbors while computing the importance scores. It also lacks scalability, i.e. its time complexity is exponential with the number of nodes in the network. To address all the challenges in developing a formal computational framework, we have come up with a novel graph theoretic method to analyze disease-related genes by constructing a weighted network of biological pathways. Weighted networks are extensively employed in genomic, and systems biologic applications [19], such as weighted gene co-expression network analysis (WGCNA) where the edge weight is defined to be the expression similarity (i.e. Pearson’s *r*) between the incident pair of genes [20]. Intuitively in weighted network, strength and weight information make better predictions about network structure than degree [21]. A theoretical advantage of weighted networks is that we can derive relationships among different network measures [19]. The most difficult task to build a weighted network is to define the (dis)similarity measure among the nodes in a context specific manner. In our proposed algorithm, we have defined the similarity among the pathways using their known and proven biological activity. Our method automatically extracts biological structures, such as functional modules in the network and their relevance, importance of each pathway, relevant gene clusters, etc. obscured in the complex network. In addition, we have also developed a novel network-centric method for identifying a set of novel genes based on the given set of disease-related genes.

### Methods

In this article we propose two graph-theoretic methods, e.g. (1) a novel weighted pathway network analytics algorithm and (2) a method to discover novel disease-related genes based on a weighted protein-protein interactions network. Next, we illustrate each method in detail.

### Weighted biological pathway network analytics

At the beginning, we identify a set of enriched biological pathways. These enriched pathways are then used to construct a weighted network of pathways. It runs in 4 stages. Next, we provide each step in details. Pseudocode of the proposed algorithm can be found in Algorithm 1.

### Identification of disease-specific enriched pathways

Kyoto Encyclopedia of Genes and Genomes (KEGG) pathways were utilized to elucidate the biological theme of a set of AD-related genes. It is a collection of databases dealing with genomes, biological pathways, diseases, drugs, and chemical substances. We employed Hypergeometric overrepresentaion test that uses the Hypergeometric distribution to calculate the statistical significance of a KEGG pathway with respect to the AD-related genes. Specifically, we computed a hypergeometric *p*-value for each of the KEGG pathway to assess whether a pathway is overrepresented with AD-related genes. In this study, pathways having Bonferroni corrected hypergeometric *p*-value smaller than 0.05 were retained for further analyses.

### Construction of a weighted network

After identifying the enriched pathways, to mirror true biological synergy, we construct an undirected weighted network of pathways. In our network, each pathway acts as a node and two nodes are connected by an edge *iff* they have some explicit and intuitive biological similarity. We define the similarity in such a way so that it strongly associates their shared biological activity and the range of numerical values it can have must be lied within 0.0 to 1.0. Let *G*(*V, E, W*_*E*_) be our undirected weighted graph where each enriched pathway *v* ∈ *V* operates as a vertex. Two vertices *v*_*i*_ and *v*_*j*_ will be connected by an edge *e* ∈ *E iff v*_*i*_ and *v*_*j*_ share at least *x* common genes in our experiment, *x* was set to 4. The functional similarity between any two genes can roughly be estimated by their common gene ontology-biological process (GO-BP) terms with a high level of accuracy. Consequently, two pathways will be functionally similar if they contain a set of functionally related genes between them. This observation in turn can be generalized by considering only the common GO-BP terms between any of the two pathways. By considering this fact stated above, we compute the Jaccard index of GO-BP terms between any two pathways. Suppose *v*_*i*_ and *v*_*j*_ pathways consist of *G*_*i*_ and *G*_*j*_ sets of genes. We extract all the GO-BP terms from the *G*_*i*_ and *G*_*j*_ sets of genes. Let *T*_*i*_ and *T*_*j*_ be sets of GO-BP terms corresponding to *G*_*i*_ and *G*_*j*_ sets of genes, respectively. To measure the standard similarity score between any pairs of pathways *v*_*i*_ and *v*_*j*_, the overlap index (*OI*_*ij*_) and the Jaccard index (*JI*_*iz*_) were calculated using the corresponding formulas:

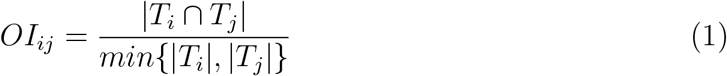

and

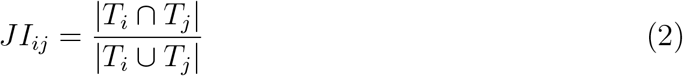

Finally, the similarity score between any two vertices *v*_*i*_ and *v*_*j*_ can then be defined as the mean of *OI*_*ij*_ and *JI*_*ij*_ as follows:

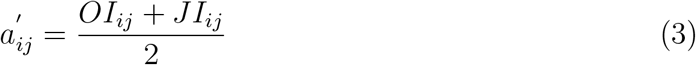

As a result, the minimum and maximum value of such a score will be 0.0 and 1.0, respectively. Intuitively, the higher the score, the more will be two pathways functionally similar.

An important aim of the proposed method is to identify subsets of nodes (i.e. modules) that are “tightly” connected to each other. Apart from just defining the similarity measure in this article, we use the topological overlap measure between the pair of pathways. Since, the topological overlap of two nodes mimics their relative association (likely non-spurious), it assists to accurately identify biologically relevant and meaningful modules [22, 23]. For unweighted networks, the topological overlap score *w*_*i,j*_ between the pair *v*_*i*_ and *v*_*j*_ in the network *G* is formulated as:

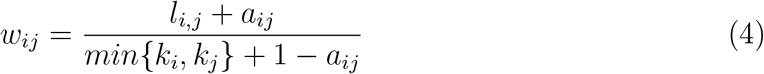

where *l*_*ij*_ = Σ_*u*_ *a*_*iu*_*a*_*uj*_, and *k*_*i*_ = Σ_*u*_ *a*_*iu*_ is the node connectivity. Consequently, *l*_*ij*_ equals the number of nodes to which both *v*_*i*_ and *v*_*j*_ are connected. It is to be noted that *w*_*ij*_ = 1 if *v*_*i*_ and *v*_*j*_ are connected and they have the identical set of neighbors. On the other hand, *w*_*ij*_ = 0 if *v*_*i*_ and *v*_*j*_ are not connected and they do not have any shared neighbors. According to [20], we generalize the topological overlap score for weighted networks by simply replacing *a*_*ij*_ with 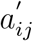 found from Equation 3. Since, 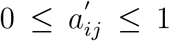, it satisfies 0 ≤ *w*_*ij*_ ≤ 1 constraint in weighted undirected graph. Now, the topological overlap score *w*_*ij*_ constitutes the weight of the edge between the two pathways *v*_*i*_ and *v*_*j*_ in the weighted network *G*.

### Identification of important pathways

In graph theory, node centrality measure has a diverse set of practical applications in a wide area of fields, such as but not limited to identifying the most influential person(s) in a social network, key infrastructure nodes in the Internet or urban networks, superspreaders of a disease, and brain networks [24, 25]. A set of graph centrality measures has been defined so far, such as degree centrality, closeness centrality, betweeness centrality, eigenvector centrality, etc. Historically first and conceptually simplest is degree centrality. It is defined as the number of edges incident upon a node. In an undirected graph, degree centrality awards one centrality point for each of the incident edges of a node. But not all nodes in a weighted undirected graph are equivalent in terms of its influence; some are more influential than others. Consequently, affirmations from influential nodes should count more. The eigenvector centrality [26] exactly considers this observation: *A central node is connected to other central nodes*. A high eigenvector score means that a node is connected to many nodes who themselves have high scores [27, 28]. For a given graph *G* = (*V, E, W*_*E*_) where *V* is set of nodes involved in the network. *E* ⊆ *V* × *V* is the set of edges in the graph and *W*_*E*_ is the set of weights. For an example *w*_*ij*_ ∈ *W*_*E*_ is the weight of an edge *e*_*ij*_ ∈ *E*. let *A* = (*a*_*i,j*_) be the adjacency matrix, i.e. *a*_*i,j*_ = *w*_*ij*_ if vertex *v*_*i*_ is connected with vertex *v*_*j*_ where their edge weight is *w*_*ij*_, and *a*_*i,j*_ = 0 otherwise. The relative centrality score of vertex *v* can be defined as:

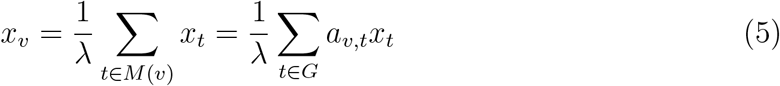

where *M* (*v*) is a set of the neighbors of *v* and *λ* is a constant. It can be rewritten in vector notation as the eigenvector equation **Ax** = *λ***x**. Usually, we can end up with a set of different eigenvalues *λ* each having a non-zero eigenvector solution. However, according to the Perron–Frobenius theorem, eigenvector having the greatest eigenvalue results in the desired centrality measure. The *v*^*th*^ component of the related eigenvector then gives the relative centrality score of the vertex *v* in the network.

### Identification of functional modules and assignment of importance

One of the primary objectives of the proposed method is to disassemble the weighted network of pathways into several clusters in such a way so that each cluster of pathways represents an explicit biological event/theme. Louvain method is an attractive and suitable choice in this context [29]. It is a greedy optimization method having time complexity 𝒪 (*n · log*^2^*n*) where *n* is the number of nodes in the network. It maximizes a modularity score for each group, where the modularity quantifies the quality of an assignment of nodes to groups. For a weighted graph, modularity is defined as:

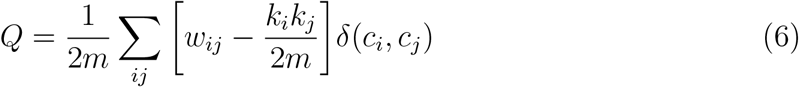

In the above formula, *w*_*ij*_ is the edge weight between nodes *v*_*i*_ and *v*_*j*_, *k*_*i*_ is the degree of node *v*_*i*_, *m* is the sum of all the edge weights in the graph, and *δ* is Kronecker delta function. Louvain algorithm runs in 2 stages. At first, small groups are identified by optimizing modularity locally on all nodes. Finally, each small group is merged into one node and the above procedure is repeated until maximum modularity is attained.

### Discovery of novel disease-specific genes

We have also identified a set of novel disease-related genes based on the given set of curated disease-related genes. We have also measured the centrality score of each novel disease-related gene by cleverly constructing a weighted network. Next, we provide each step in details.

### Disease-specific network of protein-protein interactions

After analyzing the disease-related genes *G*, the next logical step is to predict a set of novel genes by utilizing *G*. Steiner tree is the natural choice in this context (https://en.wikipedia.org/wiki/Steiner_tree_problem). The problem is defined as follows: given an undirected graph with non-negative edge weights and a set of vertices, the Steiner Tree in graph is a minimum spanning tree (say *M*) having minimum weight (i.e. sum of edge weights in *M* is as small as possible). The constraint is that it contains a subset of given nodes (usually a set of related genes, such as disease-specific genes) known as terminals. It may consist of additional nodes to construct Steiner tree *M* termed as

#### Algorithm 1

Elucidating Network of Biological Pathways

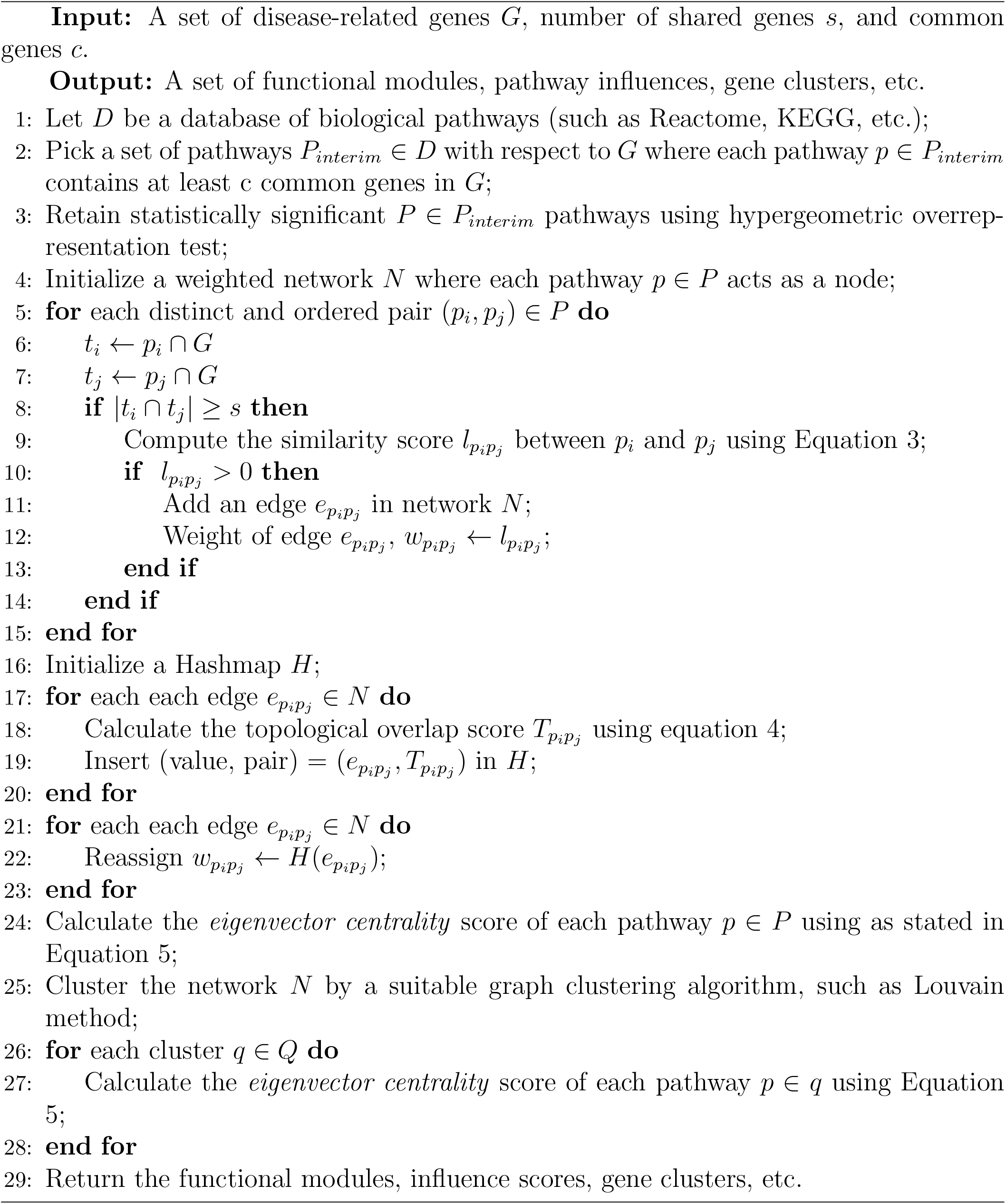

Steiner nodes. In our study terminal nodes constitute subset of our disease-related genes *G*. Consequently, the Steiner nodes can be recognized as a set of critical candidates potentially can further explain the relationship among the genes in *G*. Widely, Steiner nodes correspond to proteins or genes that are candidates for the regulation of the given set of disease-related genes as the Steiner nodes connect them in the network in a compact way.

Finding an optimal minimum weight Steiner tree is NP-complete problem, i.e. there is no known polynomial time algorithm exists [30]. A set of approximation algorithms exists to build approximate Steiner tree based on a varying set of heuristics in the current literature, such as shortest paths-based approximation (SP), minimum spanning tree based approximation (KB), randomized all shortest paths approximation (RSP) [31]. As suggested in [31], we have used SP [32] variant in constructing approximate Steiner tree. At first, we construct a protein-protein interaction network by utilizing PrePPI (Predicting Protein-Protein Interactions by Combining Structural and non-Structural Information) [33, 34]. In our network, each protein is encoded as a node and two nodes are connected via an edge if-and-only-if PrePPI predicted score between them is *>* 0.5. Note that PrePPI predicted score ranges from 0.0 to 1.0. After constructing the network, we extract an approximate minimum weight Steiner tree where the terminals constitute our curated list of genes and Steiner nodes are the predicted disease-specific genes.

### Construction of a weighted network and identification of influential genes

After forming the minimum Steiner tree, we associate a weight for each of the edges in the tree. The set of neighbors of a protein *u* is defined to be all other proteins in the protein-protein interaction network that are directly connected with protein *u*. Let *N* (*u*) and *N* (*v*) be the set of neighbors of *u* and *v*, respectively. The weight of edge *e*(*u, v*) is defined as the Jaccard index of *N* (*u*) and *N* (*v*):

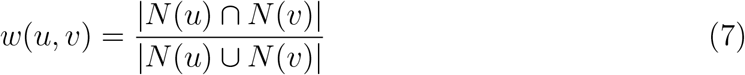

After setting weight to each edge in the Steiner tree *T*, we assign scores to each of the nodes using eigenvector centrality measure as described earlier.

## Results

To demonstrate the efficacy of our algorithms, we have performed rigorous experimental evaluations with respect to a set of curated AD-specific genes found in the existing literature. Please, note that enrichment of gene ontology (GO) and disease ontology (DO) terms were analyzed by clusterProfiler [35]. Next, we illustrate the findings in detail.

### Dataset

We curated a set of AD-specific genes from numerous sources, e.g. (1) genes annotated with Alzheimer’s disease term (DOID:10652) in Disease Ontology (DO) database (https://disease-ontology.org), (2) AD-related genes curated through literature mining by Hu et *al*. [17], (3) AD-specific genes curated by DisGeNET discovery platform by employing BeFree system [36], (4) Comparative Toxicogenomics Database (CTD) (http://ctdbase.org/), (5) ClinVar (https://www.ncbi.nlm.nih.gov/clinvar/), etc. Finally, we took 242 AD-related genes found at least 50% time across the different databases for our analyses. Throughout this article, we call this set of 242 genes as *G*.

### Biological pathway enrichment analysis

We found 45 Bonferroni corrected (adjusted *p <* 0.05) enriched biological pathways from KEGG dataase (https://www.genome.jp/kegg/) as shown in Table 1. Please, note that *p*-values are calculated based on Hypergeometric over-representation test (https://en.wikipedia.org/wiki/Hypergeometric_distribution). It is to be noted that we only retain those enriched pathways each having at least eight genes in common with our AD-related genes *G* to discard potentially spurious and smaller pathways. These pathways can be grouped into several functional clusters based on their biological activities. More details are discussed in the subsequent sections.

**Table 1:**
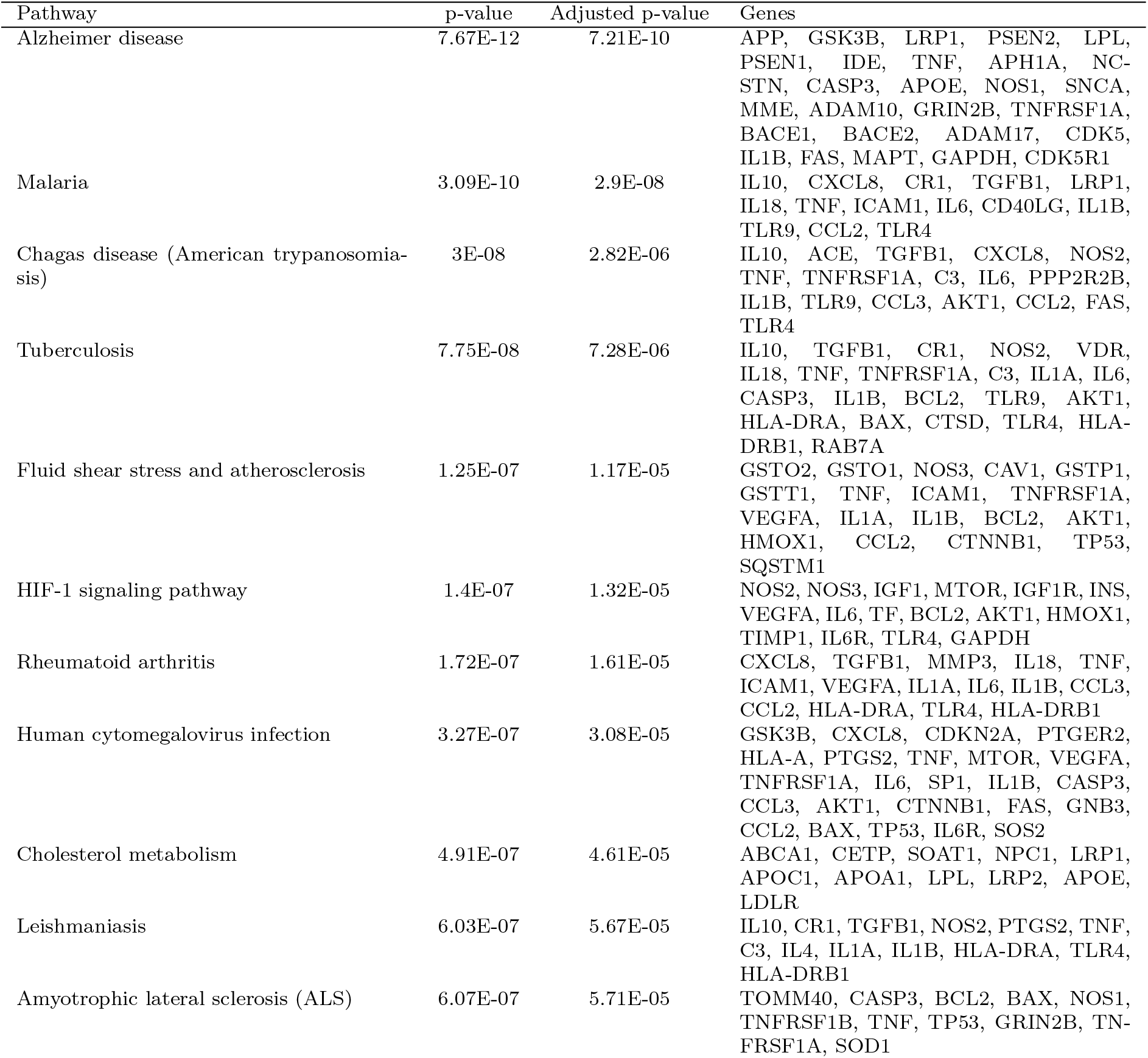
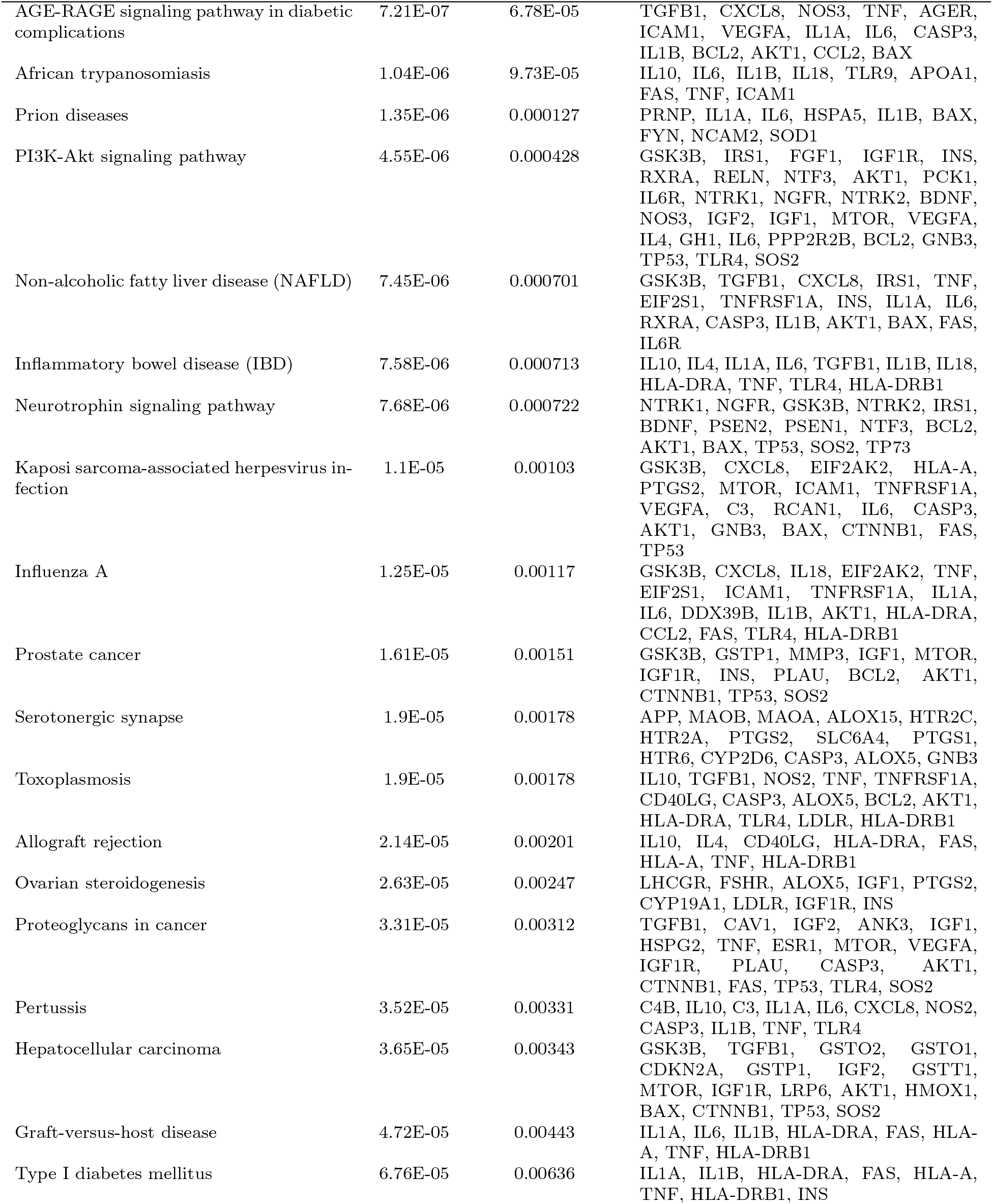
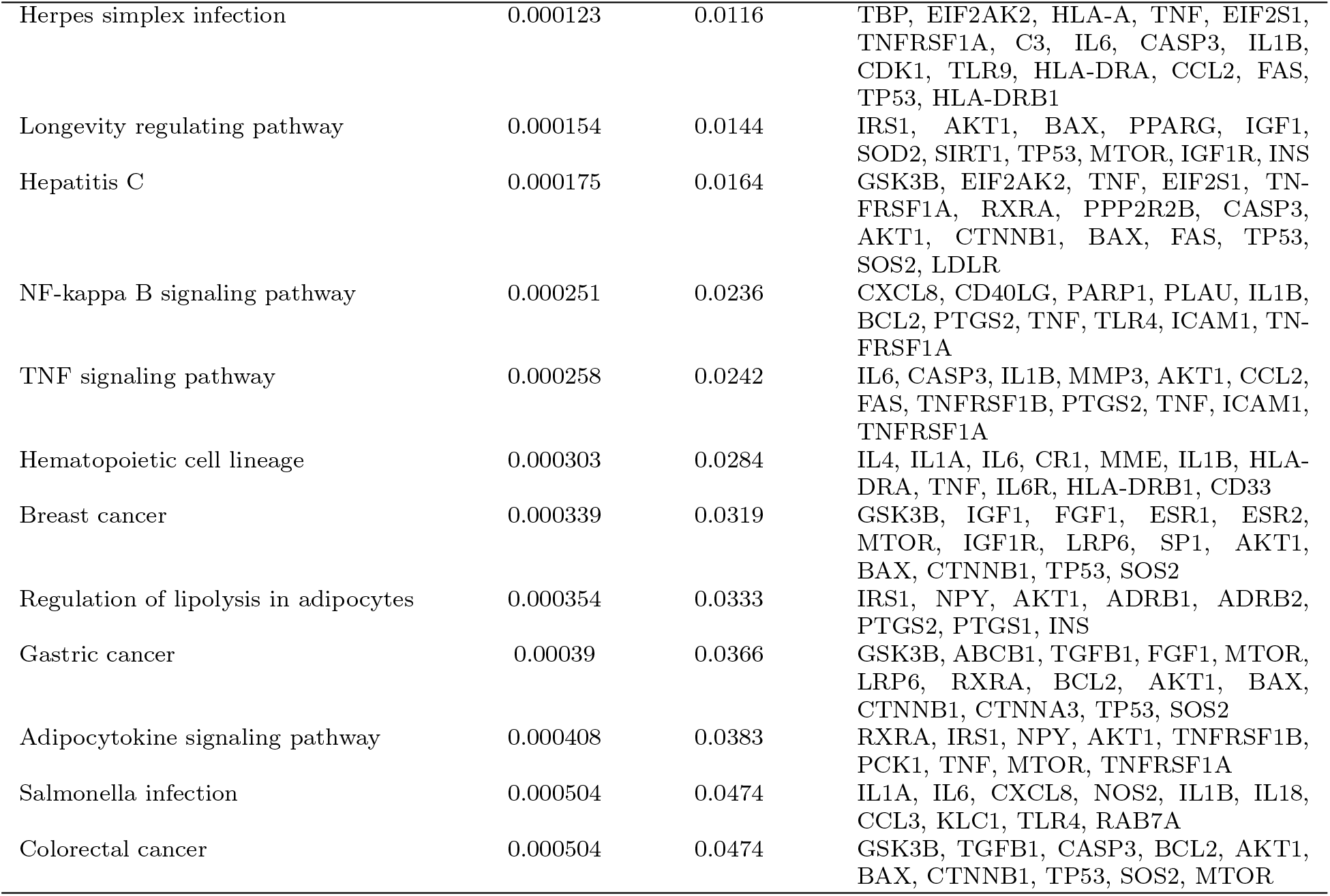
Enriched (Bonferroni corrected *p <* 0.05) pathways based on 242 curated list of AD-specific genes.

### Pathway network analysis

After finding the statistically significant pathways, we build a network from those pathways as stated in Methods section. Please, note that in our experiment any 2 nodes are connected in the network by an edge if they have 4 genes in common. Given this scenario, our network contains 41 pathways and 408 edges. In addition, we have 126 genes (out of 242 genes in *G*) harboring in those 41 pathways.

### Important pathways

As noted earlier, after constructing the weighted network, we compute the influence of each pathway based on *eigenvector* centrality. The corresponding network is shown in Figure 1. The importance of each pathway with respect to the entire network is proportional to the diameter of its representative circle and the width of each edge is proportional to its weight. Figure 2 illustrates the *eigenvector* centrality scores of each of the 41 KEGG pathways in the network. It shows that “Non-alcoholic fatty liver disease (NAFLD)” and “prion diseases” is the most important and least important pathway, respectively in the entire network.

**Figure 1:**
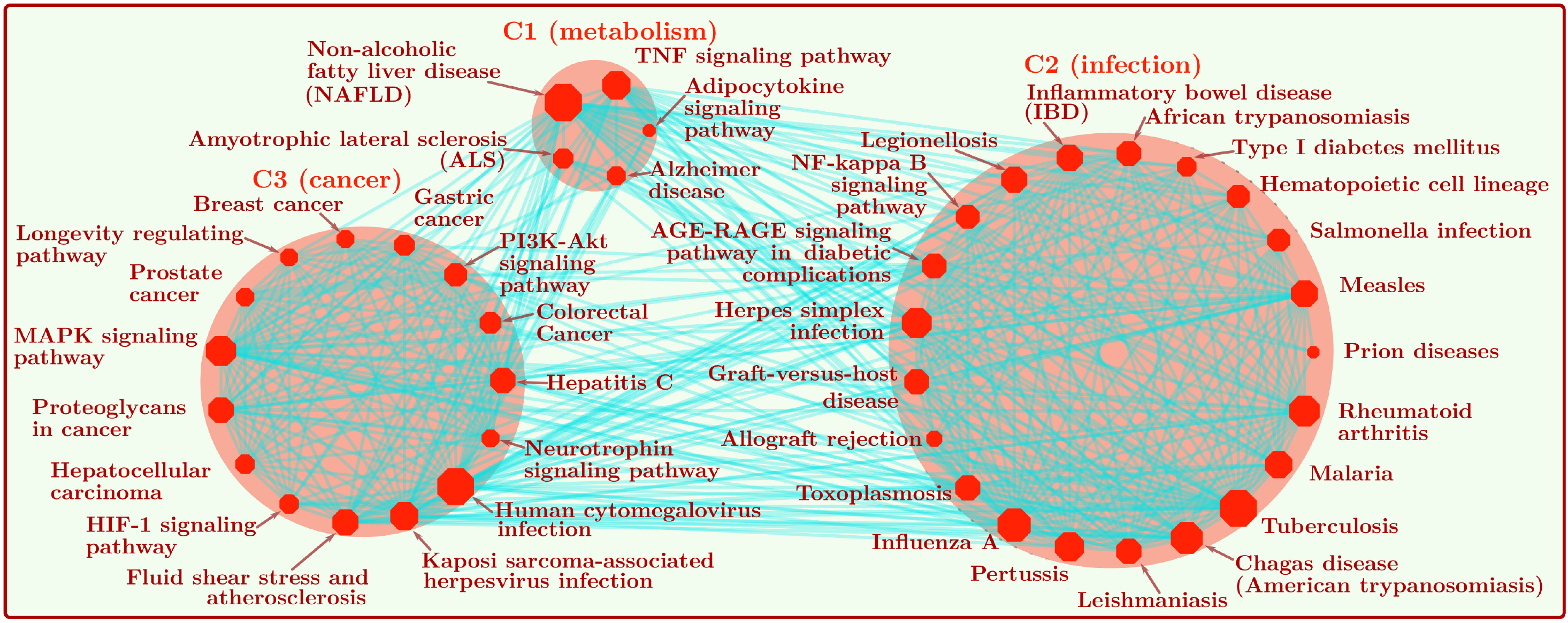
Entire network built from 41 statistically significant (Bonferroni corrected *p <* 0.05) KEGG pathways.

**Figure 2:**
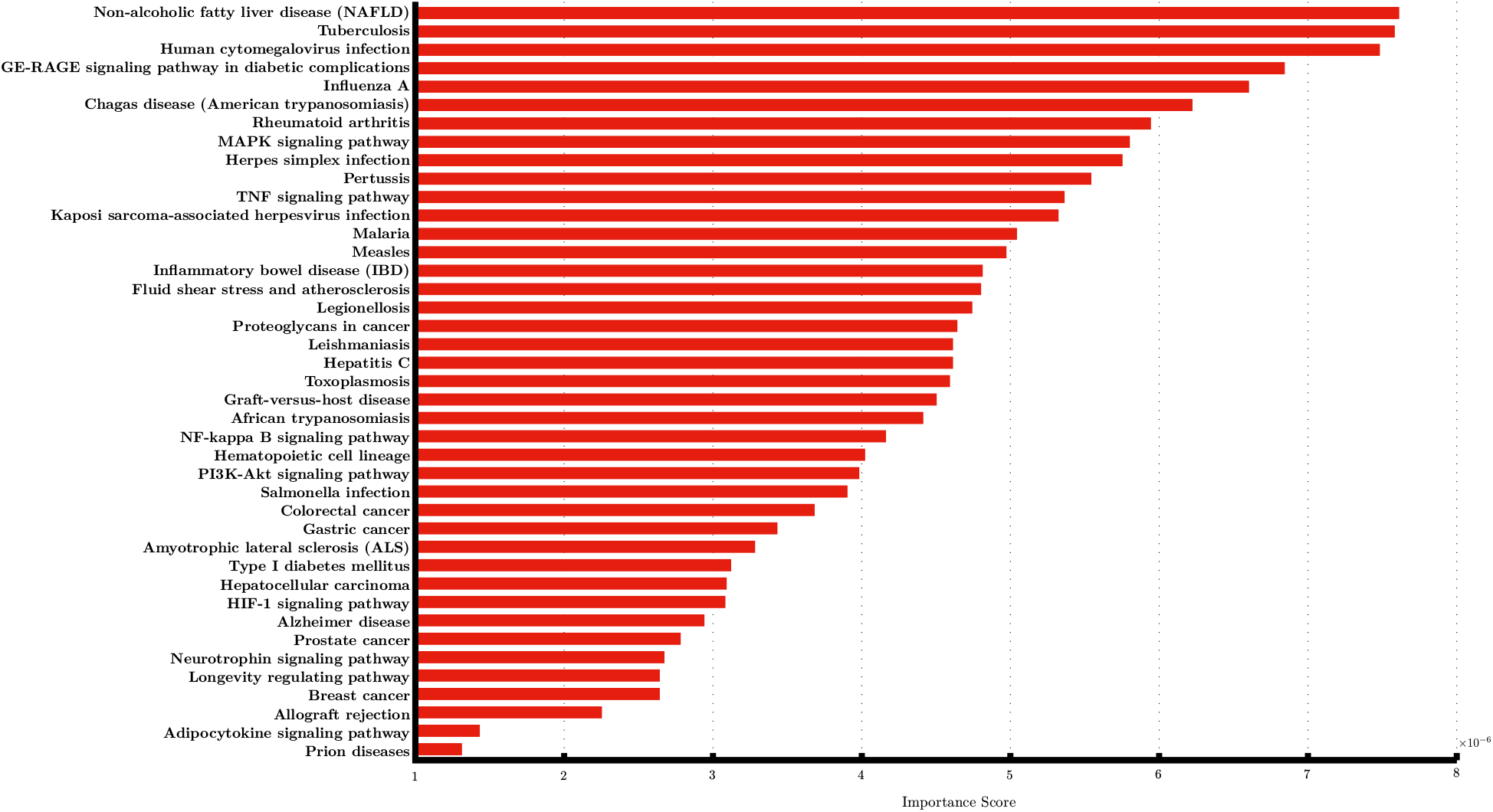
Importance scores for 41 biological pathways in the entire network.

### Functional modules

After constructing the weighted network, we cluster the network to identify functional modules. Since, the weight of an edge corresponds to the functional similarity between two pathways, each of the modules consisting of highly interconnected pathways should mimic a specific biological theme or functionality. Note that we do not impose any kind of thresholds on the weights to discard any edges. Moreover, we do not need to provide the number of clusters *a priory*. The clustering algorithm automatically dismantles the entire weighted network into three groups, namely C1, C2, and C3. Figure 1 shows the entire network along with cluster annotations. C2 and C3 functional modules essentially represent infectious disease and cancer related pathways, respectively. C1 functional module is interesting, and it seems act as a buffer or interface between C2 and C3 functional modules. It broadly represents metabolism related pathways.

#### C2: Inflammation and infectious disease related module

At first, consider the C2 functional module. It consists of 21 pathways where all most all of them are inflammation and infectious disease related pathways. This finding at first seems baffling. Pathologically, AD usually associates with the accumulation of two types of proteins in our brain, namely *amyloid* and *tau*. These earliest signs are so prevalent in AD patients, that the leading hypothesis on the progression of AD pathologies states that the disease is caused by the degenerative control of these proteins where amyloid first accumulates to form large, sticky plaques around brain cells. Eventually, the formation of plaques is responsible for hyperphosphorylation of *tau* which in turn ultimately leads to tangles and neurodegeneration. This hypothesis known as “amyloid hypothesis” is the pivotal actor into understanding and treating Alzheimer’s disease. Most of research were conducted based on this hypothesis and billions of dollars have been invested in the experiments without any breakthroughs. According to a study [37], the failure rate of drug development for Alzheimer’s has been 99%. Consequently, failure of the so called “amyloid hypothesis” encourages to formulate alternative hypotheses. One of the rising but controversial hypotheses among all is the “infectious hypothesis.”

According to the “infectious hypothesis,” a pathogen such as virus, bacteria, prion, etc. is the leading cause of AD [38, 39, 40]. In the existing literature, multiple microbial organisms have been reported that can have potential to trigger Alzheimer’s disease [41]. It largely includes three human herpes viruses and three bacteria, e.g. *Chlamydia pneumoniae, Borrelia burgdorferi*, and *Porphyromonas gingivalis*. Three types of disease are associated with those three bacteria, e.g. lung infections, Lyme disease, and gum disease, respectively. In theory, any infectious microbes that can penetrate blood-brain barrier (BBB) and so, can invade the brain might have the ability to trigger AD. Indication from current research is thriving that infection can spark the build-up of sticky protein plaques in the brain. According to the “amyloid hypothesis”, it is the hallmark of Alzheimer’s disease. Several theories exist in the current literature to elucidate the biological mechanisms behind it. One theory is particularly intuitive and next, we briefly discuss about it. The theory states that microbial organisms irritate a particular type of brain cells known as microglia. It sets an immune reaction and hikes the level of an enzyme that bolsters the production of amyloid proteins and gradually, it aggregates the plaques in the brain. Specifically, amyloid is thought to be acting as a protector by refraining microbes from brain cells. However, the failure of biological mechanisms to clear up the amyloid from the brain cause the inflammation. All these events essentially create a toxic feedback loop. In this context, “amyloid hypothesis” and “infectious hypothesis” can be fitted in one grand hypothesis. An excellent review on different hypothesis for Alzheimer’s developed over time can be found in [42].

#### C3: Cancer related module

Now, consider the C1 functional module. It consists of 14 cancer related pathways. This finding supports the published data [43, 44, 45, 46]. An encyclopedic longitudinal study conducted on more than one million participants indicated an inverse association between AD and cancer. In this study, the risk of cancer in the presence of AD was scaled down to 50% and the risk of AD in individuals with cancer was reduced by 35% [47]. In an overwhelming number of publications it is found that many factors that are upregulated in cancer to sustain growth and survival are downregulated in Alzheimer’s disease contributing to neurodegeneration and death of neurons [45]. As stated earlier, “amyloid hypothesis”, also known as “beta-amyloid (A*β*) cascade hypothesis,” suggests that it is the A*β* deposition that first triggers the AD pathogenesis, spawning the other AD hallmarks, such as NFTs, neuro-inflammation, synaptic loss, and neurodegeneration [48, 49]. According to “Cell cycle hypothesis,” abnormal re-entry into the cell cycle is believed to trigger the pathway resulting in NFTs, apoptotic avoidance, and A*β* production [50, 51, 52]. This is where (i.e. cell cycle activation), AD and cancer have common pathological phenomenon. Cancer is characterized by everlasting and abnormal repeat of cell cycle where the neurons in AD brains also show the proliferated activation of the cell cycles without completing it resulting in death of neurons [51].

#### C1: Metabolism related module

Finally, consider the C1 functional module. It consists of 5 pathways largely related to metabolism [53, 54, 55]. This finding is aligned with the published data [56, 57]. Oxidative stress is known to be induced by A*β* and neurofibrillary tangles (NFTs). On the contrary, oxidative stress caused by metabolic alterations in the body induce brain metabolic alterations, resulting in AD. This fact is also known as “metabolism hypothesis.” For example, according to [56], individuals suffering from metabolic diseases show higher levels of oxidative stress that alters brain metabolism and ultimately, it leads to progressive Alzheimer’s disease. As stated earlier, oxidative stress is widely shown in AD brains [58] and along with metabolic stress, they can be considered as the central actors to initiate cell cycle and developing AD and cancer pathology [59, 60].

### Gene clusters

Our network analytic method also provides relevant gene clusters from the set of genes in *G*. Each gene cluster is associated with each of the functional modules. The observation is that if a functional module is distinct in terms of its biological activities, the genes harboring in that module should also perform specific biological functions. Since, we get 3 distinctive functional modules from our network, we have 3 gene clusters as shown in Figure 3. It shows that we have 126 genes (out of 242 genes in *G*) harboring in those 3 gene clusters. Although each of our functional modules performs distinct biological functions, we have found statistically significant overlap between each pair of gene clusters. Note that 21 genes are common among the 3 gene clusters. Statistically significant overlaps among the gene clusters coming from distinct functional modules suggest that it might be possible to formulate a unified hypothesis by combining all three hypotheses stated above.

**Figure 3:**
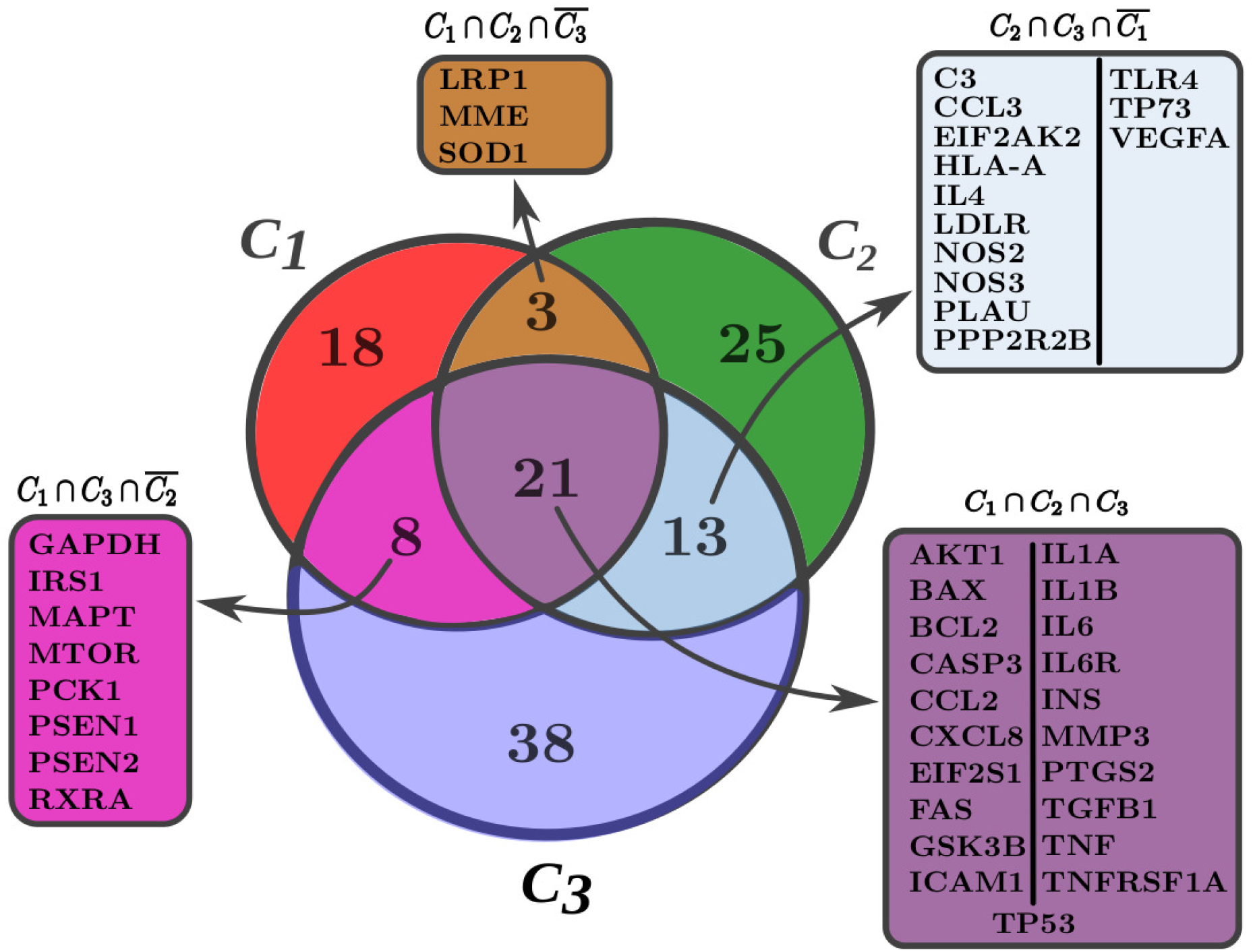
Gene clusters based on entire network of pathways.

### Few statistical analyses

We have performed a few biological enrichment analyses by considering genes in *G*. We illustrate each of them below.

### Disease Ontology (DO) term enrichment analysis

As stated above we performed disease ontology (DO) term analysis by considering genes in *G* (https://disease-ontology.org [61]). We found 462 Bonferroni corrected (adjusted *p <* 0.05) DO terms. Figure 4 illustrates top 20 such terms. All most all of them are linked to Alzheimer’s disease according to the current literature. We next illustrate few of the interesting ontology terms in brief.

**Figure 4:**
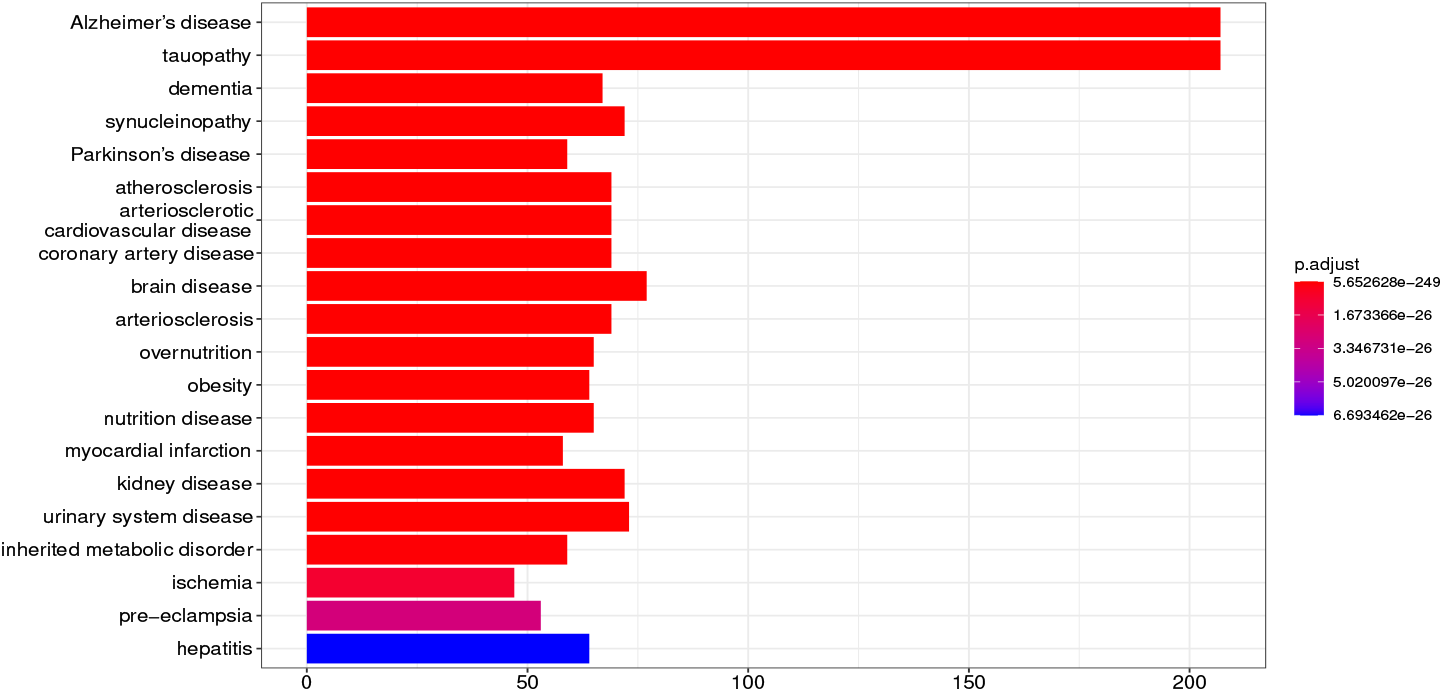
Top 20 enriched (Bonferroni corrected *p <* 0.05) DO terms based on curated list of 242 AD-related genes.

At first, consider preeclampsia disease ontology term. It is a serious blood pressure condition that can happen after the 20^*th*^ week of pregnancy or after giving birth known as postpartum preeclampsia. There is a well-documented relation between preeclampsia and cognitive impairment later in life [62]. According to a recent research on large cohort consisting of 1,178,005 women with 20,352,695 person years of follow-up, preeclampsia is associated with an increased risk of dementia, particularly vascular dementia [63]. Contrarily, only modest associations were observed for Alzheimer’s disease (hazard ratio 1.45, 1.05 to 1.99) in this study. As age is the biggest risk factor for developing dementia and most of the women participating in the study were young, more years of study would help better understanding the association between preeclampsia and Alzheimer’s.

Cardiovascular disease, such as arteriosclerotic cardiovascular disease, coronary artery disease as well as cardiovascular risk factors, such as atherosclerosis, arteriolosclerosis, obesity, diet are strongly and likely causally associated with dementia and Alzheimer’s later in life. According to the “vascular hypothesis,” AD is a vascular disease [64, 65]. It states that the pathology behind AD is induced via microvascular abnormalities. Moreover, it is a well-known fact that kidney function is connected with brain activity. In clinical studies, persons suffering from chronic kidney disease (CKD) were found to be susceptible to cognitive decline and AD [66, 67].

### Gene Ontology (GO) term enrichment analysis

We performed gene ontology - biological process (GO-BP) (http://geneontology.org) term enrichment analysis based on the genes in *G*. We retained only enriched terms having Bonferroni corrected *p <* 0.05. We found 874 enriched GO-BP terms and top 20 such terms are displayed in Figure 5.

**Figure 5:**
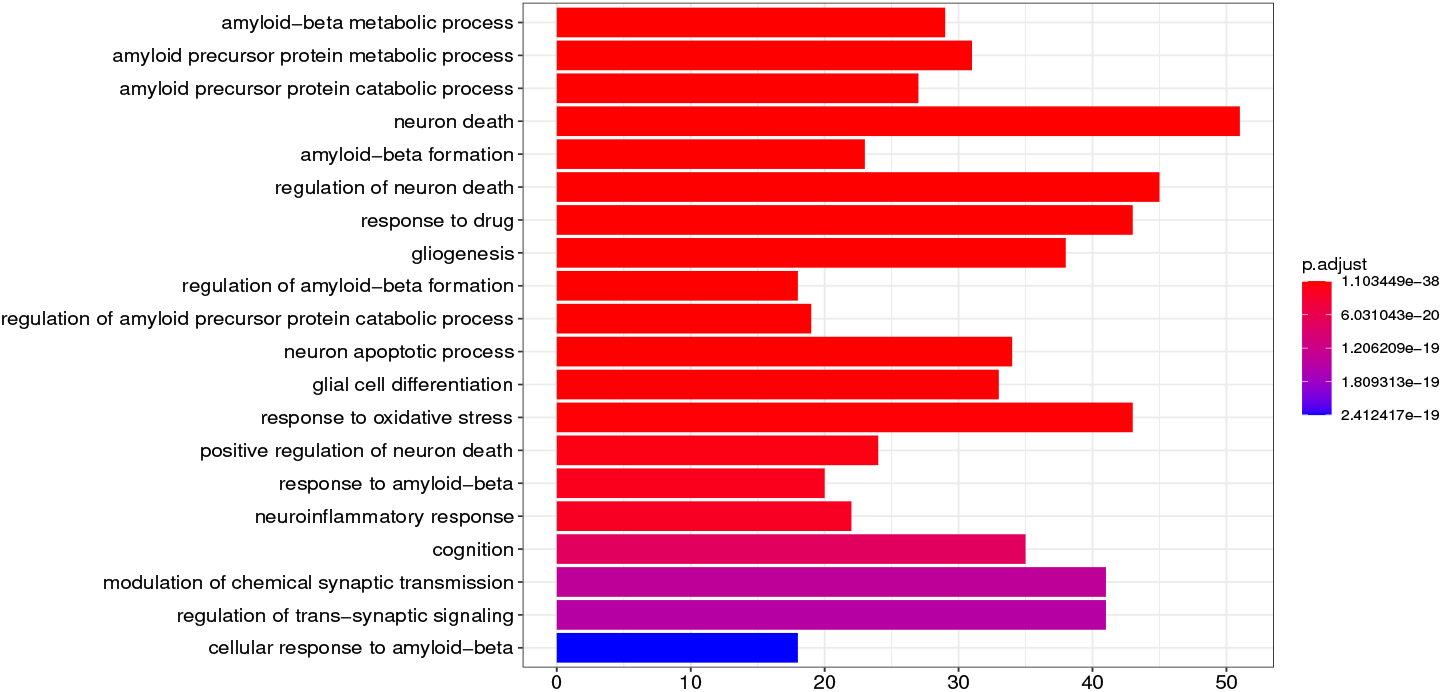
Top 20 enriched (Bonferroni corrected *p <* 0.05) GO-BP terms based on curated list of 242 AD-related genes.

### Novel AD-specific genes

We have found 24 novel AD-specific genes (say *G*_*N*_) based on our curated list of 242 genes *G* as shown in Table 2. The genes are ordered based on their importance in the Steiner tree as shown in Figure 6. Please, see Methods sections for details about the procedure. Out of 24 novel genes, at least 14 genes are involved in Alzheimer’s disease in the current literature (e.g. RB1, ABL1, EP300, JAK2, SHC3, RACK1, PPARD, EGFR, YWHAZ, CDK2, GRB2, CK2, MAPK1, and CUL3). Moreover, many of them are thought to be novel therapeutic drug target for AD. We will briefly discuss about a few novel genes next.

**Table 2:** Novel 24 AD-related genes identified in Steiner tree. The genes are ordered based on their importance.

| Symbol | Gene name | Entrez ID |
| --- | --- | --- |
| RB1 | RB transcriptional corepressor 1 | 5925 |
| SHC3 | SHC adaptor protein 3 | 53358 |
| EP300 | E1A binding protein p300 | 2033 |
| JAK2 | Janus kinase 2 | 3717 |
| VAV1 | vav guanine nucleotide exchange factor 1 | 7409 |
| YWHAB | tyrosine 3-monooxygenase/tryptophan 5-monooxygenase activation protein beta | 7529 |
| CUL3 | cullin 3 | 8452 |
| MAPK1 | mitogen-activated protein kinase 1 | 5594 |
| GRB2 | growth factor receptor bound protein 2 | 2885 |
| CDK2 | cyclin dependent kinase 2 | 1017 |
| YWHAQ | tyrosine 3-monooxygenase/tryptophan 5-monooxygenase activation protein theta | 10971 |
| EGFR | epidermal growth factor receptor | 1956 |
| ABL1 | ABL proto-oncogene 1, non-receptor tyrosine kinase | 25 |
| YWHAZ | tyrosine 3-monooxygenase/tryptophan 5-monooxygenase activation protein zeta | 7534 |
| CSNK2A1 | casein kinase 2 alpha 1 | 1457 |
| TRIO | trio Rho guanine nucleotide exchange factor | 7204 |
| SGSM3 | small G protein signaling modulator 3 | 27352 |
| PPARD | peroxisome proliferator activated receptor delta | 5467 |
| ARHGEF12 | Rho guanine nucleotide exchange factor 12 | 23365 |
| RAB1A | RAB1A, member RAS oncogene family | 5861 |
| FUT8 | fucosyltransferase 8 | 2530 |
| LCP2 | lymphocyte cytosolic protein 2 | 3937 |
| SPATA13 | spermatogenesis associated 13 | 221178 |
| RACK1 | receptor for activated C kinase 1 | 10399 |

**Figure 6:**
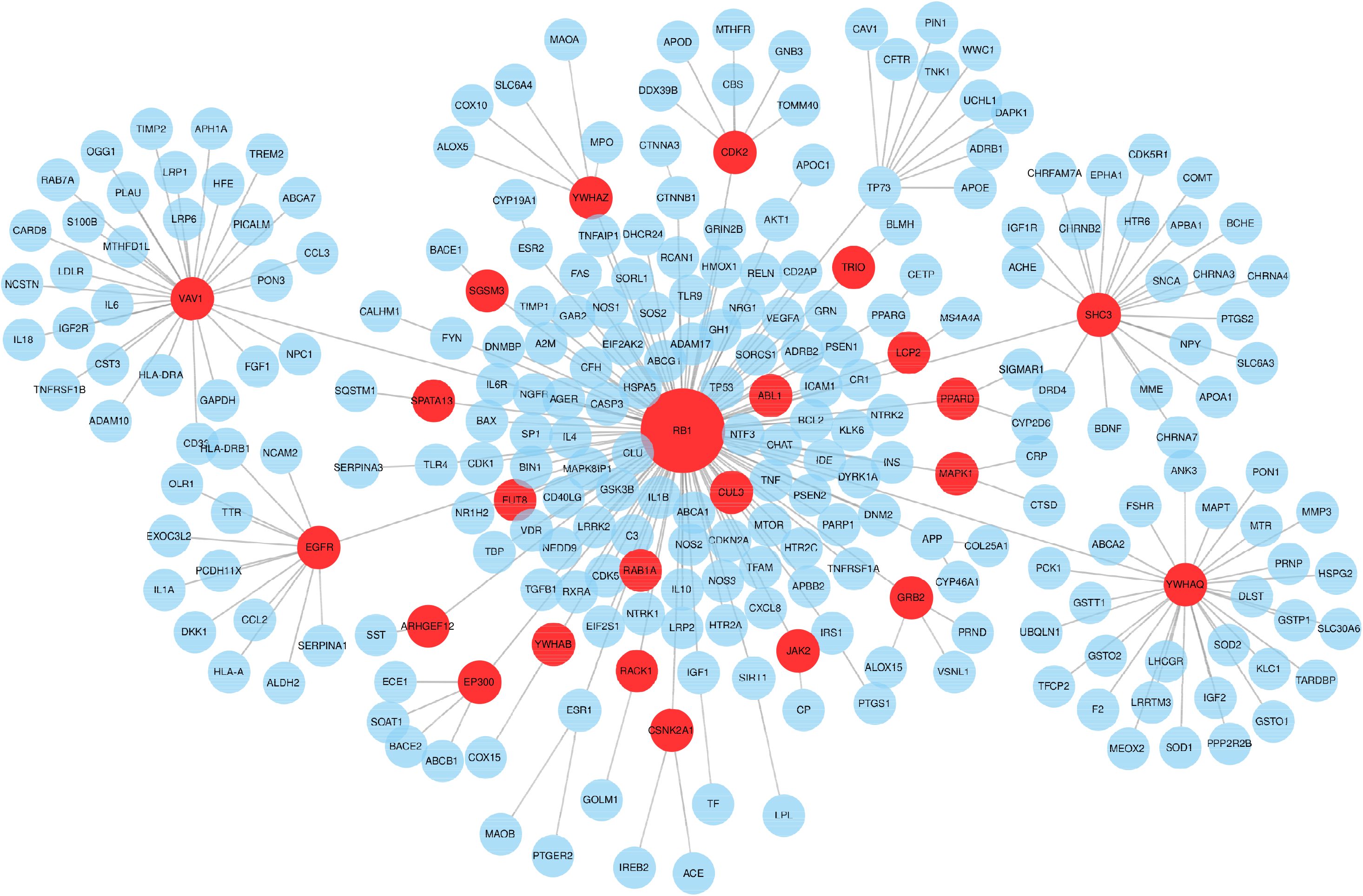
24 novel genes shown as red marked circle in the Steiner tree. Diameter of the circle is proportional to its importance.

### PPARD (peroxisome proliferator activated receptor delta)

PPARD is a nuclear hormone receptor. As a transcription factor, it regulates numerous biological processes and is thought to be involved in the development and progression of several chronic diseases, such as diabetes, obesity, atherosclerosis, and cancer [68, 69]. According to [70], PPARD agonism leads to reduction in brain A*β* levels in 5XFAD mice and it suggests that PPARD agonists may have therapeutic utility in treating AD.

#### EP300 (E1A binding protein p300)

It functions as histone acetyltransferase that regulates transcription of genes by means of chromatin remodeling. Therefore, it is critical in cell proliferation and differentiation. Epigenetic modifications, specifically, histone acetylation, is involved in Alzheimer’s disease [71]. EP300, also known as histone acetyltransferase p300, govern the expression of AD-related genes through regulating the acetylation of their promoter regions. It indicates that p300 may serve as a novel and potential therapeutic target for AD [72].

#### ABL1 (ABL proto-oncogene 1, non-receptor tyrosine kinase)

ABL1, a proto-oncogene, also known as c-ABL, encodes a cytoplasmic and nuclear protein tyrosine kinase involved in cell growth, cell division, cell adhesion, stress response, and the pathogenesis of chronic myeloid leukemia (CML) (https://www.uniprot.org/uniprot/P00519). According to several studies, c-Abl gene could be a critical actor in the early stages of AD and also has the potential to lead to new therapies for treating Alzheimer’s and other neurodegenerative diseases [73, 74, 75].

#### RACK1 (Receptor for activated C kinase 1)

RACK1 is a member of the tryptophan-aspartate (WD) repeat family known for its propeller-like structure. RACK1, a versatile hub in cancer, is found to be involved in neural development and likely to be a potential drug target for AD [76].

#### RAB1A (RAB1A,member RAS oncogene family)

Rab1a gene encodes a member of the Ras superfamily of GTPases. This small GTPase controls vesicle traffic from the endoplasmic reticulum to the Golgi apparatus. It has been implicated in many human diseases, such as cancer, Parkinson’s disease, cardiomyopathy, bacterial, and viral infection [77].

We have performed following 3 different types of biological enrichment analyses based on our novel genes *G*_*N*_. Please, note that we have only retained those terms or pathways having Bonferroni corrected *p <* 0.05. We have found 42 enriched GO-BP terms. Figure 7 illustrates top 20 such terms. Moreover, we have found 5 enriched DO terms as shown in Figure 8. Finally, 41 enriched biological pathways were found from different sources using ConsensusPathDB-human (CPDB-human) database (http://cpdb.molgen.mpg.de/). Details can be found in supplemental information. Table 3 only illustrates the 22 KEGG pathways identified.

**Table 3:**
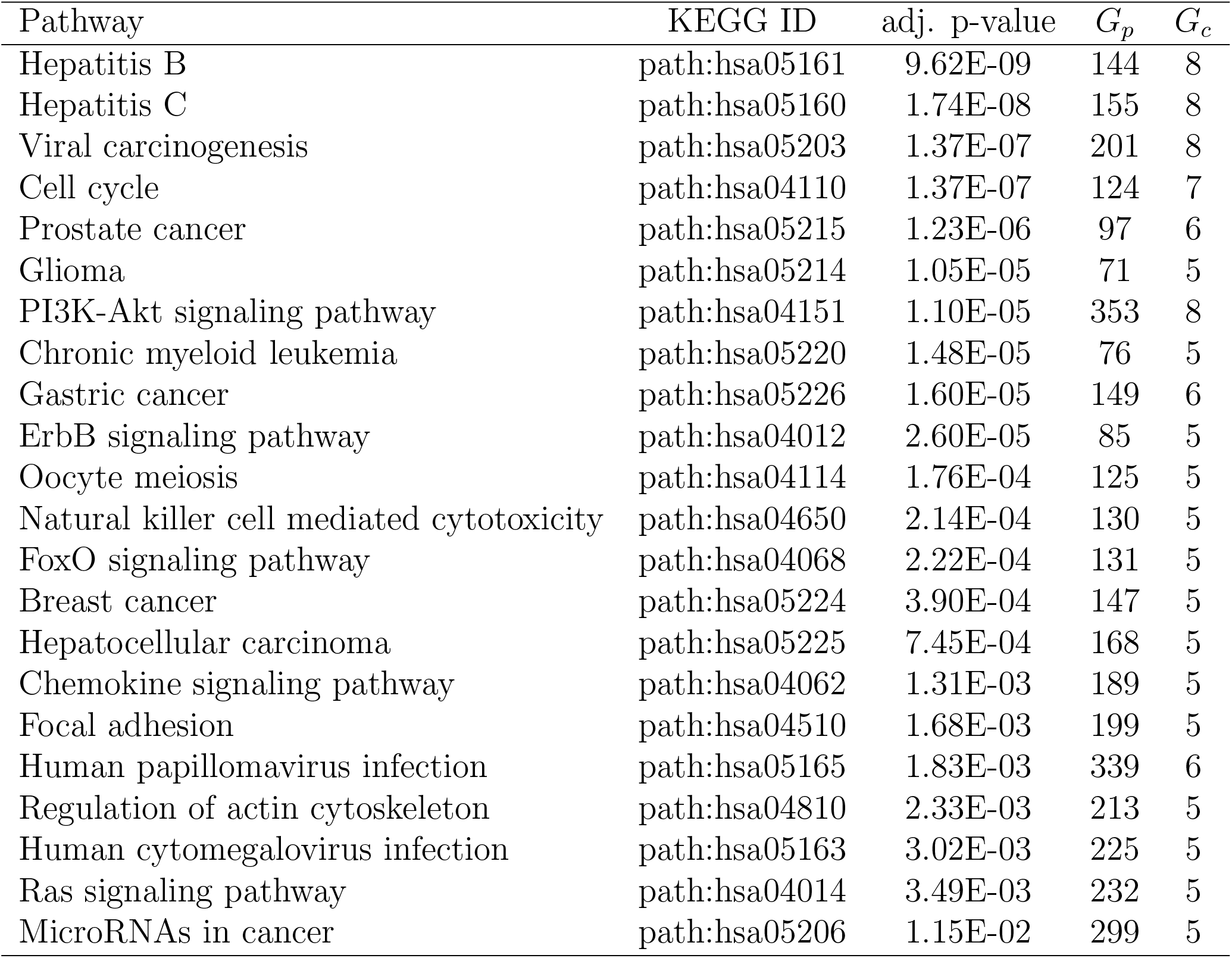
Enriched (Bonferroni corrected *p <* 0.05) KEGG pathways for the novel 24 AD-related genes. *G*_*p*_ refers to the number of genes in a pathway where *G*_*c*_ refers the common genes between the genes in a pathway and AD-related genes.

| Pathway | KEGG ID | adj. p-value | $G_p$ | $G_c$ |
| --- | --- | --- | --- | --- |
| Hepatitis B | path:hsa05161 | 9.62E-09 | 144 | 8 |
| Hepatitis C | path:hsa05160 | 1.74E-08 | 155 | 8 |
| Viral carcinogenesis | path:hsa05203 | 1.37E-07 | 201 | 8 |
| Cell cycle | path:hsa04110 | 1.37E-07 | 124 | 7 |
| Prostate cancer | path:hsa05215 | 1.23E-06 | 97 | 6 |
| Glioma | path:hsa05214 | 1.05E-05 | 71 | 5 |
| PI3K-Akt signaling pathway | path:hsa04151 | 1.10E-05 | 353 | 8 |
| Chronic myeloid leukemia | path:hsa05220 | 1.48E-05 | 76 | 5 |
| Gastric cancer | path:hsa05226 | 1.60E-05 | 149 | 6 |
| ErbB signaling pathway | path:hsa04012 | 2.60E-05 | 85 | 5 |
| Oocyte meiosis | path:hsa04114 | 1.76E-04 | 125 | 5 |
| Natural killer cell mediated cytotoxicity | path:hsa04650 | 2.14E-04 | 130 | 5 |
| FoxO signaling pathway | path:hsa04068 | 2.22E-04 | 131 | 5 |
| Breast cancer | path:hsa05224 | 3.90E-04 | 147 | 5 |
| Hepatocellular carcinoma | path:hsa05225 | 7.45E-04 | 168 | 5 |
| Chemokine signaling pathway | path:hsa04062 | 1.31E-03 | 189 | 5 |
| Focal adhesion | path:hsa04510 | 1.68E-03 | 199 | 5 |
| Human papillomavirus infection | path:hsa05165 | 1.83E-03 | 339 | 6 |
| Regulation of actin cytoskeleton | path:hsa04810 | 2.33E-03 | 213 | 5 |
| Human cytomegalovirus infection | path:hsa05163 | 3.02E-03 | 225 | 5 |
| Ras signaling pathway | path:hsa04014 | 3.49E-03 | 232 | 5 |
| MicroRNAs in cancer | path:hsa05206 | 1.15E-02 | 299 | 5 |

**Figure 7:**
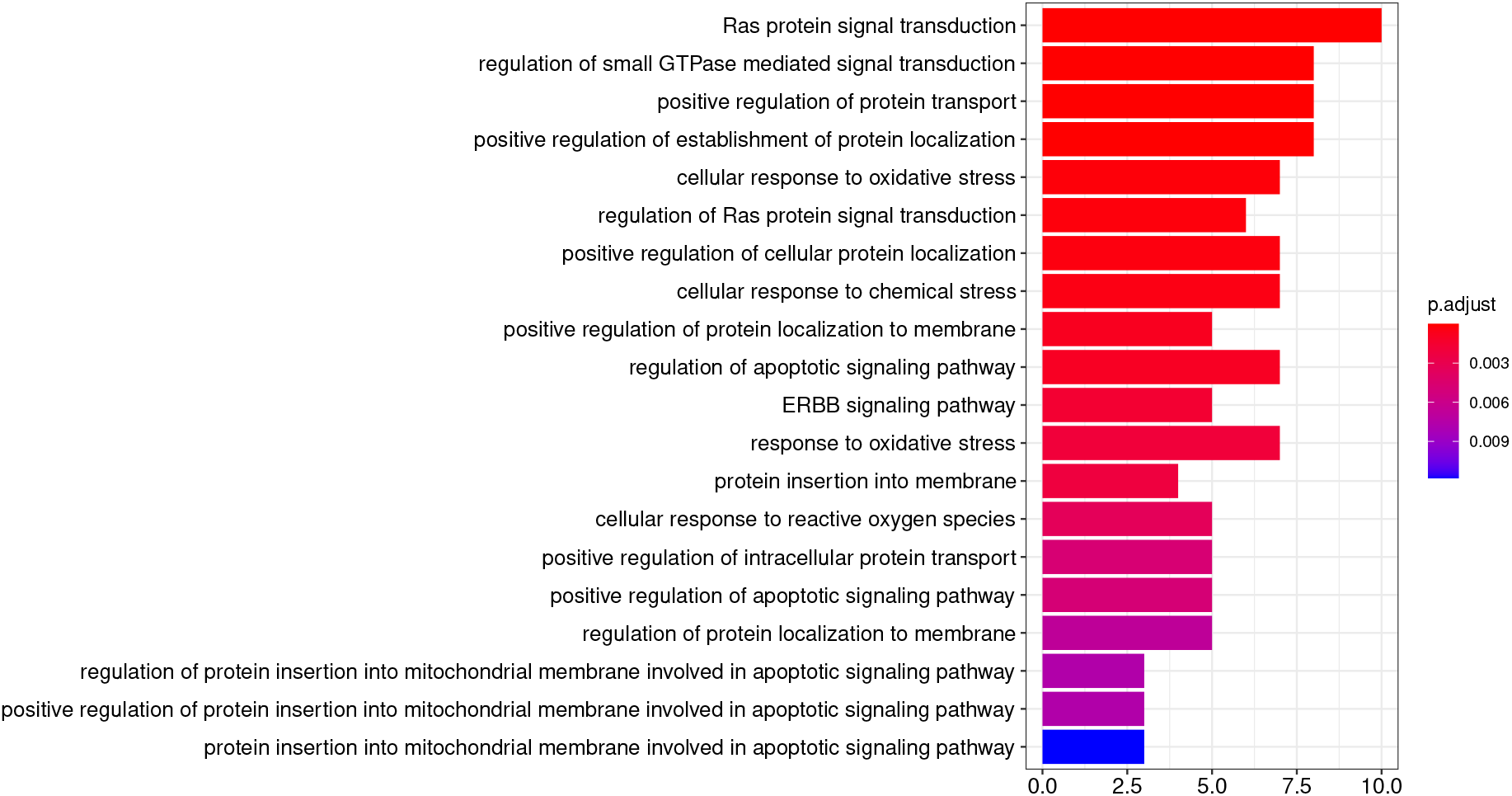
Top 20 enriched (Bonferroni corrected *p <* 0.05) GO-BP terms based on 24 novel AD-related genes.

**Figure 8:**
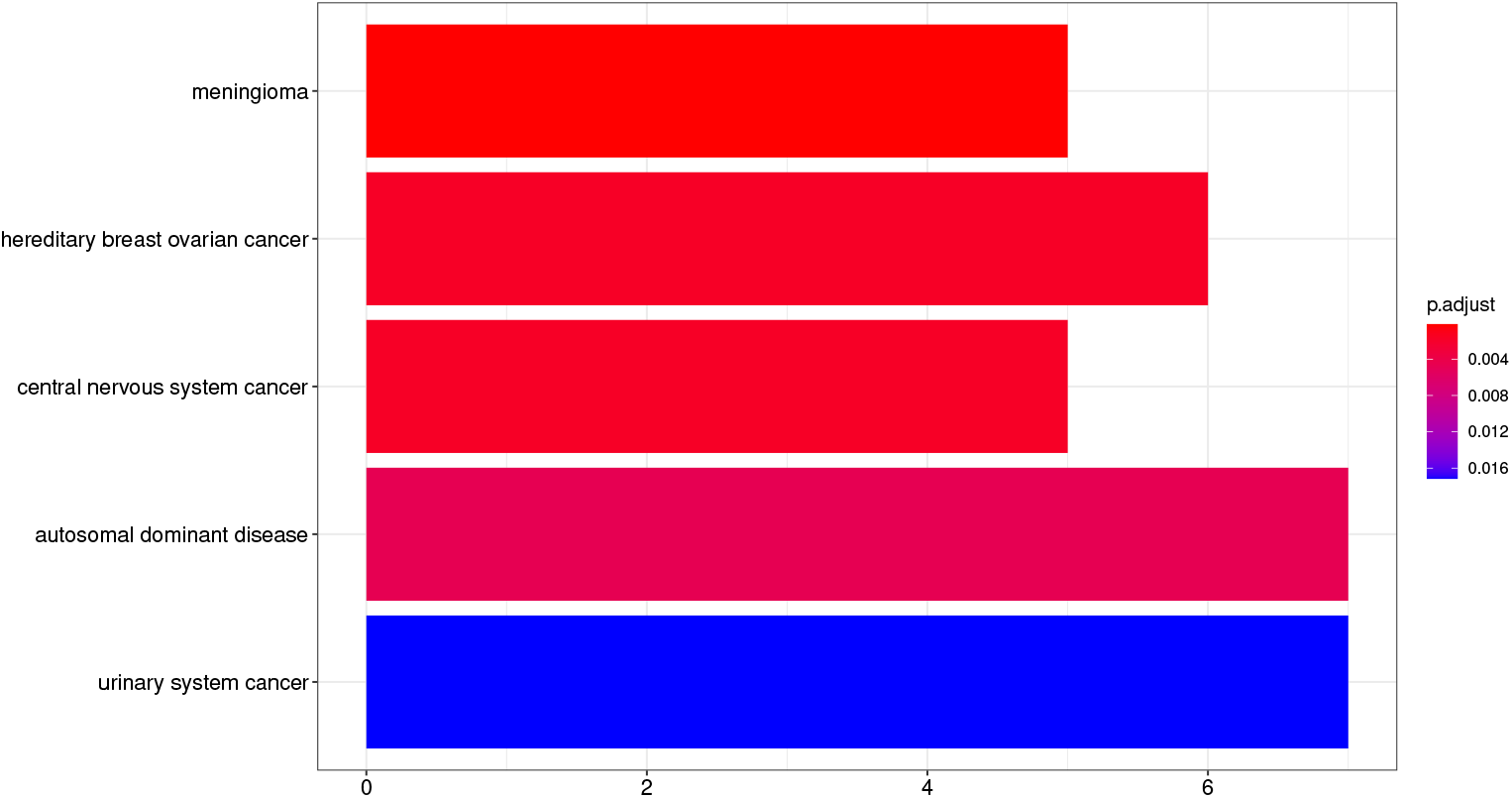
Enriched (Bonferroni corrected *p <* 0.05) DO terms based on 24 novel AD-related genes.

## Discussion

In this article, we have proposed a novel graph theory-based algorithm to elucidate complex network of biological pathways. We have shown how to cleverly construct a weighted network based on the existing biological information embedded in the pathway. We claim that if the assignment of weights towards the edges are biologically synonymous, the analyses performed on the network will be highly biologically relevant. Moreover, the importance of each pathway is calculated not based on the degree centrality but hinged on the strength of the incident interlink(s) recursively. So, by definition, each important pathway is connected with other important pathway that is highly intuitive not only in biology but also in other domains. A natural next step is to cluster the weighted network into a set of subnetworks that are highly specific with respect to their functional activities. We have found three distinct functional modules that are highly observed in Alzheimer’s disease, e.g. modules related to pathways in metabolism, cancer, and infectious disease. The findings prove the effectiveness of our proposed algorithm. In addition, we do not impose any kind of thresholds on similarity scores to remove edges from the network. Use of a threshold with respect to edge weight is particularly problematic, as it might discard vital edges that could held together a functional module. Furthermore, our algorithm is also robust. In our experiment, GO-BP terms were utilized to compute the similarity between any pair of pathways. We have also used GO-MF terms to compute the similarity scores and we observed similar results with respect to the findings identified by employing GO-BP terms.

In addition, we have also identified a set of 24 disease-specific genes based on the curated list of 242 AD-related genes *G*. Through literature search, we found at least 14 of them are already implicated in Alzheimer’s disease. In addition, some of them are thought to be therapeutic target of AD. One way to measure the similarity between the curated list of genes *G* and novel genes *G*_*N*_ is to compare the commonality between the biological pathways they enriched. We found 45 and 22 KEGG pathways for *G* and *G*_*N*_, respectively. As shown in Figure 9, they have 7 pathways in common. The overlap is statistically significant (*p <* 0.0001) according to the Fisher’s exact test. Note that the number of background pathway is set to 548 (= total number of pathways in KEGG).

**Figure 9:**
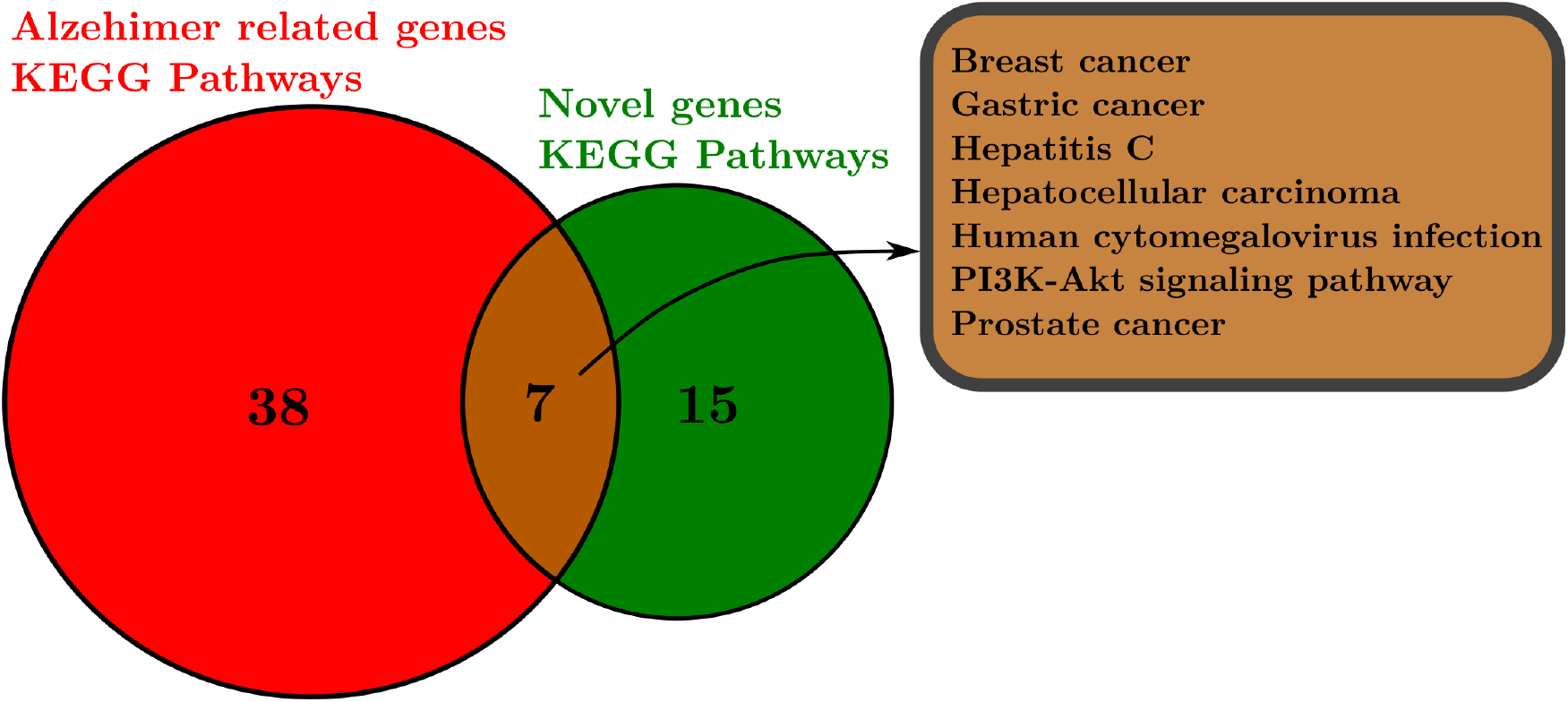
KEGG intersecting pathways between curated and novel AD-specific genes.

## Conclusions

In this article, we have proposed a graph theoretic method to decipher biology of a disease through a carefully curated list of disease-specific genes. We also develop a method to identify a set of novel genes that can further explain the curated list of disease-related genes in a compact way. The experimental evaluations on a set of AD-specific genes disclosed that our methods are indeed effective, reliable, and accurate. We found three functionally different clusters of pathways (e.g. ‘metabolism,” “infection,” and “cancer”) highly involved in the pathogenesis of Alzheimer’s disease. In addition, we have also identified a set of 24 novel genes that are, we believe, highly likely involved in AD regulation.

## Notes

### Competing Interest Statement

The authors have declared no competing interest.

