## Supplemental Tables for "Novel graph theoretic biological pathway network analytics methods for analyzing and discovering Alzheimer’s disease related genes"

### Supplemental file for the paper entitled "Novel graph theoretic biological pathway network analytics methods for analyzing and discovering Alzheimer's disease related genes"

S. Saha et al.

#### Contents

**1 Disease Ontology Terms** **2**

**2 Gene Ontology Terms** **10**

**3 Pathways** **28**

**4 Genes** **31**

**5 Pathways significances** **36**

### 1 Disease Ontology Terms

Table 1: Disease ontology terms (n=488)

| ID | Description | Gene ratio | p-value | Adjusted p-value |
|---|---|---|---|---|
| DOID:10652 | Alzheimer's disease | 117/137 | 3.74E-133 | 3.01E-130 |
| DOID:680 | tauopathy | 117/137 | 1.31E-132 | 1.05E-129 |
| DOID:18 | urinary system disease | 59/137 | 8.82E-37 | 7.09E-34 |
| DOID:1307 | dementia | 39/137 | 1.40E-36 | 1.13E-33 |
| DOID:557 | kidney disease | 58/137 | 1.49E-36 | 1.20E-33 |
| DOID:1936 | atherosclerosis | 52/137 | 1.63E-36 | 1.31E-33 |
| DOID:2348 | arteriosclerotic cardiovascular disease | 52/137 | 1.90E-36 | 1.52E-33 |
| DOID:16 | integumentary system disease | 54/137 | 9.57E-36 | 7.70E-33 |
| DOID:2349 | arteriosclerosis | 52/137 | 9.81E-36 | 7.89E-33 |
| DOID:37 | skin disease | 51/137 | 6.19E-35 | 4.97E-32 |
| DOID:0050890 | synucleinopathy | 43/137 | 9.90E-35 | 7.96E-32 |
| DOID:654 | overnutrition | 49/137 | 2.19E-34 | 1.76E-31 |
| DOID:9970 | obesity | 48/137 | 8.52E-34 | 6.85E-31 |
| DOID:374 | nutrition disease | 49/137 | 2.39E-33 | 1.92E-30 |
| DOID:3393 | coronary artery disease | 49/137 | 9.94E-33 | 7.99E-30 |
| DOID:2237 | hepatitis | 53/137 | 1.47E-32 | 1.18E-29 |
| DOID:850 | lung disease | 55/137 | 2.21E-31 | 1.78E-28 |
| DOID:936 | brain disease | 51/137 | 2.81E-29 | 2.26E-26 |
| DOID:655 | inherited metabolic disorder | 45/137 | 3.31E-29 | 2.66E-26 |
| DOID:2043 | hepatitis B | 37/137 | 4.61E-29 | 3.71E-26 |
| DOID:5844 | myocardial infarction | 42/137 | 5.24E-29 | 4.22E-26 |
| DOID:2621 | autonomic nervous system neoplasm | 47/137 | 5.84E-29 | 4.69E-26 |
| DOID:769 | neuroblastoma | 47/137 | 5.84E-29 | 4.69E-26 |
| DOID:14330 | Parkinson's disease | 36/137 | 1.16E-28 | 9.33E-26 |
| DOID:3083 | chronic obstructive pulmonary disease | 38/137 | 1.92E-28 | 1.54E-25 |
| DOID:1192 | peripheral nervous system neoplasm | 47/137 | 7.00E-28 | 5.62E-25 |
| DOID:28 | endocrine system disease | 47/137 | 9.85E-28 | 7.92E-25 |
| DOID:2320 | obstructive lung disease | 42/137 | 3.24E-27 | 2.61E-24 |
| DOID:3388 | periodontal disease | 31/137 | 9.84E-27 | 7.91E-24 |
| DOID:3082 | interstitial lung disease | 33/137 | 2.29E-26 | 1.84E-23 |
| DOID:104 | bacterial infectious disease | 39/137 | 4.23E-26 | 3.40E-23 |
| DOID:326 | ischemia | 35/137 | 4.77E-26 | 3.84E-23 |
| DOID:0080000 | muscular disease | 45/137 | 3.01E-25 | 2.42E-22 |
| DOID:0050338 | primary bacterial infectious disease | 36/137 | 8.11E-25 | 6.52E-22 |
| DOID:824 | periodontitis | 28/137 | 9.49E-25 | 7.63E-22 |
| DOID:423 | myopathy | 44/137 | 1.07E-24 | 8.62E-22 |
| DOID:66 | muscle tissue disease | 44/137 | 1.07E-24 | 8.62E-22 |
| DOID:1091 | tooth disease | 31/137 | 1.45E-24 | 1.17E-21 |
| DOID:1398 | parasitic infectious disease | 30/137 | 2.26E-24 | 1.81E-21 |
| DOID:633 | myositis | 24/137 | 5.09E-24 | 4.09E-21 |
| DOID:399 | tuberculosis | 30/137 | 7.69E-24 | 6.18E-21 |
| DOID:5082 | liver cirrhosis | 34/137 | 9.77E-24 | 7.85E-21 |
| DOID:403 | mouth disease | 32/137 | 1.10E-23 | 8.84E-21 |
| DOID:10952 | nephritis | 29/137 | 2.19E-23 | 1.76E-20 |
| DOID:263 | kidney cancer | 44/137 | 5.16E-23 | 4.15E-20 |
| DOID:11612 | polycystic ovary syndrome | 30/137 | 5.26E-23 | 4.23E-20 |
| DOID:4195 | hyperglycemia | 28/137 | 7.46E-23 | 6.00E-20 |
| DOID:2789 | parasitic protozoa infectious disease | 27/137 | 1.03E-22 | 8.27E-20 |
| DOID:10283 | prostate cancer | 43/137 | 1.51E-22 | 1.21E-19 |
| DOID:5679 | retinal disease | 40/137 | 1.58E-22 | 1.27E-19 |
| DOID:4451 | renal carcinoma | 41/137 | 1.60E-22 | 1.28E-19 |
| DOID:74 | hematopoietic system disease | 45/137 | 1.68E-22 | 1.35E-19 |
| DOID:0060084 | cell type benign neoplasm | 45/137 | 1.83E-22 | 1.47E-19 |
| DOID:3996 | urinary system cancer | 46/137 | 1.86E-22 | 1.50E-19 |

| | | | | |
|---|---|---|---|---|
| DOID:10591 | pre-eclampsia | 36/137 | 1.99E-22 | 1.60E-19 |
| DOID:50 | thyroid gland disease | 29/137 | 2.31E-22 | 1.85E-19 |
| DOID:120 | female reproductive organ cancer | 44/137 | 2.69E-22 | 2.16E-19 |
| DOID:4450 | renal cell carcinoma | 39/137 | 2.84E-22 | 2.29E-19 |
| DOID:3856 | male reproductive organ cancer | 43/137 | 3.83E-22 | 3.08E-19 |
| DOID:26 | pancreas disease | 29/137 | 1.98E-21 | 1.60E-18 |
| DOID:12365 | malaria | 24/137 | 2.91E-21 | 2.34E-18 |
| DOID:1883 | hepatitis C | 32/137 | 1.03E-20 | 8.30E-18 |
| DOID:0050737 | autosomal recessive disease | 37/137 | 1.16E-20 | 9.35E-18 |
| DOID:13241 | Behcet's disease | 22/137 | 1.64E-20 | 1.31E-17 |
| DOID:3459 | breast carcinoma | 39/137 | 1.81E-20 | 1.45E-17 |
| DOID:0060005 | autoimmune disease of endocrine system | 22/137 | 2.16E-20 | 1.74E-17 |
| DOID:1793 | pancreatic cancer | 36/137 | 2.29E-20 | 1.84E-17 |
| DOID:15 | reproductive system disease | 39/137 | 2.40E-20 | 1.93E-17 |
| DOID:1575 | rheumatic disease | 29/137 | 3.14E-20 | 2.52E-17 |
| DOID:418 | systemic scleroderma | 29/137 | 3.14E-20 | 2.52E-17 |
| DOID:419 | scleroderma | 29/137 | 3.14E-20 | 2.52E-17 |
| DOID:2355 | anemia | 32/137 | 4.34E-20 | 3.49E-17 |
| DOID:9455 | lipid storage disease | 25/137 | 6.27E-20 | 5.04E-17 |
| DOID:3211 | lysosomal storage disease | 26/137 | 1.67E-19 | 1.35E-16 |
| DOID:865 | vasculitis | 24/137 | 2.12E-19 | 1.70E-16 |
| DOID:854 | collagen disease | 29/137 | 2.14E-19 | 1.72E-16 |
| DOID:4989 | pancreatitis | 23/137 | 1.30E-18 | 1.04E-15 |
| DOID:3146 | lipid metabolism disorder | 21/137 | 1.71E-18 | 1.37E-15 |
| DOID:1168 | familial hyperlipidemia | 20/137 | 2.86E-18 | 2.30E-15 |
| DOID:201 | connective tissue cancer | 36/137 | 3.32E-18 | 2.67E-15 |
| DOID:552 | pneumonia | 22/137 | 3.44E-18 | 2.77E-15 |
| DOID:8466 | retinal degeneration | 31/137 | 3.49E-18 | 2.81E-15 |
| DOID:127 | leiomyoma | 21/137 | 3.50E-18 | 2.81E-15 |
| DOID:1686 | glaucoma | 22/137 | 4.25E-18 | 3.42E-15 |
| DOID:3770 | pulmonary fibrosis | 23/137 | 4.26E-18 | 3.42E-15 |
| DOID:1485 | cystic fibrosis | 24/137 | 4.40E-18 | 3.54E-15 |
| DOID:9538 | multiple myeloma | 31/137 | 4.89E-18 | 3.93E-15 |
| DOID:6364 | migraine | 19/137 | 5.53E-18 | 4.44E-15 |
| DOID:3324 | mood disorder | 27/137 | 5.70E-18 | 4.58E-15 |
| DOID:2916 | hypersensitivity reaction type IV disease | 20/137 | 1.30E-17 | 1.04E-14 |
| DOID:0060100 | musculoskeletal system cancer | 38/137 | 1.76E-17 | 1.41E-14 |
| DOID:6713 | cerebrovascular disease | 22/137 | 1.76E-17 | 1.41E-14 |
| DOID:12361 | Graves' disease | 19/137 | 3.64E-17 | 2.93E-14 |
| DOID:8398 | osteoarthritis | 26/137 | 5.03E-17 | 4.04E-14 |
| DOID:2151 | malignant ovarian surface epithelial-stromal neoplasm | 31/137 | 5.05E-17 | 4.06E-14 |
| DOID:2152 | ovary epithelial cancer | 31/137 | 5.05E-17 | 4.06E-14 |
| DOID:4001 | ovarian carcinoma | 31/137 | 5.05E-17 | 4.06E-14 |
| DOID:5517 | stomach carcinoma | 24/137 | 5.31E-17 | 4.27E-14 |
| DOID:2394 | ovarian cancer | 32/137 | 7.47E-17 | 6.00E-14 |
| DOID:526 | Human immunodeficiency virus infectious disease | 25/137 | 8.39E-17 | 6.74E-14 |
| DOID:11335 | sarcoidosis | 19/137 | 9.81E-17 | 7.89E-14 |
| DOID:229 | female reproductive system disease | 26/137 | 1.12E-16 | 8.98E-14 |
| DOID:9352 | type 2 diabetes mellitus | 27/137 | 1.66E-16 | 1.33E-13 |
| DOID:1542 | head and neck carcinoma | 28/137 | 1.72E-16 | 1.38E-13 |
| DOID:5041 | esophageal cancer | 23/137 | 1.96E-16 | 1.58E-13 |
| DOID:0070004 | myeloma | 32/137 | 2.07E-16 | 1.67E-13 |
| DOID:4905 | pancreatic carcinoma | 28/137 | 2.15E-16 | 1.73E-13 |
| DOID:75 | lymphatic system disease | 20/137 | 2.35E-16 | 1.89E-13 |
| DOID:11934 | head and neck cancer | 28/137 | 2.69E-16 | 2.16E-13 |
| DOID:13378 | Kawasaki disease | 16/137 | 2.69E-16 | 2.16E-13 |
| DOID:4960 | bone marrow cancer | 32/137 | 3.25E-16 | 2.61E-13 |
| DOID:750 | peptic ulcer disease | 17/137 | 4.16E-16 | 3.35E-13 |
| DOID:9256 | colorectal cancer | 30/137 | 4.82E-16 | 3.87E-13 |

| | | | | |
|---|---|---|---|---|
| DOID:5672 | large intestine cancer | 30/137 | 5.31E-16 | 4.27E-13 |
| DOID:231 | motor neuron disease | 25/137 | 5.58E-16 | 4.49E-13 |
| DOID:5223 | infertility | 26/137 | 5.72E-16 | 4.60E-13 |
| DOID:9408 | acute myocardial infarction | 19/137 | 6.06E-16 | 4.88E-13 |
| DOID:0050736 | autosomal dominant disease | 35/137 | 6.27E-16 | 5.04E-13 |
| DOID:2723 | dermatitis | 25/137 | 7.22E-16 | 5.80E-13 |
| DOID:657 | adenoma | 32/137 | 9.27E-16 | 7.45E-13 |
| DOID:1602 | lymphadenitis | 16/137 | 9.63E-16 | 7.74E-13 |
| DOID:9942 | lymph node disease | 16/137 | 9.63E-16 | 7.74E-13 |
| DOID:10534 | stomach cancer | 29/137 | 1.15E-15 | 9.24E-13 |
| DOID:1781 | thyroid cancer | 26/137 | 1.31E-15 | 1.05E-12 |
| DOID:870 | neuropathy | 25/137 | 1.35E-15 | 1.09E-12 |
| DOID:7998 | hyperthyroidism | 19/137 | 1.41E-15 | 1.13E-12 |
| DOID:10155 | intestinal cancer | 30/137 | 1.64E-15 | 1.32E-12 |
| DOID:381 | arthropathy | 19/137 | 1.72E-15 | 1.39E-12 |
| DOID:219 | colon cancer | 29/137 | 2.49E-15 | 2.00E-12 |
| DOID:3908 | non-small cell lung carcinoma | 35/137 | 3.18E-15 | 2.56E-12 |
| DOID:184 | bone cancer | 27/137 | 3.79E-15 | 3.04E-12 |
| DOID:2213 | hemorrhagic disease | 22/137 | 6.89E-15 | 5.54E-12 |
| DOID:3310 | atopic dermatitis | 22/137 | 6.89E-15 | 5.54E-12 |
| DOID:1123 | spondyloarthropathy | 17/137 | 6.93E-15 | 5.57E-12 |
| DOID:6432 | pulmonary hypertension | 17/137 | 6.93E-15 | 5.57E-12 |
| DOID:76 | stomach disease | 17/137 | 6.93E-15 | 5.57E-12 |
| DOID:1176 | bronchial disease | 22/137 | 7.94E-15 | 6.38E-12 |
| DOID:332 | amyotrophic lateral sclerosis | 22/137 | 1.05E-14 | 8.44E-12 |
| DOID:1205 | hypersensitivity reaction type I disease | 20/137 | 1.10E-14 | 8.87E-12 |
| DOID:1074 | kidney failure | 22/137 | 1.21E-14 | 9.69E-12 |
| DOID:1115 | sarcoma | 24/137 | 1.21E-14 | 9.74E-12 |
| DOID:1107 | esophageal carcinoma | 20/137 | 1.30E-14 | 1.04E-11 |
| DOID:3717 | gastric adenocarcinoma | 18/137 | 1.53E-14 | 1.23E-11 |
| DOID:0060031 | autoimmune disease of gastrointestinal tract | 19/137 | 1.66E-14 | 1.34E-11 |
| DOID:1247 | blood coagulation disease | 23/137 | 2.53E-14 | 2.03E-11 |
| DOID:3326 | purpura | 16/137 | 3.20E-14 | 2.57E-11 |
| DOID:2377 | multiple sclerosis | 22/137 | 6.62E-14 | 5.32E-11 |
| DOID:5520 | head and neck squamous cell carcinoma | 22/137 | 1.22E-13 | 9.83E-11 |
| DOID:3213 | demyelinating disease | 22/137 | 1.38E-13 | 1.11E-10 |
| DOID:3347 | osteosarcoma | 24/137 | 1.39E-13 | 1.12E-10 |
| DOID:10223 | dermatomyositis | 12/137 | 1.54E-13 | 1.24E-10 |
| DOID:3910 | lung adenocarcinoma | 22/137 | 1.97E-13 | 1.58E-10 |
| DOID:10316 | pneumoconiosis | 12/137 | 2.31E-13 | 1.86E-10 |
| DOID:4074 | pancreas adenocarcinoma | 21/137 | 2.49E-13 | 2.00E-10 |
| DOID:3963 | thyroid carcinoma | 23/137 | 2.51E-13 | 2.02E-10 |
| DOID:974 | upper respiratory tract disease | 19/137 | 2.60E-13 | 2.09E-10 |
| DOID:6590 | spondylitis | 15/137 | 2.98E-13 | 2.39E-10 |
| DOID:3069 | astrocytoma | 18/137 | 3.72E-13 | 2.99E-10 |
| DOID:7147 | ankylosing spondylitis | 15/137 | 3.75E-13 | 3.02E-10 |
| DOID:6000 | congestive heart failure | 24/137 | 5.68E-13 | 4.56E-10 |
| DOID:2234 | focal epilepsy | 14/137 | 6.50E-13 | 5.22E-10 |
| DOID:883 | parasitic helminthiasis infectious disease | 12/137 | 7.17E-13 | 5.77E-10 |
| DOID:631 | fibromyalgia | 10/137 | 8.39E-13 | 6.75E-10 |
| DOID:3328 | temporal lobe epilepsy | 13/137 | 9.41E-13 | 7.57E-10 |
| DOID:1067 | open-angle glaucoma | 14/137 | 1.07E-12 | 8.59E-10 |
| DOID:0060116 | sensory system cancer | 19/137 | 1.60E-12 | 1.29E-09 |
| DOID:2174 | ocular cancer | 19/137 | 1.60E-12 | 1.29E-09 |
| DOID:1564 | fungal infectious disease | 15/137 | 1.70E-12 | 1.37E-09 |
| DOID:10825 | essential hypertension | 18/137 | 1.70E-12 | 1.37E-09 |
| DOID:4607 | biliary tract cancer | 21/137 | 1.85E-12 | 1.49E-09 |
| DOID:10608 | celiac disease | 16/137 | 1.95E-12 | 1.57E-09 |
| DOID:1040 | chronic lymphocytic leukemia | 23/137 | 2.12E-12 | 1.70E-09 |

| | | | | |
|---|---|---|---|---|
| DOID:4029 | gastritis | 14/137 | 2.73E-12 | 2.20E-09 |
| DOID:7166 | thyroiditis | 13/137 | 2.85E-12 | 2.29E-09 |
| DOID:9452 | fatty liver disease | 16/137 | 3.90E-12 | 3.14E-09 |
| DOID:10325 | silicosis | 9/137 | 4.12E-12 | 3.31E-09 |
| DOID:3070 | malignant glioma | 22/137 | 4.41E-12 | 3.54E-09 |
| DOID:11476 | osteoporosis | 16/137 | 4.61E-12 | 3.71E-09 |
| DOID:1184 | nephrotic syndrome | 13/137 | 4.77E-12 | 3.83E-09 |
| DOID:0080011 | bone resorption disease | 16/137 | 5.44E-12 | 4.38E-09 |
| DOID:0060041 | autism spectrum disorder | 21/137 | 6.65E-12 | 5.35E-09 |
| DOID:12849 | autistic disorder | 21/137 | 6.65E-12 | 5.35E-09 |
| DOID:0080008 | ischemic bone disease | 12/137 | 6.88E-12 | 5.53E-09 |
| DOID:2527 | nephrosis | 13/137 | 7.81E-12 | 6.28E-09 |
| DOID:768 | retinoblastoma | 17/137 | 8.25E-12 | 6.63E-09 |
| DOID:771 | retinal cell cancer | 17/137 | 8.25E-12 | 6.63E-09 |
| DOID:2841 | asthma | 18/137 | 8.77E-12 | 7.05E-09 |
| DOID:1037 | lymphoblastic leukemia | 31/137 | 8.79E-12 | 7.07E-09 |
| DOID:3602 | toxic encephalopathy | 12/137 | 9.17E-12 | 7.38E-09 |
| DOID:2462 | retinal vascular disease | 13/137 | 9.92E-12 | 7.98E-09 |
| DOID:8947 | diabetic retinopathy | 13/137 | 9.92E-12 | 7.98E-09 |
| DOID:2163 | nasal cavity disease | 16/137 | 1.04E-11 | 8.33E-09 |
| DOID:2825 | nose disease | 16/137 | 1.04E-11 | 8.33E-09 |
| DOID:4483 | rhinitis | 16/137 | 1.04E-11 | 8.33E-09 |
| DOID:4645 | retinal cancer | 17/137 | 1.10E-11 | 8.80E-09 |
| DOID:10871 | age related macular degeneration | 14/137 | 1.21E-11 | 9.72E-09 |
| DOID:2007 | degeneration of macula and posterior pole | 14/137 | 1.21E-11 | 9.72E-09 |
| DOID:3312 | bipolar disorder | 19/137 | 1.30E-11 | 1.05E-08 |
| DOID:289 | endometriosis | 15/137 | 1.59E-11 | 1.28E-08 |
| DOID:4448 | macular degeneration | 14/137 | 1.79E-11 | 1.44E-08 |
| DOID:4606 | bile duct cancer | 18/137 | 1.87E-11 | 1.50E-08 |
| DOID:4897 | bile duct carcinoma | 18/137 | 1.87E-11 | 1.50E-08 |
| DOID:10286 | prostate carcinoma | 17/137 | 1.90E-11 | 1.53E-08 |
| DOID:341 | peripheral vascular disease | 13/137 | 1.98E-11 | 1.59E-08 |
| DOID:0060040 | pervasive developmental disorder | 21/137 | 1.98E-11 | 1.59E-08 |
| DOID:0080005 | bone remodeling disease | 18/137 | 2.11E-11 | 1.70E-08 |
| DOID:3454 | brain infarction | 13/137 | 2.47E-11 | 1.98E-08 |
| DOID:3620 | central nervous system cancer | 17/137 | 2.48E-11 | 2.00E-08 |
| DOID:1070 | primary open angle glaucoma | 12/137 | 2.70E-11 | 2.17E-08 |
| DOID:2921 | glomerulonephritis | 14/137 | 4.55E-11 | 3.66E-08 |
| DOID:230 | lateral sclerosis | 16/137 | 4.57E-11 | 3.68E-08 |
| DOID:1520 | colon carcinoma | 19/137 | 4.89E-11 | 3.93E-08 |
| DOID:2021 | placenta cancer | 11/137 | 5.01E-11 | 4.03E-08 |
| DOID:3594 | choriocarcinoma | 11/137 | 5.01E-11 | 4.03E-08 |
| DOID:0050136 | systemic mycosis | 11/137 | 6.69E-11 | 5.38E-08 |
| DOID:11077 | brucellosis | 11/137 | 6.69E-11 | 5.38E-08 |
| DOID:0060085 | organ system benign neoplasm | 23/137 | 7.22E-11 | 5.81E-08 |
| DOID:13375 | temporal arteritis | 10/137 | 7.24E-11 | 5.82E-08 |
| DOID:525 | central nervous system vasculitis | 10/137 | 7.24E-11 | 5.82E-08 |
| DOID:0060095 | uterine benign neoplasm | 11/137 | 8.86E-11 | 7.12E-08 |
| DOID:13223 | uterine fibroid | 11/137 | 8.86E-11 | 7.12E-08 |
| DOID:438 | autoimmune disease of the nervous system | 12/137 | 9.07E-11 | 7.30E-08 |
| DOID:4481 | allergic rhinitis | 15/137 | 9.23E-11 | 7.42E-08 |
| DOID:6543 | acne | 9/137 | 1.02E-10 | 8.16E-08 |
| DOID:9098 | sebaceous gland disease | 9/137 | 1.02E-10 | 8.16E-08 |
| DOID:1483 | gingival disease | 10/137 | 1.02E-10 | 8.23E-08 |
| DOID:3526 | cerebral infarction | 12/137 | 1.14E-10 | 9.15E-08 |
| DOID:0060037 | developmental disorder of mental health | 27/137 | 1.14E-10 | 9.19E-08 |
| DOID:784 | chronic kidney failure | 13/137 | 1.25E-10 | 1.01E-07 |
| DOID:13564 | aspergillosis | 8/137 | 1.28E-10 | 1.03E-07 |
| DOID:5295 | intestinal disease | 18/137 | 1.31E-10 | 1.05E-07 |

| | | | | |
|---|---|---|---|---|
| DOID:1927 | sphingolipidosis | 10/137 | 1.43E-10 | 1.15E-07 |
| DOID:0060086 | female reproductive organ benign neoplasm | 11/137 | 1.52E-10 | 1.22E-07 |
| DOID:1826 | epilepsy syndrome | 20/137 | 1.52E-10 | 1.22E-07 |
| DOID:14069 | cerebral malaria | 9/137 | 1.56E-10 | 1.26E-07 |
| DOID:1428 | endocrine pancreas disease | 12/137 | 2.18E-10 | 1.76E-07 |
| DOID:4248 | coronary stenosis | 8/137 | 2.27E-10 | 1.82E-07 |
| DOID:0050622 | reproductive organ benign neoplasm | 11/137 | 2.53E-10 | 2.03E-07 |
| DOID:14250 | Down syndrome | 12/137 | 3.30E-10 | 2.65E-07 |
| DOID:9778 | irritable bowel syndrome | 13/137 | 3.70E-10 | 2.97E-07 |
| DOID:1596 | mental depression | 10/137 | 4.86E-10 | 3.91E-07 |
| DOID:783 | end stage renal failure | 10/137 | 4.86E-10 | 3.91E-07 |
| DOID:866 | vein disease | 10/137 | 4.86E-10 | 3.91E-07 |
| DOID:3748 | esophagus squamous cell carcinoma | 12/137 | 4.91E-10 | 3.95E-07 |
| DOID:799 | varicose veins | 8/137 | 6.34E-10 | 5.10E-07 |
| DOID:2473 | opportunistic mycosis | 10/137 | 6.44E-10 | 5.18E-07 |
| DOID:8567 | Hodgkin's lymphoma | 11/137 | 6.53E-10 | 5.25E-07 |
| DOID:13189 | gout | 9/137 | 7.23E-10 | 5.82E-07 |
| DOID:48 | male reproductive system disease | 16/137 | 7.31E-10 | 5.88E-07 |
| DOID:4752 | multiple system atrophy | 8/137 | 1.01E-09 | 8.12E-07 |
| DOID:12306 | vitiligo | 12/137 | 1.04E-09 | 8.37E-07 |
| DOID:299 | adenocarcinoma | 17/137 | 1.21E-09 | 9.72E-07 |
| DOID:576 | proteinuria | 12/137 | 1.25E-09 | 1.00E-06 |
| DOID:11394 | adult respiratory distress syndrome | 9/137 | 1.41E-09 | 1.14E-06 |
| DOID:5100 | middle ear disease | 9/137 | 1.41E-09 | 1.14E-06 |
| DOID:10923 | sickle cell anemia | 10/137 | 1.43E-09 | 1.15E-06 |
| DOID:11162 | respiratory failure | 11/137 | 1.55E-09 | 1.24E-06 |
| DOID:14221 | metabolic syndrome X | 8/137 | 1.56E-09 | 1.26E-06 |
| DOID:3068 | glioblastoma multiforme | 12/137 | 1.77E-09 | 1.42E-06 |
| DOID:4138 | bile duct disease | 13/137 | 1.82E-09 | 1.46E-06 |
| DOID:8857 | lupus erythematosus | 13/137 | 2.11E-09 | 1.70E-06 |
| DOID:9741 | biliary tract disease | 13/137 | 2.11E-09 | 1.70E-06 |
| DOID:11054 | urinary bladder cancer | 14/137 | 2.96E-09 | 2.38E-06 |
| DOID:574 | peripheral nervous system disease | 12/137 | 3.44E-09 | 2.77E-06 |
| DOID:811 | lipodystrophy | 9/137 | 3.52E-09 | 2.83E-06 |
| DOID:11870 | Pick's disease | 7/137 | 3.70E-09 | 2.98E-06 |
| DOID:1923 | sex differentiation disease | 9/137 | 4.67E-09 | 3.76E-06 |
| DOID:4766 | embryoma | 24/137 | 4.78E-09 | 3.84E-06 |
| DOID:11613 | hyperandrogenism | 8/137 | 5.07E-09 | 4.07E-06 |
| DOID:13810 | familial hypercholesterolemia | 8/137 | 5.07E-09 | 4.07E-06 |
| DOID:345 | uterine disease | 10/137 | 5.82E-09 | 4.68E-06 |
| DOID:8469 | influenza | 14/137 | 6.12E-09 | 4.92E-06 |
| DOID:84 | osteochondritis dissecans | 7/137 | 6.21E-09 | 4.99E-06 |
| DOID:1036 | chronic leukemia | 19/137 | 6.32E-09 | 5.08E-06 |
| DOID:7693 | abdominal aortic aneurysm | 10/137 | 7.20E-09 | 5.79E-06 |
| DOID:9074 | systemic lupus erythematosus | 12/137 | 1.00E-08 | 8.06E-06 |
| DOID:1724 | duodenal ulcer | 8/137 | 1.01E-08 | 8.12E-06 |
| DOID:2742 | auditory system disease | 13/137 | 1.21E-08 | 9.76E-06 |
| DOID:12858 | Huntington's disease | 10/137 | 1.33E-08 | 1.07E-05 |
| DOID:987 | alopecia | 10/137 | 1.33E-08 | 1.07E-05 |
| DOID:688 | embryonal cancer | 24/137 | 1.49E-08 | 1.20E-05 |
| DOID:8692 | myeloid leukemia | 15/137 | 1.57E-08 | 1.26E-05 |
| DOID:0050339 | commensal bacterial infectious disease | 9/137 | 1.68E-08 | 1.35E-05 |
| DOID:12894 | Sjogren's syndrome | 11/137 | 1.92E-08 | 1.55E-05 |
| DOID:2018 | hyperinsulinism | 10/137 | 1.95E-08 | 1.57E-05 |
| DOID:2994 | germ cell cancer | 25/137 | 2.11E-08 | 1.70E-05 |
| DOID:635 | acquired immunodeficiency syndrome | 11/137 | 2.62E-08 | 2.11E-05 |
| DOID:0060058 | lymphoma | 12/137 | 2.63E-08 | 2.11E-05 |
| DOID:3298 | vaccinia | 9/137 | 2.67E-08 | 2.14E-05 |
| DOID:3962 | follicular thyroid carcinoma | 10/137 | 2.82E-08 | 2.27E-05 |

| | | | | |
|---|---|---|---|---|
| DOID:4896 | bile duct adenocarcinoma | 14/137 | 3.12E-08 | 2.51E-05 |
| DOID:4947 | cholangiocarcinoma | 14/137 | 3.12E-08 | 2.51E-05 |
| DOID:612 | primary immunodeficiency disease | 17/137 | 3.26E-08 | 2.62E-05 |
| DOID:8029 | sporadic breast cancer | 8/137 | 3.39E-08 | 2.72E-05 |
| DOID:9744 | type 1 diabetes mellitus | 8/137 | 3.39E-08 | 2.72E-05 |
| DOID:0050589 | inflammatory bowel disease | 12/137 | 3.40E-08 | 2.74E-05 |
| DOID:3087 | gingivitis | 7/137 | 3.51E-08 | 2.82E-05 |
| DOID:8283 | peritonitis | 7/137 | 3.51E-08 | 2.82E-05 |
| DOID:0060049 | autoimmune disease of urogenital tract | 11/137 | 3.54E-08 | 2.85E-05 |
| DOID:12236 | primary biliary cirrhosis | 11/137 | 3.54E-08 | 2.85E-05 |
| DOID:5683 | hereditary breast ovarian cancer | 18/137 | 3.56E-08 | 2.86E-05 |
| DOID:10603 | glucose intolerance | 9/137 | 4.12E-08 | 3.31E-05 |
| DOID:1555 | urticaria | 9/137 | 4.12E-08 | 3.31E-05 |
| DOID:2797 | idiopathic interstitial pneumonia | 8/137 | 4.45E-08 | 3.58E-05 |
| DOID:4535 | hypotrichosis | 10/137 | 4.76E-08 | 3.83E-05 |
| DOID:11123 | Henoch-Schoenlein purpura | 7/137 | 5.07E-08 | 4.07E-05 |
| DOID:13089 | intracranial arterial disease | 7/137 | 5.07E-08 | 4.07E-05 |
| DOID:1557 | hypersensitivity reaction type III disease | 7/137 | 5.07E-08 | 4.07E-05 |
| DOID:3527 | cerebral arterial disease | 7/137 | 5.07E-08 | 4.07E-05 |
| DOID:9809 | hypersensitivity vasculitis | 7/137 | 5.07E-08 | 4.07E-05 |
| DOID:11714 | gestational diabetes | 10/137 | 5.63E-08 | 4.53E-05 |
| DOID:3627 | aortic aneurysm | 10/137 | 5.63E-08 | 4.53E-05 |
| DOID:4007 | bladder carcinoma | 10/137 | 5.63E-08 | 4.53E-05 |
| DOID:10140 | dry eye syndrome | 6/137 | 6.03E-08 | 4.85E-05 |
| DOID:12206 | dengue hemorrhagic fever | 6/137 | 6.03E-08 | 4.85E-05 |
| DOID:1400 | lacrimal apparatus disease | 6/137 | 6.03E-08 | 4.85E-05 |
| DOID:0060036 | intrinsic cardiomyopathy | 14/137 | 6.18E-08 | 4.97E-05 |
| DOID:0080014 | chromosomal disease | 14/137 | 6.18E-08 | 4.97E-05 |
| DOID:3480 | uveal disease | 9/137 | 6.22E-08 | 5.00E-05 |
| DOID:520 | aortic disease | 10/137 | 6.64E-08 | 5.34E-05 |
| DOID:11981 | morbid obesity | 8/137 | 7.47E-08 | 6.00E-05 |
| DOID:4251 | conjunctival disease | 8/137 | 7.47E-08 | 6.00E-05 |
| DOID:13580 | cholestasis | 10/137 | 7.80E-08 | 6.27E-05 |
| DOID:0060122 | integumentary system cancer | 12/137 | 8.98E-08 | 7.22E-05 |
| DOID:4159 | skin cancer | 12/137 | 8.98E-08 | 7.22E-05 |
| DOID:1116 | pertussis | 8/137 | 9.54E-08 | 7.67E-05 |
| DOID:13207 | proliferative diabetic retinopathy | 8/137 | 9.54E-08 | 7.67E-05 |
| DOID:1395 | schistosomiasis | 6/137 | 9.92E-08 | 7.97E-05 |
| DOID:1100 | ovarian disease | 10/137 | 1.07E-07 | 8.58E-05 |
| DOID:421 | hair disease | 10/137 | 1.24E-07 | 1.00E-04 |
| DOID:255 | hemangioma | 8/137 | 1.52E-07 | 1.22E-04 |
| DOID:11729 | Lyme disease | 6/137 | 1.56E-07 | 1.26E-04 |
| DOID:12205 | dengue disease | 6/137 | 1.56E-07 | 1.26E-04 |
| DOID:1443 | cerebral degeneration | 10/137 | 1.67E-07 | 1.34E-04 |
| DOID:2893 | cervix carcinoma | 12/137 | 1.75E-07 | 1.41E-04 |
| DOID:2600 | laryngeal carcinoma | 7/137 | 1.85E-07 | 1.48E-04 |
| DOID:9471 | meningitis | 7/137 | 1.85E-07 | 1.48E-04 |
| DOID:4362 | cervical cancer | 12/137 | 1.95E-07 | 1.57E-04 |
| DOID:0050700 | cardiomyopathy | 14/137 | 2.15E-07 | 1.73E-04 |
| DOID:0060180 | colitis | 10/137 | 2.23E-07 | 1.79E-04 |
| DOID:2030 | anxiety disorder | 10/137 | 2.23E-07 | 1.79E-04 |
| DOID:8577 | ulcerative colitis | 10/137 | 2.23E-07 | 1.79E-04 |
| DOID:0060033 | autoimmune disease of peripheral nervous system | 6/137 | 2.38E-07 | 1.92E-04 |
| DOID:10762 | portal hypertension | 6/137 | 2.38E-07 | 1.92E-04 |
| DOID:10941 | intracranial aneurysm | 6/137 | 2.38E-07 | 1.92E-04 |
| DOID:12842 | Guillain-Barre syndrome | 6/137 | 2.38E-07 | 1.92E-04 |
| DOID:2055 | post-traumatic stress disorder | 6/137 | 2.38E-07 | 1.92E-04 |
| DOID:8689 | anorexia nervosa | 6/137 | 2.38E-07 | 1.92E-04 |
| DOID:9119 | acute myeloid leukemia | 12/137 | 2.40E-07 | 1.93E-04 |

| | | | | |
|---|---|---|---|---|
| DOID:9588 | encephalitis | 9/137 | 3.15E-07 | 2.53E-04 |
| DOID:13025 | retinopathy of prematurity | 5/137 | 3.21E-07 | 2.58E-04 |
| DOID:9446 | cholangitis | 7/137 | 3.23E-07 | 2.60E-04 |
| DOID:9428 | intracranial hypertension | 6/137 | 3.53E-07 | 2.83E-04 |
| DOID:437 | myasthenia gravis | 8/137 | 3.55E-07 | 2.86E-04 |
| DOID:12603 | acute leukemia | 12/137 | 3.98E-07 | 3.20E-04 |
| DOID:9500 | leukocyte disease | 11/137 | 4.06E-07 | 3.26E-04 |
| DOID:1712 | aortic valve stenosis | 7/137 | 4.20E-07 | 3.37E-04 |
| DOID:4798 | aggressive systemic mastocytosis | 7/137 | 4.20E-07 | 3.37E-04 |
| DOID:12217 | Lewy body dementia | 8/137 | 4.33E-07 | 3.48E-04 |
| DOID:439 | neuromuscular junction disease | 8/137 | 4.33E-07 | 3.48E-04 |
| DOID:12449 | aplastic anemia | 10/137 | 4.38E-07 | 3.52E-04 |
| DOID:12930 | dilated cardiomyopathy | 11/137 | 4.53E-07 | 3.65E-04 |
| DOID:3451 | skin carcinoma | 9/137 | 5.05E-07 | 4.06E-04 |
| DOID:1387 | hypolipoproteinemia | 5/137 | 5.80E-07 | 4.66E-04 |
| DOID:4988 | alcoholic pancreatitis | 5/137 | 5.80E-07 | 4.66E-04 |
| DOID:2218 | blood platelet disease | 11/137 | 6.28E-07 | 5.05E-04 |
| DOID:350 | mastocytosis | 8/137 | 6.31E-07 | 5.08E-04 |
| DOID:2583 | agammaglobulinemia | 7/137 | 6.87E-07 | 5.53E-04 |
| DOID:620 | blood protein disease | 7/137 | 6.87E-07 | 5.53E-04 |
| DOID:9120 | amyloidosis | 6/137 | 7.16E-07 | 5.75E-04 |
| DOID:3247 | rhabdomyosarcoma | 10/137 | 8.16E-07 | 6.56E-04 |
| DOID:10124 | corneal disease | 9/137 | 9.08E-07 | 7.30E-04 |
| DOID:12337 | varicocele | 5/137 | 9.80E-07 | 7.88E-04 |
| DOID:14499 | Fabry disease | 5/137 | 9.80E-07 | 7.88E-04 |
| DOID:7188 | autoimmune thyroiditis | 5/137 | 9.80E-07 | 7.88E-04 |
| DOID:9742 | pelvic varices | 5/137 | 9.80E-07 | 7.88E-04 |
| DOID:9743 | diabetic neuropathy | 6/137 | 9.88E-07 | 7.94E-04 |
| DOID:767 | muscular atrophy | 8/137 | 1.07E-06 | 8.62E-04 |
| DOID:1094 | attention deficit hyperactivity disorder | 7/137 | 1.08E-06 | 8.72E-04 |
| DOID:2115 | B cell deficiency | 7/137 | 1.08E-06 | 8.72E-04 |
| DOID:4043 | skeletal muscle cancer | 10/137 | 1.30E-06 | 1.05E-03 |
| DOID:10754 | otitis media | 6/137 | 1.34E-06 | 1.08E-03 |
| DOID:649 | prion disease | 6/137 | 1.34E-06 | 1.08E-03 |
| DOID:13250 | diarrhea | 7/137 | 1.35E-06 | 1.08E-03 |
| DOID:62 | aortic valve disease | 7/137 | 1.35E-06 | 1.08E-03 |
| DOID:12895 | keratoconjunctivitis sicca | 5/137 | 1.57E-06 | 1.26E-03 |
| DOID:6132 | bronchitis | 5/137 | 1.57E-06 | 1.26E-03 |
| DOID:9368 | keratoconjunctivitis | 5/137 | 1.57E-06 | 1.26E-03 |
| DOID:12716 | newborn respiratory distress syndrome | 7/137 | 1.66E-06 | 1.33E-03 |
| DOID:2596 | larynx cancer | 7/137 | 1.66E-06 | 1.33E-03 |
| DOID:3376 | bone osteosarcoma | 8/137 | 1.75E-06 | 1.41E-03 |
| DOID:4914 | esophagus adenocarcinoma | 6/137 | 1.79E-06 | 1.44E-03 |
| DOID:4045 | muscle cancer | 11/137 | 1.88E-06 | 1.51E-03 |
| DOID:4079 | heart valve disease | 8/137 | 2.05E-06 | 1.65E-03 |
| DOID:4948 | gallbladder carcinoma | 8/137 | 2.05E-06 | 1.65E-03 |
| DOID:3969 | papillary thyroid carcinoma | 11/137 | 2.06E-06 | 1.66E-03 |
| DOID:272 | hepatic vascular disease | 6/137 | 2.35E-06 | 1.89E-03 |
| DOID:3121 | gallbladder cancer | 8/137 | 2.39E-06 | 1.92E-03 |
| DOID:9415 | allergic asthma | 8/137 | 2.39E-06 | 1.92E-03 |
| DOID:1019 | osteomyelitis | 5/137 | 2.41E-06 | 1.94E-03 |
| DOID:121 | vaginal disease | 5/137 | 2.41E-06 | 1.94E-03 |
| DOID:13976 | peptic esophagitis | 5/137 | 2.41E-06 | 1.94E-03 |
| DOID:2170 | vaginitis | 5/137 | 2.41E-06 | 1.94E-03 |
| DOID:2913 | acute pancreatitis | 5/137 | 2.41E-06 | 1.94E-03 |
| DOID:3385 | bacterial vaginosis | 5/137 | 2.41E-06 | 1.94E-03 |
| DOID:4330 | non-langerhans-cell histiocytosis | 5/137 | 2.41E-06 | 1.94E-03 |
| DOID:2277 | gonadal disease | 9/137 | 2.59E-06 | 2.09E-03 |
| DOID:349 | systemic mastocytosis | 7/137 | 2.98E-06 | 2.40E-03 |

| | | | | |
|---|---|---|---|---|
| DOID:13515 | tuberous sclerosis | 6/137 | 3.05E-06 | 2.45E-03 |
| DOID:687 | hepatoblastoma | 6/137 | 3.05E-06 | 2.45E-03 |
| DOID:5409 | lung small cell carcinoma | 9/137 | 3.30E-06 | 2.65E-03 |
| DOID:1884 | viral hepatitis | 5/137 | 3.57E-06 | 2.87E-03 |
| DOID:5183 | hereditary Wilms' tumor | 8/137 | 3.70E-06 | 2.98E-03 |
| DOID:6050 | esophageal disease | 8/137 | 3.70E-06 | 2.98E-03 |
| DOID:0002116 | pterygium | 6/137 | 3.91E-06 | 3.14E-03 |
| DOID:10139 | conjunctival degeneration | 6/137 | 3.91E-06 | 3.14E-03 |
| DOID:10526 | conjunctival pterygium | 6/137 | 3.91E-06 | 3.14E-03 |
| DOID:13809 | familial combined hyperlipidemia | 6/137 | 3.91E-06 | 3.14E-03 |
| DOID:4866 | salivary gland adenoid cystic carcinoma | 7/137 | 4.28E-06 | 3.45E-03 |
| DOID:1024 | leprosy | 7/137 | 5.09E-06 | 4.10E-03 |
| DOID:10808 | gastric ulcer | 5/137 | 5.12E-06 | 4.11E-03 |
| DOID:1993 | rectum cancer | 5/137 | 5.12E-06 | 4.11E-03 |
| DOID:2154 | nephroblastoma | 9/137 | 5.81E-06 | 4.67E-03 |
| DOID:13141 | uveitis | 6/137 | 6.21E-06 | 5.00E-03 |
| DOID:8670 | eating disorder | 6/137 | 6.21E-06 | 5.00E-03 |
| DOID:10976 | membranous glomerulonephritis | 5/137 | 7.15E-06 | 5.75E-03 |
| DOID:1969 | cerebral palsy | 5/137 | 7.15E-06 | 5.75E-03 |
| DOID:9253 | gastrointestinal stromal tumor | 5/137 | 7.15E-06 | 5.75E-03 |
| DOID:11260 | rabies | 8/137 | 7.24E-06 | 5.82E-03 |
| DOID:11963 | esophagitis | 6/137 | 7.73E-06 | 6.21E-03 |
| DOID:12177 | common variable immunodeficiency | 6/137 | 7.73E-06 | 6.21E-03 |
| DOID:8618 | oral cavity cancer | 8/137 | 8.21E-06 | 6.60E-03 |
| DOID:1459 | hypothyroidism | 7/137 | 8.32E-06 | 6.69E-03 |
| DOID:3371 | chondrosarcoma | 8/137 | 1.05E-05 | 8.44E-03 |
| DOID:440 | neuromuscular disease | 9/137 | 1.09E-05 | 8.73E-03 |
| DOID:319 | spinal cord disease | 6/137 | 1.16E-05 | 9.36E-03 |
| DOID:303 | substance-related disorder | 8/137 | 1.18E-05 | 9.51E-03 |
| DOID:10930 | borderline personality disorder | 5/137 | 1.31E-05 | 1.05E-02 |
| DOID:2957 | pulmonary tuberculosis | 5/137 | 1.31E-05 | 1.05E-02 |
| DOID:12336 | male infertility | 10/137 | 1.44E-05 | 1.16E-02 |
| DOID:0050598 | extrapulmonary tuberculosis | 4/137 | 1.59E-05 | 1.28E-02 |
| DOID:106 | pleural tuberculosis | 4/137 | 1.59E-05 | 1.28E-02 |
| DOID:10908 | hydrocephalus | 4/137 | 1.59E-05 | 1.28E-02 |
| DOID:1159 | functional gastric disease | 4/137 | 1.59E-05 | 1.28E-02 |
| DOID:1229 | paranoid schizophrenia | 4/137 | 1.59E-05 | 1.28E-02 |
| DOID:14203 | childhood type dermatomyositis | 4/137 | 1.59E-05 | 1.28E-02 |
| DOID:14256 | adult-onset Still's disease | 4/137 | 1.59E-05 | 1.28E-02 |
| DOID:9993 | hypoglycemia | 4/137 | 1.59E-05 | 1.28E-02 |
| DOID:0060115 | nervous system benign neoplasm | 6/137 | 1.70E-05 | 1.37E-02 |
| DOID:10883 | herpangina | 5/137 | 1.72E-05 | 1.38E-02 |
| DOID:12140 | Chagas disease | 5/137 | 1.72E-05 | 1.38E-02 |
| DOID:12252 | Cushing's syndrome | 5/137 | 1.72E-05 | 1.38E-02 |
| DOID:14067 | Plasmodium falciparum malaria | 5/137 | 1.72E-05 | 1.38E-02 |
| DOID:1510 | personality disorder | 5/137 | 1.72E-05 | 1.38E-02 |
| DOID:8632 | Kaposi's sarcoma | 5/137 | 1.72E-05 | 1.38E-02 |
| DOID:9007 | sudden infant death syndrome | 5/137 | 1.72E-05 | 1.38E-02 |
| DOID:1588 | thrombocytopenia | 8/137 | 1.86E-05 | 1.50E-02 |
| DOID:0050624 | gastrointestinal system benign neoplasm | 7/137 | 1.99E-05 | 1.60E-02 |
| DOID:8778 | Crohn's disease | 7/137 | 1.99E-05 | 1.60E-02 |
| DOID:10113 | trypanosomiasis | 5/137 | 2.22E-05 | 1.79E-02 |
| DOID:1272 | telangiectasis | 6/137 | 2.43E-05 | 1.95E-02 |
| DOID:11396 | pulmonary edema | 4/137 | 2.47E-05 | 1.98E-02 |
| DOID:1586 | rheumatic fever | 4/137 | 2.47E-05 | 1.98E-02 |
| DOID:3369 | peripheral primitive neuroectodermal tumor | 5/137 | 2.84E-05 | 2.28E-02 |
| DOID:594 | panic disorder | 6/137 | 2.87E-05 | 2.31E-02 |
| DOID:3744 | cervical squamous cell carcinoma | 8/137 | 3.16E-05 | 2.54E-02 |
| DOID:0060262 | gallbladder disease | 6/137 | 3.38E-05 | 2.72E-02 |

| | | | | |
|---|---|---|---|---|
| DOID:3405 | histiocytosis | 6/137 | 3.38E-05 | 2.72E-02 |
| DOID:2048 | autoimmune hepatitis | 5/137 | 3.58E-05 | 2.88E-02 |
| DOID:14018 | alcoholic liver cirrhosis | 4/137 | 3.65E-05 | 2.93E-02 |
| DOID:14504 | Niemann-Pick disease | 4/137 | 3.65E-05 | 2.93E-02 |
| DOID:4305 | bone giant cell tumor | 4/137 | 3.65E-05 | 2.93E-02 |
| DOID:1380 | endometrial cancer | 9/137 | 3.86E-05 | 3.10E-02 |
| DOID:9261 | nasopharynx carcinoma | 6/137 | 3.96E-05 | 3.18E-02 |
| DOID:363 | uterine cancer | 9/137 | 4.19E-05 | 3.37E-02 |
| DOID:0060089 | endocrine organ benign neoplasm | 9/137 | 4.54E-05 | 3.65E-02 |
| DOID:3565 | meningioma | 8/137 | 4.66E-05 | 3.75E-02 |
| DOID:0050904 | salivary gland carcinoma | 7/137 | 4.77E-05 | 3.83E-02 |
| DOID:12571 | phacogenic glaucoma | 4/137 | 5.20E-05 | 4.18E-02 |
| DOID:13641 | exfoliation syndrome | 4/137 | 5.20E-05 | 4.18E-02 |
| DOID:1803 | neuritis | 4/137 | 5.20E-05 | 4.18E-02 |
| DOID:288 | endometriosis of uterus | 4/137 | 5.20E-05 | 4.18E-02 |
| DOID:3086 | gingival overgrowth | 4/137 | 5.20E-05 | 4.18E-02 |
| DOID:678 | progressive supranuclear palsy | 4/137 | 5.20E-05 | 4.18E-02 |
| DOID:3829 | pituitary adenoma | 7/137 | 5.35E-05 | 4.30E-02 |
| DOID:8850 | salivary gland cancer | 7/137 | 5.35E-05 | 4.30E-02 |
| DOID:0060119 | pharynx cancer | 6/137 | 5.35E-05 | 4.30E-02 |
| DOID:12704 | ataxia telangiectasia | 6/137 | 5.35E-05 | 4.30E-02 |

Table 2: Novel disease ontology terms (n=5)

| ID | Description | Gene ratio | p-value | Adjusted p-value |
|---|---|---|---|---|
| DOID:3565 | meningioma | 5/18 | 5.60E-07 | 1.56E-04 |
| DOID:5683 | hereditary breast ovarian cancer | 6/18 | 5.96E-06 | 1.66E-03 |
| DOID:3620 | central nervous system cancer | 5/18 | 5.99E-06 | 1.67E-03 |
| DOID:0050736 | autosomal dominant disease | 7/18 | 1.68E-05 | 4.67E-03 |
| DOID:3996 | urinary system cancer | 7/18 | 6.17E-05 | 1.72E-02 |

### 2 Gene Ontology Terms

Table 3: Gene ontology terms (n=905)

| ID | Description | Gene ratio | p-value | Adjusted p-value |
|---|---|---|---|---|
| GO:0070997 | neuron death | 41/137 | 9.51E-38 | 4.16E-34 |
| GO:1901214 | regulation of neuron death | 35/137 | 1.99E-31 | 8.71E-28 |
| GO:0051402 | neuron apoptotic process | 31/137 | 7.64E-30 | 3.34E-26 |
| GO:0050435 | amyloid-beta metabolic process | 17/137 | 1.87E-24 | 8.17E-21 |
| GO:0042063 | gliogenesis | 29/137 | 3.20E-24 | 1.40E-20 |
| GO:0062012 | regulation of small molecule metabolic process | 33/137 | 7.88E-24 | 3.45E-20 |
| GO:0043523 | regulation of neuron apoptotic process | 25/137 | 3.04E-23 | 1.33E-19 |
| GO:0042110 | T cell activation | 33/137 | 4.86E-23 | 2.12E-19 |
| GO:0150076 | neuroinflammatory response | 18/137 | 1.13E-22 | 4.96E-19 |
| GO:0050727 | regulation of inflammatory response | 31/137 | 1.74E-22 | 7.59E-19 |
| GO:0010001 | glial cell differentiation | 25/137 | 1.87E-22 | 8.17E-19 |
| GO:1901216 | positive regulation of neuron death | 19/137 | 2.74E-22 | 1.20E-18 |
| GO:0050730 | regulation of peptidyl-tyrosine phosphorylation | 26/137 | 2.91E-22 | 1.27E-18 |
| GO:0042493 | response to drug | 30/137 | 3.29E-22 | 1.44E-18 |
| GO:1901653 | cellular response to peptide | 30/137 | 3.54E-22 | 1.55E-18 |
| GO:1904645 | response to amyloid-beta | 16/137 | 7.91E-22 | 3.46E-18 |
| GO:0018108 | peptidyl-tyrosine phosphorylation | 29/137 | 8.63E-22 | 3.77E-18 |

| | | | | |
|---|---|---|---|---|
| GO:1904646 | cellular response to amyloid-beta | 15/137 | 1.07E-21 | 4.69E-18 |
| GO:0018212 | peptidyl-tyrosine modification | 29/137 | 1.08E-21 | 4.72E-18 |
| GO:0050731 | positive regulation of peptidyl-tyrosine phosphorylation | 23/137 | 1.57E-21 | 6.86E-18 |
| GO:0033619 | membrane protein proteolysis | 16/137 | 3.74E-21 | 1.63E-17 |
| GO:0062197 | cellular response to chemical stress | 28/137 | 4.37E-21 | 1.91E-17 |
| GO:0062013 | positive regulation of small molecule metabolic process | 21/137 | 4.50E-21 | 1.97E-17 |
| GO:0050890 | cognition | 26/137 | 1.00E-20 | 4.38E-17 |
| GO:2001233 | regulation of apoptotic signaling pathway | 29/137 | 1.37E-20 | 5.99E-17 |
| GO:0006979 | response to oxidative stress | 30/137 | 1.98E-20 | 8.66E-17 |
| GO:0070661 | leukocyte proliferation | 26/137 | 2.48E-20 | 1.08E-16 |
| GO:0006509 | membrane protein ectodomain proteolysis | 14/137 | 2.55E-20 | 1.12E-16 |
| GO:0032943 | mononuclear cell proliferation | 25/137 | 4.13E-20 | 1.80E-16 |
| GO:0002237 | response to molecule of bacterial origin | 27/137 | 4.62E-20 | 2.02E-16 |
| GO:0072593 | reactive oxygen species metabolic process | 25/137 | 4.89E-20 | 2.14E-16 |
| GO:0042982 | amyloid precursor protein metabolic process | 16/137 | 5.55E-20 | 2.43E-16 |
| GO:2001234 | negative regulation of apoptotic signaling pathway | 23/137 | 9.47E-20 | 4.14E-16 |
| GO:1901215 | negative regulation of neuron death | 22/137 | 2.13E-19 | 9.30E-16 |
| GO:0071902 | positive regulation of protein serine/threonine kinase activity | 26/137 | 2.87E-19 | 1.25E-15 |
| GO:0042987 | amyloid precursor protein catabolic process | 14/137 | 4.31E-19 | 1.88E-15 |
| GO:0046651 | lymphocyte proliferation | 24/137 | 5.05E-19 | 2.21E-15 |
| GO:0007568 | aging | 25/137 | 5.86E-19 | 2.56E-15 |
| GO:0042098 | T cell proliferation | 21/137 | 6.85E-19 | 2.99E-15 |
| GO:0032496 | response to lipopolysaccharide | 25/137 | 1.77E-18 | 7.74E-15 |
| GO:1903409 | reactive oxygen species biosynthetic process | 18/137 | 1.94E-18 | 8.46E-15 |
| GO:0050804 | modulation of chemical synaptic transmission | 28/137 | 2.13E-18 | 9.30E-15 |
| GO:0099177 | regulation of trans-synaptic signaling | 28/137 | 2.26E-18 | 9.86E-15 |
| GO:0043405 | regulation of MAP kinase activity | 25/137 | 3.12E-18 | 1.37E-14 |
| GO:0051222 | positive regulation of protein transport | 25/137 | 7.12E-18 | 3.11E-14 |
| GO:0022407 | regulation of cell-cell adhesion | 27/137 | 9.99E-18 | 4.37E-14 |
| GO:0051235 | maintenance of location | 24/137 | 1.16E-17 | 5.08E-14 |
| GO:0007611 | learning or memory | 22/137 | 1.77E-17 | 7.74E-14 |
| GO:0032147 | activation of protein kinase activity | 24/137 | 1.90E-17 | 8.31E-14 |
| GO:1904951 | positive regulation of establishment of protein localization | 25/137 | 2.04E-17 | 8.89E-14 |
| GO:0050863 | regulation of T cell activation | 24/137 | 2.04E-17 | 8.90E-14 |
| GO:0021782 | glial cell development | 17/137 | 2.11E-17 | 9.24E-14 |
| GO:0097191 | extrinsic apoptotic signaling pathway | 21/137 | 2.12E-17 | 9.28E-14 |
| GO:2000377 | regulation of reactive oxygen species metabolic process | 20/137 | 2.16E-17 | 9.45E-14 |
| GO:0050673 | epithelial cell proliferation | 27/137 | 2.21E-17 | 9.65E-14 |
| GO:0051091 | positive regulation of DNA-binding transcription factor activity | 22/137 | 3.96E-17 | 1.73E-13 |
| GO:0070663 | regulation of leukocyte proliferation | 21/137 | 5.09E-17 | 2.23E-13 |
| GO:0034599 | cellular response to oxidative stress | 23/137 | 5.71E-17 | 2.49E-13 |
| GO:0019216 | regulation of lipid metabolic process | 26/137 | 6.93E-17 | 3.03E-13 |
| GO:0071216 | cellular response to biotic stimulus | 21/137 | 8.44E-17 | 3.69E-13 |
| GO:0050670 | regulation of lymphocyte proliferation | 20/137 | 1.29E-16 | 5.66E-13 |
| GO:0032944 | regulation of mononuclear cell proliferation | 20/137 | 1.55E-16 | 6.76E-13 |
| GO:0010038 | response to metal ion | 24/137 | 1.88E-16 | 8.21E-13 |
| GO:0051090 | regulation of DNA-binding transcription factor activity | 26/137 | 2.57E-16 | 1.12E-12 |
| GO:0061900 | glial cell activation | 13/137 | 3.38E-16 | 1.48E-12 |
| GO:0031667 | response to nutrient levels | 26/137 | 6.54E-16 | 2.86E-12 |
| GO:0046890 | regulation of lipid biosynthetic process | 19/137 | 7.16E-16 | 3.13E-12 |
| GO:0045834 | positive regulation of lipid metabolic process | 17/137 | 1.05E-15 | 4.60E-12 |
| GO:0051051 | negative regulation of transport | 26/137 | 1.08E-15 | 4.71E-12 |
| GO:0022409 | positive regulation of cell-cell adhesion | 21/137 | 1.09E-15 | 4.74E-12 |
| GO:0048608 | reproductive structure development | 25/137 | 1.38E-15 | 6.02E-12 |
| GO:0001819 | positive regulation of cytokine production | 25/137 | 1.70E-15 | 7.42E-12 |
| GO:0061458 | reproductive system development | 25/137 | 1.70E-15 | 7.42E-12 |
| GO:0048143 | astrocyte activation | 10/137 | 2.22E-15 | 9.70E-12 |
| GO:0009896 | positive regulation of catabolic process | 25/137 | 2.43E-15 | 1.06E-11 |

| | | | | |
|---|---|---|---|---|
| GO:1903037 | regulation of leukocyte cell-cell adhesion | 22/137 | 2.54E-15 | 1.11E-11 |
| GO:0034205 | amyloid-beta formation | 11/137 | 2.78E-15 | 1.21E-11 |
| GO:0008202 | steroid metabolic process | 22/137 | 3.07E-15 | 1.34E-11 |
| GO:0018209 | peptidyl-serine modification | 22/137 | 3.26E-15 | 1.43E-11 |
| GO:1903829 | positive regulation of cellular protein localization | 22/137 | 4.45E-15 | 1.94E-11 |
| GO:0042129 | regulation of T cell proliferation | 17/137 | 4.64E-15 | 2.03E-11 |
| GO:0043406 | positive regulation of MAP kinase activity | 20/137 | 4.86E-15 | 2.12E-11 |
| GO:0045785 | positive regulation of cell adhesion | 24/137 | 6.26E-15 | 2.74E-11 |
| GO:0009266 | response to temperature stimulus | 19/137 | 6.40E-15 | 2.80E-11 |
| GO:0015850 | organic hydroxy compound transport | 20/137 | 6.48E-15 | 2.83E-11 |
| GO:0032602 | chemokine production | 13/137 | 7.54E-15 | 3.30E-11 |
| GO:0050806 | positive regulation of synaptic transmission | 17/137 | 7.63E-15 | 3.33E-11 |
| GO:0031331 | positive regulation of cellular catabolic process | 23/137 | 8.33E-15 | 3.64E-11 |
| GO:0018105 | peptidyl-serine phosphorylation | 21/137 | 8.93E-15 | 3.90E-11 |
| GO:0150077 | regulation of neuroinflammatory response | 11/137 | 9.29E-15 | 4.06E-11 |
| GO:0050769 | positive regulation of neurogenesis | 25/137 | 1.10E-14 | 4.82E-11 |
| GO:0071222 | cellular response to lipopolysaccharide | 18/137 | 1.23E-14 | 5.36E-11 |
| GO:0043491 | protein kinase B signaling | 20/137 | 1.30E-14 | 5.70E-11 |
| GO:0033002 | muscle cell proliferation | 19/137 | 1.49E-14 | 6.51E-11 |
| GO:0014002 | astrocyte development | 11/137 | 1.62E-14 | 7.08E-11 |
| GO:0043434 | response to peptide hormone | 24/137 | 1.63E-14 | 7.14E-11 |
| GO:0007159 | leukocyte cell-cell adhesion | 22/137 | 2.05E-14 | 8.95E-11 |
| GO:0014065 | phosphatidylinositol 3-kinase signaling | 16/137 | 2.23E-14 | 9.77E-11 |
| GO:0051098 | regulation of binding | 22/137 | 2.42E-14 | 1.06E-10 |
| GO:0071219 | cellular response to molecule of bacterial origin | 18/137 | 3.82E-14 | 1.67E-10 |
| GO:2001237 | negative regulation of extrinsic apoptotic signaling pathway | 14/137 | 3.95E-14 | 1.72E-10 |
| GO:0052547 | regulation of peptidase activity | 24/137 | 4.07E-14 | 1.78E-10 |
| GO:0048167 | regulation of synaptic plasticity | 17/137 | 4.39E-14 | 1.92E-10 |
| GO:0043467 | regulation of generation of precursor metabolites and energy | 16/137 | 4.99E-14 | 2.18E-10 |
| GO:0033138 | positive regulation of peptidyl-serine phosphorylation | 14/137 | 5.14E-14 | 2.25E-10 |
| GO:0006816 | calcium ion transport | 23/137 | 5.43E-14 | 2.37E-10 |
| GO:0006066 | alcohol metabolic process | 22/137 | 6.44E-14 | 2.81E-10 |
| GO:0014068 | positive regulation of phosphatidylinositol 3-kinase signaling | 13/137 | 6.61E-14 | 2.89E-10 |
| GO:0046889 | positive regulation of lipid biosynthetic process | 13/137 | 6.61E-14 | 2.89E-10 |
| GO:0032642 | regulation of chemokine production | 12/137 | 6.79E-14 | 2.97E-10 |
| GO:0048511 | rhythmic process | 20/137 | 7.54E-14 | 3.30E-10 |
| GO:0032102 | negative regulation of response to external stimulus | 23/137 | 7.66E-14 | 3.35E-10 |
| GO:0097242 | amyloid-beta clearance | 10/137 | 8.62E-14 | 3.77E-10 |
| GO:0016125 | sterol metabolic process | 16/137 | 9.72E-14 | 4.25E-10 |
| GO:1903039 | positive regulation of leukocyte cell-cell adhesion | 18/137 | 1.02E-13 | 4.48E-10 |
| GO:0010876 | lipid localization | 23/137 | 1.07E-13 | 4.69E-10 |
| GO:0050678 | regulation of epithelial cell proliferation | 22/137 | 1.08E-13 | 4.73E-10 |
| GO:0010959 | regulation of metal ion transport | 22/137 | 1.14E-13 | 4.98E-10 |
| GO:1905952 | regulation of lipid localization | 16/137 | 1.17E-13 | 5.11E-10 |
| GO:0106027 | neuron projection organization | 13/137 | 1.20E-13 | 5.24E-10 |
| GO:1900407 | regulation of cellular response to oxidative stress | 13/137 | 1.20E-13 | 5.24E-10 |
| GO:0048732 | gland development | 23/137 | 1.24E-13 | 5.41E-10 |
| GO:0048660 | regulation of smooth muscle cell proliferation | 16/137 | 1.40E-13 | 6.13E-10 |
| GO:0051099 | positive regulation of binding | 16/137 | 1.54E-13 | 6.71E-10 |
| GO:0033135 | regulation of peptidyl-serine phosphorylation | 15/137 | 1.61E-13 | 7.04E-10 |
| GO:0048659 | smooth muscle cell proliferation | 16/137 | 1.68E-13 | 7.34E-10 |
| GO:0046660 | female sex differentiation | 14/137 | 1.78E-13 | 7.79E-10 |
| GO:0032612 | interleukin-1 production | 14/137 | 2.26E-13 | 9.85E-10 |
| GO:0043524 | negative regulation of neuron apoptotic process | 15/137 | 2.41E-13 | 1.06E-09 |
| GO:0070371 | ERK1 and ERK2 cascade | 20/137 | 2.48E-13 | 1.08E-09 |
| GO:0031293 | membrane protein intracellular domain proteolysis | 8/137 | 2.59E-13 | 1.13E-09 |
| GO:0071496 | cellular response to external stimulus | 20/137 | 2.63E-13 | 1.15E-09 |
| GO:0045862 | positive regulation of proteolysis | 21/137 | 2.88E-13 | 1.26E-09 |
| GO:0008203 | cholesterol metabolic process | 15/137 | 3.58E-13 | 1.56E-09 |

| | | | | |
|---|---|---|---|---|
| GO:0008585 | female gonad development | 13/137 | 3.63E-13 | 1.59E-09 |
| GO:0051924 | regulation of calcium ion transport | 18/137 | 3.64E-13 | 1.59E-09 |
| GO:1902882 | regulation of response to oxidative stress | 13/137 | 4.15E-13 | 1.81E-09 |
| GO:0070838 | divalent metal ion transport | 23/137 | 4.45E-13 | 1.94E-09 |
| GO:0006809 | nitric oxide biosynthetic process | 12/137 | 4.46E-13 | 1.95E-09 |
| GO:0014013 | regulation of gliogenesis | 14/137 | 4.96E-13 | 2.17E-09 |
| GO:0043271 | negative regulation of ion transport | 15/137 | 5.24E-13 | 2.29E-09 |
| GO:0072511 | divalent inorganic cation transport | 23/137 | 6.04E-13 | 2.64E-09 |
| GO:0003018 | vascular process in circulatory system | 16/137 | 6.06E-13 | 2.65E-09 |
| GO:0019915 | lipid storage | 12/137 | 6.08E-13 | 2.66E-09 |
| GO:0002367 | cytokine production involved in immune response | 13/137 | 6.10E-13 | 2.67E-09 |
| GO:0010721 | negative regulation of cell development | 20/137 | 6.76E-13 | 2.95E-09 |
| GO:0050808 | synapse organization | 22/137 | 6.87E-13 | 3.00E-09 |
| GO:0046545 | development of primary female sexual characteristics | 13/137 | 6.92E-13 | 3.02E-09 |
| GO:0048015 | phosphatidylinositol-mediated signaling | 16/137 | 7.12E-13 | 3.11E-09 |
| GO:0052548 | regulation of endopeptidase activity | 22/137 | 7.20E-13 | 3.15E-09 |
| GO:0032722 | positive regulation of chemokine production | 10/137 | 7.51E-13 | 3.28E-09 |
| GO:0032611 | interleukin-1 beta production | 13/137 | 7.84E-13 | 3.43E-09 |
| GO:2000379 | positive regulation of reactive oxygen species metabolic process | 13/137 | 7.84E-13 | 3.43E-09 |
| GO:1902652 | secondary alcohol metabolic process | 15/137 | 8.31E-13 | 3.63E-09 |
| GO:2001236 | regulation of extrinsic apoptotic signaling pathway | 15/137 | 8.31E-13 | 3.63E-09 |
| GO:1903426 | regulation of reactive oxygen species biosynthetic process | 13/137 | 8.87E-13 | 3.88E-09 |
| GO:0046209 | nitric oxide metabolic process | 12/137 | 9.54E-13 | 4.17E-09 |
| GO:0048017 | inositol lipid-mediated signaling | 16/137 | 9.80E-13 | 4.28E-09 |
| GO:0050728 | negative regulation of inflammatory response | 16/137 | 9.80E-13 | 4.28E-09 |
| GO:0016049 | cell growth | 23/137 | 1.01E-12 | 4.41E-09 |
| GO:0031348 | negative regulation of defense response | 18/137 | 1.04E-12 | 4.53E-09 |
| GO:0070482 | response to oxygen levels | 21/137 | 1.07E-12 | 4.65E-09 |
| GO:0048638 | regulation of developmental growth | 20/137 | 1.15E-12 | 5.02E-09 |
| GO:0001935 | endothelial cell proliferation | 16/137 | 1.24E-12 | 5.41E-09 |
| GO:1903532 | positive regulation of secretion by cell | 19/137 | 1.31E-12 | 5.73E-09 |
| GO:0000302 | response to reactive oxygen species | 17/137 | 1.31E-12 | 5.73E-09 |
| GO:0060291 | long-term synaptic potentiation | 12/137 | 1.47E-12 | 6.43E-09 |
| GO:2001057 | reactive nitrogen species metabolic process | 12/137 | 1.47E-12 | 6.43E-09 |
| GO:1900271 | regulation of long-term synaptic potentiation | 10/137 | 1.51E-12 | 6.61E-09 |
| GO:0002696 | positive regulation of leukocyte activation | 21/137 | 1.72E-12 | 7.50E-09 |
| GO:0034612 | response to tumor necrosis factor | 19/137 | 1.94E-12 | 8.47E-09 |
| GO:0035296 | regulation of tube diameter | 14/137 | 2.31E-12 | 1.01E-08 |
| GO:0045598 | regulation of fat cell differentiation | 14/137 | 2.31E-12 | 1.01E-08 |
| GO:0097746 | regulation of blood vessel diameter | 14/137 | 2.31E-12 | 1.01E-08 |
| GO:0002700 | regulation of production of molecular mediator of immune response | 14/137 | 2.55E-12 | 1.11E-08 |
| GO:0035150 | regulation of tube size | 14/137 | 2.55E-12 | 1.11E-08 |
| GO:0045861 | negative regulation of proteolysis | 20/137 | 2.60E-12 | 1.13E-08 |
| GO:0006109 | regulation of carbohydrate metabolic process | 16/137 | 2.83E-12 | 1.24E-08 |
| GO:0002703 | regulation of leukocyte mediated immunity | 16/137 | 3.05E-12 | 1.33E-08 |
| GO:0015918 | sterol transport | 13/137 | 3.18E-12 | 1.39E-08 |
| GO:0002685 | regulation of leukocyte migration | 16/137 | 3.28E-12 | 1.43E-08 |
| GO:0051701 | interaction with host | 16/137 | 3.28E-12 | 1.43E-08 |
| GO:0001558 | regulation of cell growth | 21/137 | 3.28E-12 | 1.43E-08 |
| GO:0048771 | tissue remodeling | 15/137 | 3.29E-12 | 1.44E-08 |
| GO:0050867 | positive regulation of cell activation | 21/137 | 3.43E-12 | 1.50E-08 |
| GO:0071402 | cellular response to lipoprotein particle stimulus | 9/137 | 3.43E-12 | 1.50E-08 |
| GO:0002697 | regulation of immune effector process | 22/137 | 3.50E-12 | 1.53E-08 |
| GO:0009636 | response to toxic substance | 17/137 | 3.56E-12 | 1.55E-08 |
| GO:0097193 | intrinsic apoptotic signaling pathway | 18/137 | 3.68E-12 | 1.61E-08 |
| GO:0070374 | positive regulation of ERK1 and ERK2 cascade | 16/137 | 4.06E-12 | 1.77E-08 |
| GO:0051896 | regulation of protein kinase B signaling | 17/137 | 4.31E-12 | 1.88E-08 |

| | | | | |
|---|---|---|---|---|
| GO:0002791 | regulation of peptide secretion | 20/137 | 4.66E-12 | 2.04E-08 |
| GO:0001936 | regulation of endothelial cell proliferation | 15/137 | 5.32E-12 | 2.33E-08 |
| GO:0031334 | positive regulation of protein-containing complex assembly | 17/137 | 5.54E-12 | 2.42E-08 |
| GO:0036473 | cell death in response to oxidative stress | 12/137 | 5.56E-12 | 2.43E-08 |
| GO:0051047 | positive regulation of secretion | 19/137 | 5.62E-12 | 2.46E-08 |
| GO:0071241 | cellular response to inorganic substance | 16/137 | 6.18E-12 | 2.70E-08 |
| GO:0038034 | signal transduction in absence of ligand | 11/137 | 6.50E-12 | 2.84E-08 |
| GO:0097192 | extrinsic apoptotic signaling pathway in absence of ligand | 11/137 | 6.50E-12 | 2.84E-08 |
| GO:0014066 | regulation of phosphatidylinositol 3-kinase signaling | 13/137 | 8.24E-12 | 3.60E-08 |
| GO:0014015 | positive regulation of gliogenesis | 11/137 | 8.76E-12 | 3.83E-08 |
| GO:1903201 | regulation of oxidative stress-induced cell death | 11/137 | 8.76E-12 | 3.83E-08 |
| GO:0070372 | regulation of ERK1 and ERK2 cascade | 18/137 | 9.05E-12 | 3.95E-08 |
| GO:0042632 | cholesterol homeostasis | 12/137 | 9.07E-12 | 3.96E-08 |
| GO:1901342 | regulation of vasculature development | 21/137 | 9.37E-12 | 4.10E-08 |
| GO:0006631 | fatty acid metabolic process | 20/137 | 9.40E-12 | 4.11E-08 |
| GO:0055092 | sterol homeostasis | 12/137 | 1.02E-11 | 4.46E-08 |
| GO:0050708 | regulation of protein secretion | 19/137 | 1.03E-11 | 4.49E-08 |
| GO:0050807 | regulation of synapse organization | 16/137 | 1.06E-11 | 4.63E-08 |
| GO:0071248 | cellular response to metal ion | 15/137 | 1.06E-11 | 4.64E-08 |
| GO:1905475 | regulation of protein localization to membrane | 15/137 | 1.23E-11 | 5.38E-08 |
| GO:0051251 | positive regulation of lymphocyte activation | 19/137 | 1.31E-11 | 5.74E-08 |
| GO:0043525 | positive regulation of neuron apoptotic process | 10/137 | 1.36E-11 | 5.93E-08 |
| GO:0055088 | lipid homeostasis | 14/137 | 1.39E-11 | 6.08E-08 |
| GO:0001666 | response to hypoxia | 19/137 | 1.45E-11 | 6.32E-08 |
| GO:0030301 | cholesterol transport | 12/137 | 1.45E-11 | 6.33E-08 |
| GO:0007259 | receptor signaling pathway via JAK-STAT | 14/137 | 1.51E-11 | 6.60E-08 |
| GO:0019932 | second-messenger-mediated signaling | 21/137 | 1.55E-11 | 6.76E-08 |
| GO:0009408 | response to heat | 14/137 | 1.78E-11 | 7.78E-08 |
| GO:0010952 | positive regulation of peptidase activity | 15/137 | 1.90E-11 | 8.31E-08 |
| GO:0050671 | positive regulation of lymphocyte proliferation | 13/137 | 1.98E-11 | 8.66E-08 |
| GO:0002718 | regulation of cytokine production involved in immune response | 11/137 | 2.04E-11 | 8.91E-08 |
| GO:0048708 | astrocyte differentiation | 11/137 | 2.04E-11 | 8.91E-08 |
| GO:0002683 | negative regulation of immune system process | 21/137 | 2.06E-11 | 8.99E-08 |
| GO:0002819 | regulation of adaptive immune response | 14/137 | 2.09E-11 | 9.15E-08 |
| GO:0050803 | regulation of synapse structure or activity | 16/137 | 2.15E-11 | 9.41E-08 |
| GO:0007548 | sex differentiation | 17/137 | 2.16E-11 | 9.45E-08 |
| GO:0032946 | positive regulation of mononuclear cell proliferation | 13/137 | 2.18E-11 | 9.51E-08 |
| GO:0045913 | positive regulation of carbohydrate metabolic process | 11/137 | 2.33E-11 | 1.02E-07 |
| GO:0002460 | adaptive immune response based on somatic recombination of immune receptors built from immunoglobulin superfamily domains | 19/137 | 2.44E-11 | 1.07E-07 |
| GO:0036293 | response to decreased oxygen levels | 19/137 | 2.55E-11 | 1.12E-07 |
| GO:0032868 | response to insulin | 17/137 | 2.56E-11 | 1.12E-07 |
| GO:0007220 | Notch receptor processing | 6/137 | 2.69E-11 | 1.18E-07 |
| GO:0043270 | positive regulation of ion transport | 17/137 | 2.71E-11 | 1.18E-07 |
| GO:0002526 | acute inflammatory response | 12/137 | 2.82E-11 | 1.23E-07 |
| GO:0032652 | regulation of interleukin-1 production | 12/137 | 2.82E-11 | 1.23E-07 |
| GO:0051346 | negative regulation of hydrolase activity | 21/137 | 3.07E-11 | 1.34E-07 |
| GO:0071375 | cellular response to peptide hormone stimulus | 18/137 | 3.16E-11 | 1.38E-07 |
| GO:0033344 | cholesterol efflux | 10/137 | 3.17E-11 | 1.39E-07 |
| GO:0097696 | receptor signaling pathway via STAT | 14/137 | 3.37E-11 | 1.47E-07 |
| GO:0046425 | regulation of receptor signaling pathway via JAK-STAT | 13/137 | 3.44E-11 | 1.50E-07 |
| GO:0002573 | myeloid leukocyte differentiation | 15/137 | 3.55E-11 | 1.55E-07 |
| GO:0034764 | positive regulation of transmembrane transport | 15/137 | 3.55E-11 | 1.55E-07 |
| GO:0032768 | regulation of monooxygenase activity | 10/137 | 4.37E-11 | 1.91E-07 |
| GO:0050870 | positive regulation of T cell activation | 15/137 | 4.65E-11 | 2.03E-07 |
| GO:0050768 | negative regulation of neurogenesis | 17/137 | 4.92E-11 | 2.15E-07 |
| GO:0051348 | negative regulation of transferase activity | 17/137 | 5.19E-11 | 2.27E-07 |

| | | | | |
|---|---|---|---|---|
| GO:0090316 | positive regulation of intracellular protein transport | 14/137 | 6.16E-11 | 2.69E-07 |
| GO:0006869 | lipid transport | 19/137 | 6.86E-11 | 3.00E-07 |
| GO:0070665 | positive regulation of leukocyte proliferation | 13/137 | 6.88E-11 | 3.01E-07 |
| GO:0002673 | regulation of acute inflammatory response | 9/137 | 6.92E-11 | 3.02E-07 |
| GO:1904892 | regulation of receptor signaling pathway via STAT | 13/137 | 7.49E-11 | 3.27E-07 |
| GO:0007613 | memory | 12/137 | 7.87E-11 | 3.44E-07 |
| GO:1905954 | positive regulation of lipid localization | 11/137 | 8.15E-11 | 3.56E-07 |
| GO:0042176 | regulation of protein catabolic process | 19/137 | 8.16E-11 | 3.57E-07 |
| GO:0008406 | gonad development | 15/137 | 8.33E-11 | 3.64E-07 |
| GO:0001774 | microglial cell activation | 9/137 | 8.38E-11 | 3.66E-07 |
| GO:0002269 | leukocyte activation involved in inflammatory response | 9/137 | 8.38E-11 | 3.66E-07 |
| GO:0045981 | positive regulation of nucleotide metabolic process | 9/137 | 8.38E-11 | 3.66E-07 |
| GO:1900544 | positive regulation of purine nucleotide metabolic process | 9/137 | 8.38E-11 | 3.66E-07 |
| GO:0043254 | regulation of protein-containing complex assembly | 20/137 | 8.93E-11 | 3.90E-07 |
| GO:0055094 | response to lipoprotein particle | 8/137 | 9.74E-11 | 4.26E-07 |
| GO:0032651 | regulation of interleukin-1 beta production | 11/137 | 1.15E-10 | 5.04E-07 |
| GO:0006874 | cellular calcium ion homeostasis | 20/137 | 1.17E-10 | 5.13E-07 |
| GO:0045137 | development of primary sexual characteristics | 15/137 | 1.21E-10 | 5.29E-07 |
| GO:0050920 | regulation of chemotaxis | 15/137 | 1.21E-10 | 5.29E-07 |
| GO:0050729 | positive regulation of inflammatory response | 13/137 | 1.32E-10 | 5.79E-07 |
| GO:1905477 | positive regulation of protein localization to membrane | 12/137 | 1.39E-10 | 6.09E-07 |
| GO:0051092 | positive regulation of NF-kappaB transcription factor activity | 13/137 | 1.43E-10 | 6.26E-07 |
| GO:0043255 | regulation of carbohydrate biosynthetic process | 11/137 | 1.44E-10 | 6.31E-07 |
| GO:0097305 | response to alcohol | 15/137 | 1.64E-10 | 7.17E-07 |
| GO:0051961 | negative regulation of nervous system development | 17/137 | 1.67E-10 | 7.28E-07 |
| GO:0019217 | regulation of fatty acid metabolic process | 11/137 | 1.80E-10 | 7.86E-07 |
| GO:0042116 | macrophage activation | 11/137 | 1.80E-10 | 7.86E-07 |
| GO:0055074 | calcium ion homeostasis | 20/137 | 1.86E-10 | 8.12E-07 |
| GO:0030098 | lymphocyte differentiation | 18/137 | 1.88E-10 | 8.20E-07 |
| GO:0032872 | regulation of stress-activated MAPK cascade | 15/137 | 1.96E-10 | 8.57E-07 |
| GO:0010623 | programmed cell death involved in cell development | 6/137 | 2.16E-10 | 9.45E-07 |
| GO:0048661 | positive regulation of smooth muscle cell proliferation | 11/137 | 2.23E-10 | 9.74E-07 |
| GO:0098869 | cellular oxidant detoxification | 11/137 | 2.23E-10 | 9.74E-07 |
| GO:0070302 | regulation of stress-activated protein kinase signaling cascade | 15/137 | 2.34E-10 | 1.02E-06 |
| GO:0032885 | regulation of polysaccharide biosynthetic process | 8/137 | 2.56E-10 | 1.12E-06 |
| GO:0032368 | regulation of lipid transport | 12/137 | 2.85E-10 | 1.25E-06 |
| GO:0019233 | sensory perception of pain | 11/137 | 3.05E-10 | 1.33E-06 |
| GO:0042593 | glucose homeostasis | 15/137 | 3.12E-10 | 1.36E-06 |
| GO:0033500 | carbohydrate homeostasis | 15/137 | 3.30E-10 | 1.44E-06 |
| GO:0032635 | interleukin-6 production | 13/137 | 3.30E-10 | 1.44E-06 |
| GO:0042326 | negative regulation of phosphorylation | 20/137 | 3.35E-10 | 1.46E-06 |
| GO:1902105 | regulation of leukocyte differentiation | 16/137 | 3.59E-10 | 1.57E-06 |
| GO:1902991 | regulation of amyloid precursor protein catabolic process | 8/137 | 3.98E-10 | 1.74E-06 |
| GO:0031663 | lipopolysaccharide-mediated signaling pathway | 9/137 | 3.98E-10 | 1.74E-06 |
| GO:0043393 | regulation of protein binding | 14/137 | 4.39E-10 | 1.92E-06 |
| GO:0072503 | cellular divalent inorganic cation homeostasis | 20/137 | 4.46E-10 | 1.95E-06 |
| GO:0006641 | triglyceride metabolic process | 11/137 | 4.56E-10 | 1.99E-06 |
| GO:0050994 | regulation of lipid catabolic process | 9/137 | 4.65E-10 | 2.03E-06 |
| GO:0031668 | cellular response to extracellular stimulus | 15/137 | 4.88E-10 | 2.13E-06 |
| GO:0071356 | cellular response to tumor necrosis factor | 16/137 | 5.08E-10 | 2.22E-06 |
| GO:0097061 | dendritic spine organization | 10/137 | 5.12E-10 | 2.24E-06 |
| GO:0002687 | positive regulation of leukocyte migration | 12/137 | 5.15E-10 | 2.25E-06 |
| GO:0001933 | negative regulation of protein phosphorylation | 19/137 | 5.39E-10 | 2.36E-06 |
| GO:0032092 | positive regulation of protein binding | 10/137 | 5.76E-10 | 2.52E-06 |
| GO:0048545 | response to steroid hormone | 17/137 | 5.82E-10 | 2.54E-06 |
| GO:0051897 | positive regulation of protein kinase B signaling | 13/137 | 5.85E-10 | 2.55E-06 |
| GO:0060401 | cytosolic calcium ion transport | 13/137 | 5.85E-10 | 2.55E-06 |
| GO:0090594 | inflammatory response to wounding | 6/137 | 6.23E-10 | 2.72E-06 |

| | | | | |
|---|---|---|---|---|
| GO:0042509 | regulation of tyrosine phosphorylation of STAT protein | 10/137 | 6.48E-10 | 2.83E-06 |
| GO:1990748 | cellular detoxification | 11/137 | 6.72E-10 | 2.94E-06 |
| GO:0010950 | positive regulation of endopeptidase activity | 13/137 | 7.19E-10 | 3.14E-06 |
| GO:0046688 | response to copper ion | 8/137 | 7.37E-10 | 3.22E-06 |
| GO:0002440 | production of molecular mediator of immune response | 16/137 | 7.86E-10 | 3.43E-06 |
| GO:0031669 | cellular response to nutrient levels | 14/137 | 8.06E-10 | 3.52E-06 |
| GO:0045765 | regulation of angiogenesis | 18/137 | 8.08E-10 | 3.53E-06 |
| GO:0032755 | positive regulation of interleukin-6 production | 10/137 | 8.16E-10 | 3.57E-06 |
| GO:0010675 | regulation of cellular carbohydrate metabolic process | 12/137 | 8.31E-10 | 3.63E-06 |
| GO:0032388 | positive regulation of intracellular transport | 14/137 | 8.55E-10 | 3.74E-06 |
| GO:0032881 | regulation of polysaccharide metabolic process | 8/137 | 8.95E-10 | 3.91E-06 |
| GO:0007260 | tyrosine phosphorylation of STAT protein | 10/137 | 9.14E-10 | 3.99E-06 |
| GO:0021987 | cerebral cortex development | 11/137 | 9.75E-10 | 4.26E-06 |
| GO:0046427 | positive regulation of receptor signaling pathway via JAK-STAT | 10/137 | 1.02E-09 | 4.47E-06 |
| GO:0009306 | protein secretion | 19/137 | 1.05E-09 | 4.57E-06 |
| GO:0032869 | cellular response to insulin stimulus | 14/137 | 1.08E-09 | 4.72E-06 |
| GO:0035592 | establishment of protein localization to extracellular region | 19/137 | 1.08E-09 | 4.73E-06 |
| GO:0002822 | regulation of adaptive immune response based on somatic recombination of immune receptors built from immunoglobulin superfamily domains | 12/137 | 1.22E-09 | 5.33E-06 |
| GO:0032373 | positive regulation of sterol transport | 7/137 | 1.25E-09 | 5.45E-06 |
| GO:0032376 | positive regulation of cholesterol transport | 7/137 | 1.25E-09 | 5.45E-06 |
| GO:0002706 | regulation of lymphocyte mediated immunity | 12/137 | 1.31E-09 | 5.74E-06 |
| GO:0071692 | protein localization to extracellular region | 19/137 | 1.39E-09 | 6.07E-06 |
| GO:0097237 | cellular response to toxic substance | 11/137 | 1.40E-09 | 6.10E-06 |
| GO:0061097 | regulation of protein tyrosine kinase activity | 10/137 | 1.42E-09 | 6.19E-06 |
| GO:1904894 | positive regulation of receptor signaling pathway via STAT | 10/137 | 1.42E-09 | 6.19E-06 |
| GO:0045927 | positive regulation of growth | 15/137 | 1.47E-09 | 6.41E-06 |
| GO:0045428 | regulation of nitric oxide biosynthetic process | 9/137 | 1.48E-09 | 6.45E-06 |
| GO:0050766 | positive regulation of phagocytosis | 9/137 | 1.48E-09 | 6.45E-06 |
| GO:0005979 | regulation of glycogen biosynthetic process | 7/137 | 1.62E-09 | 7.06E-06 |
| GO:0010962 | regulation of glucan biosynthetic process | 7/137 | 1.62E-09 | 7.06E-06 |
| GO:0007204 | positive regulation of cytosolic calcium ion concentration | 16/137 | 1.64E-09 | 7.18E-06 |
| GO:0022612 | gland morphogenesis | 11/137 | 1.66E-09 | 7.26E-06 |
| GO:0009895 | negative regulation of catabolic process | 16/137 | 1.72E-09 | 7.51E-06 |
| GO:0060402 | calcium ion transport into cytosol | 12/137 | 1.76E-09 | 7.71E-06 |
| GO:0045444 | fat cell differentiation | 14/137 | 1.79E-09 | 7.84E-06 |
| GO:0050777 | negative regulation of immune response | 12/137 | 1.90E-09 | 8.29E-06 |
| GO:0032370 | positive regulation of lipid transport | 9/137 | 1.93E-09 | 8.41E-06 |
| GO:0046777 | protein autophosphorylation | 14/137 | 2.00E-09 | 8.75E-06 |
| GO:0006898 | receptor-mediated endocytosis | 16/137 | 2.14E-09 | 9.37E-06 |
| GO:0050764 | regulation of phagocytosis | 10/137 | 2.15E-09 | 9.41E-06 |
| GO:0043112 | receptor metabolic process | 13/137 | 2.16E-09 | 9.45E-06 |
| GO:0032371 | regulation of sterol transport | 9/137 | 2.19E-09 | 9.58E-06 |
| GO:0042531 | positive regulation of tyrosine phosphorylation of STAT protein | 9/137 | 2.19E-09 | 9.58E-06 |
| GO:1990090 | cellular response to nerve growth factor stimulus | 8/137 | 2.21E-09 | 9.65E-06 |
| GO:2000116 | regulation of cysteine-type endopeptidase activity | 14/137 | 2.23E-09 | 9.75E-06 |
| GO:0051403 | stress-activated MAPK cascade | 15/137 | 2.64E-09 | 1.15E-05 |
| GO:0050848 | regulation of calcium-mediated signaling | 10/137 | 2.91E-09 | 1.27E-05 |
| GO:0019218 | regulation of steroid metabolic process | 11/137 | 2.98E-09 | 1.30E-05 |
| GO:2001242 | regulation of intrinsic apoptotic signaling pathway | 12/137 | 3.10E-09 | 1.36E-05 |
| GO:0042102 | positive regulation of T cell proliferation | 10/137 | 3.21E-09 | 1.40E-05 |
| GO:0060191 | regulation of lipase activity | 10/137 | 3.21E-09 | 1.40E-05 |
| GO:1902003 | regulation of amyloid-beta formation | 7/137 | 3.33E-09 | 1.46E-05 |
| GO:0032409 | regulation of transporter activity | 15/137 | 3.34E-09 | 1.46E-05 |
| GO:0042692 | muscle cell differentiation | 17/137 | 3.57E-09 | 1.56E-05 |
| GO:1903706 | regulation of hemopoiesis | 19/137 | 3.59E-09 | 1.57E-05 |

| | | | | |
|---|---|---|---|---|
| GO:0045685 | regulation of glial cell differentiation | 9/137 | 3.61E-09 | 1.58E-05 |
| GO:0050999 | regulation of nitric-oxide synthase activity | 8/137 | 3.62E-09 | 1.58E-05 |
| GO:1990089 | response to nerve growth factor | 8/137 | 3.62E-09 | 1.58E-05 |
| GO:0019058 | viral life cycle | 16/137 | 3.74E-09 | 1.63E-05 |
| GO:0052126 | movement in host environment | 12/137 | 3.81E-09 | 1.66E-05 |
| GO:0032148 | activation of protein kinase B activity | 7/137 | 4.17E-09 | 1.82E-05 |
| GO:0005996 | monosaccharide metabolic process | 15/137 | 4.21E-09 | 1.84E-05 |
| GO:0030307 | positive regulation of cell growth | 12/137 | 4.35E-09 | 1.90E-05 |
| GO:0043122 | regulation of I-kappaB kinase/NF-kappaB signaling | 14/137 | 4.41E-09 | 1.93E-05 |
| GO:0045787 | positive regulation of cell cycle | 17/137 | 4.48E-09 | 1.96E-05 |
| GO:0050714 | positive regulation of protein secretion | 12/137 | 4.65E-09 | 2.03E-05 |
| GO:0010875 | positive regulation of cholesterol efflux | 6/137 | 4.68E-09 | 2.05E-05 |
| GO:0060965 | negative regulation of gene silencing by miRNA | 6/137 | 4.68E-09 | 2.05E-05 |
| GO:0097062 | dendritic spine maintenance | 6/137 | 4.68E-09 | 2.05E-05 |
| GO:0097006 | regulation of plasma lipoprotein particle levels | 10/137 | 4.70E-09 | 2.05E-05 |
| GO:0010506 | regulation of autophagy | 16/137 | 4.80E-09 | 2.10E-05 |
| GO:0021543 | pallium development | 12/137 | 4.96E-09 | 2.17E-05 |
| GO:0031098 | stress-activated protein kinase signaling cascade | 15/137 | 5.04E-09 | 2.20E-05 |
| GO:0006638 | neutral lipid metabolic process | 11/137 | 5.18E-09 | 2.27E-05 |
| GO:0006639 | acylglycerol metabolic process | 11/137 | 5.18E-09 | 2.27E-05 |
| GO:0098754 | detoxification | 11/137 | 5.18E-09 | 2.27E-05 |
| GO:0016051 | carbohydrate biosynthetic process | 13/137 | 5.24E-09 | 2.29E-05 |
| GO:0001659 | temperature homeostasis | 12/137 | 5.30E-09 | 2.32E-05 |
| GO:0006006 | glucose metabolic process | 13/137 | 5.55E-09 | 2.42E-05 |
| GO:0071887 | leukocyte apoptotic process | 10/137 | 6.19E-09 | 2.71E-05 |
| GO:0051651 | maintenance of location in cell | 13/137 | 6.21E-09 | 2.71E-05 |
| GO:0070873 | regulation of glycogen metabolic process | 7/137 | 6.39E-09 | 2.79E-05 |
| GO:1900182 | positive regulation of protein localization to nucleus | 9/137 | 6.46E-09 | 2.82E-05 |
| GO:0051480 | regulation of cytosolic calcium ion concentration | 16/137 | 7.18E-09 | 3.14E-05 |
| GO:0001541 | ovarian follicle development | 8/137 | 7.71E-09 | 3.37E-05 |
| GO:0016236 | macroautophagy | 15/137 | 7.85E-09 | 3.43E-05 |
| GO:0033157 | regulation of intracellular protein transport | 14/137 | 8.00E-09 | 3.50E-05 |
| GO:0050921 | positive regulation of chemotaxis | 11/137 | 8.12E-09 | 3.55E-05 |
| GO:0060326 | cell chemotaxis | 15/137 | 8.20E-09 | 3.58E-05 |
| GO:0002699 | positive regulation of immune effector process | 13/137 | 9.11E-09 | 3.98E-05 |
| GO:0010742 | macrophage derived foam cell differentiation | 7/137 | 9.55E-09 | 4.17E-05 |
| GO:0090077 | foam cell differentiation | 7/137 | 9.55E-09 | 4.17E-05 |
| GO:0033673 | negative regulation of kinase activity | 14/137 | 9.69E-09 | 4.23E-05 |
| GO:0046328 | regulation of JNK cascade | 12/137 | 9.96E-09 | 4.35E-05 |
| GO:0044409 | entry into host | 11/137 | 1.01E-08 | 4.41E-05 |
| GO:0045732 | positive regulation of protein catabolic process | 13/137 | 1.01E-08 | 4.43E-05 |
| GO:0002449 | lymphocyte mediated immunity | 16/137 | 1.02E-08 | 4.46E-05 |
| GO:0010466 | negative regulation of peptidase activity | 14/137 | 1.17E-08 | 5.11E-05 |
| GO:0002285 | lymphocyte activation involved in immune response | 12/137 | 1.19E-08 | 5.22E-05 |
| GO:0008217 | regulation of blood pressure | 12/137 | 1.19E-08 | 5.22E-05 |
| GO:0048639 | positive regulation of developmental growth | 12/137 | 1.19E-08 | 5.22E-05 |
| GO:0051043 | regulation of membrane protein ectodomain proteolysis | 6/137 | 1.20E-08 | 5.23E-05 |
| GO:0060149 | negative regulation of posttranscriptional gene silencing | 6/137 | 1.20E-08 | 5.23E-05 |
| GO:0060967 | negative regulation of gene silencing by RNA | 6/137 | 1.20E-08 | 5.23E-05 |
| GO:2001056 | positive regulation of cysteine-type endopeptidase activity | 11/137 | 1.34E-08 | 5.84E-05 |
| GO:0043534 | blood vessel endothelial cell migration | 12/137 | 1.35E-08 | 5.88E-05 |
| GO:0051341 | regulation of oxidoreductase activity | 10/137 | 1.35E-08 | 5.90E-05 |
| GO:1904019 | epithelial cell apoptotic process | 10/137 | 1.35E-08 | 5.90E-05 |
| GO:0022898 | regulation of transmembrane transporter activity | 14/137 | 1.41E-08 | 6.15E-05 |
| GO:1902950 | regulation of dendritic spine maintenance | 5/137 | 1.43E-08 | 6.23E-05 |
| GO:0030595 | leukocyte chemotaxis | 13/137 | 1.46E-08 | 6.39E-05 |
| GO:0035265 | organ growth | 12/137 | 1.51E-08 | 6.61E-05 |
| GO:0010676 | positive regulation of cellular carbohydrate metabolic process | 8/137 | 1.53E-08 | 6.69E-05 |
| GO:0034765 | regulation of ion transmembrane transport | 18/137 | 1.67E-08 | 7.29E-05 |

| | | | | |
|---|---|---|---|---|
| GO:0008625 | extrinsic apoptotic signaling pathway via death domain receptors | 9/137 | 1.68E-08 | 7.32E-05 |
| GO:0002793 | positive regulation of peptide secretion | 12/137 | 1.70E-08 | 7.43E-05 |
| GO:0071453 | cellular response to oxygen levels | 13/137 | 1.70E-08 | 7.44E-05 |
| GO:1904018 | positive regulation of vasculature development | 13/137 | 1.70E-08 | 7.44E-05 |
| GO:0060560 | developmental growth involved in morphogenesis | 13/137 | 1.79E-08 | 7.83E-05 |
| GO:0010906 | regulation of glucose metabolic process | 10/137 | 1.87E-08 | 8.17E-05 |
| GO:1900542 | regulation of purine nucleotide metabolic process | 10/137 | 1.87E-08 | 8.17E-05 |
| GO:0000187 | activation of MAPK activity | 11/137 | 1.88E-08 | 8.20E-05 |
| GO:0048588 | developmental cell growth | 13/137 | 1.88E-08 | 8.23E-05 |
| GO:0071214 | cellular response to abiotic stimulus | 15/137 | 1.89E-08 | 8.24E-05 |
| GO:0104004 | cellular response to environmental stimulus | 15/137 | 1.89E-08 | 8.24E-05 |
| GO:0051926 | negative regulation of calcium ion transport | 8/137 | 1.98E-08 | 8.65E-05 |
| GO:0006140 | regulation of nucleotide metabolic process | 10/137 | 2.19E-08 | 9.56E-05 |
| GO:0007249 | I-kappaB kinase/NF-kappaB signaling | 14/137 | 2.20E-08 | 9.63E-05 |
| GO:0010573 | vascular endothelial growth factor production | 8/137 | 2.24E-08 | 9.80E-05 |
| GO:0022604 | regulation of cell morphogenesis | 18/137 | 2.28E-08 | 9.94E-05 |
| GO:0032675 | regulation of interleukin-6 production | 11/137 | 2.29E-08 | 9.99E-05 |
| GO:0008631 | intrinsic apoptotic signaling pathway in response to oxidative stress | 7/137 | 2.36E-08 | 1.03E-04 |
| GO:0002688 | regulation of leukocyte chemotaxis | 10/137 | 2.37E-08 | 1.03E-04 |
| GO:1903578 | regulation of ATP metabolic process | 10/137 | 2.56E-08 | 1.12E-04 |
| GO:0009314 | response to radiation | 17/137 | 2.68E-08 | 1.17E-04 |
| GO:0034114 | regulation of heterotypic cell-cell adhesion | 6/137 | 2.68E-08 | 1.17E-04 |
| GO:1904035 | regulation of epithelial cell apoptotic process | 9/137 | 2.72E-08 | 1.19E-04 |
| GO:0034341 | response to interferon-gamma | 12/137 | 2.82E-08 | 1.23E-04 |
| GO:0006469 | negative regulation of protein kinase activity | 13/137 | 2.93E-08 | 1.28E-04 |
| GO:1900180 | regulation of protein localization to nucleus | 10/137 | 2.98E-08 | 1.30E-04 |
| GO:0016042 | lipid catabolic process | 15/137 | 3.02E-08 | 1.32E-04 |
| GO:0031960 | response to corticosteroid | 11/137 | 3.15E-08 | 1.38E-04 |
| GO:0043535 | regulation of blood vessel endothelial cell migration | 11/137 | 3.15E-08 | 1.38E-04 |
| GO:0051146 | striated muscle cell differentiation | 14/137 | 3.25E-08 | 1.42E-04 |
| GO:0045429 | positive regulation of nitric oxide biosynthetic process | 7/137 | 3.29E-08 | 1.44E-04 |
| GO:1902175 | regulation of oxidative stress-induced intrinsic apoptotic signaling pathway | 6/137 | 3.43E-08 | 1.50E-04 |
| GO:1903799 | negative regulation of production of miRNAs involved in gene silencing by miRNA | 5/137 | 3.56E-08 | 1.56E-04 |
| GO:1904062 | regulation of cation transmembrane transport | 15/137 | 3.80E-08 | 1.66E-04 |
| GO:0005978 | glycogen biosynthetic process | 7/137 | 3.86E-08 | 1.69E-04 |
| GO:0009250 | glucan biosynthetic process | 7/137 | 3.86E-08 | 1.69E-04 |
| GO:0010874 | regulation of cholesterol efflux | 7/137 | 3.86E-08 | 1.69E-04 |
| GO:0050798 | activated T cell proliferation | 7/137 | 3.86E-08 | 1.69E-04 |
| GO:1904407 | positive regulation of nitric oxide metabolic process | 7/137 | 3.86E-08 | 1.69E-04 |
| GO:0032374 | regulation of cholesterol transport | 8/137 | 4.07E-08 | 1.78E-04 |
| GO:0030217 | T cell differentiation | 13/137 | 4.08E-08 | 1.78E-04 |
| GO:0010632 | regulation of epithelial cell migration | 14/137 | 4.18E-08 | 1.83E-04 |
| GO:0019318 | hexose metabolic process | 13/137 | 4.27E-08 | 1.87E-04 |
| GO:0001101 | response to acid chemical | 10/137 | 4.31E-08 | 1.88E-04 |
| GO:0010575 | positive regulation of vascular endothelial growth factor production | 6/137 | 4.34E-08 | 1.90E-04 |
| GO:0030100 | regulation of endocytosis | 12/137 | 4.34E-08 | 1.90E-04 |
| GO:0006953 | acute-phase response | 7/137 | 4.51E-08 | 1.97E-04 |
| GO:0022602 | ovulation cycle process | 7/137 | 4.51E-08 | 1.97E-04 |
| GO:0048806 | genitalia development | 7/137 | 4.51E-08 | 1.97E-04 |
| GO:0032874 | positive regulation of stress-activated MAPK cascade | 11/137 | 4.57E-08 | 2.00E-04 |
| GO:0034614 | cellular response to reactive oxygen species | 11/137 | 4.57E-08 | 2.00E-04 |
| GO:0050679 | positive regulation of epithelial cell proliferation | 12/137 | 4.57E-08 | 2.00E-04 |
| GO:0045807 | positive regulation of endocytosis | 9/137 | 4.69E-08 | 2.05E-04 |
| GO:0007254 | JNK cascade | 12/137 | 5.07E-08 | 2.22E-04 |

| | | | | |
|---|---|---|---|---|
| GO:0010951 | negative regulation of endopeptidase activity | 13/137 | 5.13E-08 | 2.24E-04 |
| GO:1901617 | organic hydroxy compound biosynthetic process | 13/137 | 5.13E-08 | 2.24E-04 |
| GO:0070304 | positive regulation of stress-activated protein kinase signaling cascade | 11/137 | 5.15E-08 | 2.25E-04 |
| GO:0007263 | nitric oxide mediated signal transduction | 6/137 | 5.44E-08 | 2.38E-04 |
| GO:0006633 | fatty acid biosynthetic process | 11/137 | 5.47E-08 | 2.39E-04 |
| GO:0043281 | regulation of cysteine-type endopeptidase activity involved in apoptotic process | 12/137 | 5.62E-08 | 2.46E-04 |
| GO:0051966 | regulation of synaptic transmission; glutamatergic | 8/137 | 5.69E-08 | 2.49E-04 |
| GO:0010769 | regulation of cell morphogenesis involved in differentiation | 14/137 | 6.02E-08 | 2.63E-04 |
| GO:0008630 | intrinsic apoptotic signaling pathway in response to DNA damage | 9/137 | 6.07E-08 | 2.65E-04 |
| GO:0048709 | oligodendrocyte differentiation | 9/137 | 6.07E-08 | 2.65E-04 |
| GO:0030183 | B cell differentiation | 10/137 | 6.14E-08 | 2.68E-04 |
| GO:0006801 | superoxide metabolic process | 8/137 | 6.34E-08 | 2.77E-04 |
| GO:0016241 | regulation of macroautophagy | 11/137 | 6.52E-08 | 2.85E-04 |
| GO:0070588 | calcium ion transmembrane transport | 14/137 | 6.52E-08 | 2.85E-04 |
| GO:0007623 | circadian rhythm | 12/137 | 6.55E-08 | 2.86E-04 |
| GO:0032649 | regulation of interferon-gamma production | 9/137 | 6.61E-08 | 2.89E-04 |
| GO:0010631 | epithelial cell migration | 15/137 | 6.84E-08 | 2.99E-04 |
| GO:1903580 | positive regulation of ATP metabolic process | 7/137 | 7.03E-08 | 3.07E-04 |
| GO:0050805 | negative regulation of synaptic transmission | 8/137 | 7.05E-08 | 3.08E-04 |
| GO:0062014 | negative regulation of small molecule metabolic process | 9/137 | 7.18E-08 | 3.14E-04 |
| GO:0009411 | response to UV | 10/137 | 7.54E-08 | 3.30E-04 |
| GO:0097553 | calcium ion transmembrane import into cytosol | 10/137 | 7.54E-08 | 3.30E-04 |
| GO:0090132 | epithelium migration | 15/137 | 7.61E-08 | 3.33E-04 |
| GO:0034116 | positive regulation of heterotypic cell-cell adhesion | 5/137 | 7.69E-08 | 3.36E-04 |
| GO:0045725 | positive regulation of glycogen biosynthetic process | 5/137 | 7.69E-08 | 3.36E-04 |
| GO:0007409 | axonogenesis | 17/137 | 7.98E-08 | 3.49E-04 |
| GO:0019722 | calcium-mediated signaling | 12/137 | 7.99E-08 | 3.49E-04 |
| GO:0032355 | response to estradiol | 10/137 | 8.07E-08 | 3.52E-04 |
| GO:0071827 | plasma lipoprotein particle organization | 7/137 | 8.10E-08 | 3.54E-04 |
| GO:0032770 | positive regulation of monooxygenase activity | 6/137 | 8.33E-08 | 3.64E-04 |
| GO:0045940 | positive regulation of steroid metabolic process | 6/137 | 8.33E-08 | 3.64E-04 |
| GO:0016485 | protein processing | 12/137 | 8.40E-08 | 3.67E-04 |
| GO:0099175 | regulation of postsynapse organization | 9/137 | 8.46E-08 | 3.70E-04 |
| GO:0008361 | regulation of cell size | 11/137 | 8.68E-08 | 3.79E-04 |
| GO:0071346 | cellular response to interferon-gamma | 11/137 | 9.18E-08 | 4.01E-04 |
| GO:0010883 | regulation of lipid storage | 7/137 | 9.30E-08 | 4.06E-04 |
| GO:0090130 | tissue migration | 15/137 | 9.40E-08 | 4.11E-04 |
| GO:0000271 | polysaccharide biosynthetic process | 8/137 | 9.63E-08 | 4.21E-04 |
| GO:1903034 | regulation of response to wounding | 11/137 | 9.71E-08 | 4.24E-04 |
| GO:2001235 | positive regulation of apoptotic signaling pathway | 11/137 | 9.71E-08 | 4.24E-04 |
| GO:0044706 | multi-multicellular organism process | 12/137 | 9.72E-08 | 4.25E-04 |
| GO:0030099 | myeloid cell differentiation | 16/137 | 9.87E-08 | 4.31E-04 |
| GO:0010827 | regulation of glucose transmembrane transport | 8/137 | 1.06E-07 | 4.65E-04 |
| GO:0043536 | positive regulation of blood vessel endothelial cell migration | 8/137 | 1.06E-07 | 4.65E-04 |
| GO:0032731 | positive regulation of interleukin-1 beta production | 7/137 | 1.06E-07 | 4.65E-04 |
| GO:0002821 | positive regulation of adaptive immune response | 9/137 | 1.08E-07 | 4.70E-04 |
| GO:0070875 | positive regulation of glycogen metabolic process | 5/137 | 1.08E-07 | 4.73E-04 |
| GO:0050770 | regulation of axonogenesis | 11/137 | 1.15E-07 | 5.01E-04 |
| GO:0051147 | regulation of muscle cell differentiation | 11/137 | 1.15E-07 | 5.01E-04 |
| GO:0051149 | positive regulation of muscle cell differentiation | 9/137 | 1.16E-07 | 5.09E-04 |
| GO:1901224 | positive regulation of NIK/NF-kappaB signaling | 8/137 | 1.18E-07 | 5.14E-04 |
| GO:0051384 | response to glucocorticoid | 10/137 | 1.20E-07 | 5.22E-04 |
| GO:0015980 | energy derivation by oxidation of organic compounds | 13/137 | 1.23E-07 | 5.35E-04 |
| GO:0006909 | phagocytosis | 15/137 | 1.24E-07 | 5.40E-04 |
| GO:0048011 | neurotrophin TRK receptor signaling pathway | 6/137 | 1.24E-07 | 5.41E-04 |

| | | | | |
|---|---|---|---|---|
| GO:0071901 | negative regulation of protein serine/threonine kinase activity | 10/137 | 1.27E-07 | 5.57E-04 |
| GO:0001655 | urogenital system development | 14/137 | 1.30E-07 | 5.69E-04 |
| GO:0010565 | regulation of cellular ketone metabolic process | 11/137 | 1.35E-07 | 5.89E-04 |
| GO:0008347 | glial cell migration | 7/137 | 1.39E-07 | 6.05E-04 |
| GO:0051353 | positive regulation of oxidoreductase activity | 7/137 | 1.39E-07 | 6.05E-04 |
| GO:0071825 | protein-lipid complex subunit organization | 7/137 | 1.39E-07 | 6.05E-04 |
| GO:1902930 | regulation of alcohol biosynthetic process | 8/137 | 1.43E-07 | 6.24E-04 |
| GO:0042391 | regulation of membrane potential | 16/137 | 1.43E-07 | 6.26E-04 |
| GO:0150078 | positive regulation of neuroinflammatory response | 5/137 | 1.49E-07 | 6.51E-04 |
| GO:1990000 | amyloid fibril formation | 5/137 | 1.49E-07 | 6.51E-04 |
| GO:0048145 | regulation of fibroblast proliferation | 8/137 | 1.57E-07 | 6.86E-04 |
| GO:0032609 | interferon-gamma production | 9/137 | 1.58E-07 | 6.91E-04 |
| GO:0030900 | forebrain development | 15/137 | 1.67E-07 | 7.30E-04 |
| GO:0043542 | endothelial cell migration | 13/137 | 1.70E-07 | 7.43E-04 |
| GO:0010594 | regulation of endothelial cell migration | 12/137 | 1.70E-07 | 7.45E-04 |
| GO:0048144 | fibroblast proliferation | 8/137 | 1.73E-07 | 7.54E-04 |
| GO:0034390 | smooth muscle cell apoptotic process | 6/137 | 1.79E-07 | 7.84E-04 |
| GO:0034391 | regulation of smooth muscle cell apoptotic process | 6/137 | 1.79E-07 | 7.84E-04 |
| GO:0070884 | regulation of calcineurin-NFAT signaling cascade | 6/137 | 1.79E-07 | 7.84E-04 |
| GO:0018107 | peptidyl-threonine phosphorylation | 9/137 | 1.83E-07 | 8.01E-04 |
| GO:0010507 | negative regulation of autophagy | 8/137 | 1.89E-07 | 8.27E-04 |
| GO:0007565 | female pregnancy | 11/137 | 1.95E-07 | 8.51E-04 |
| GO:0007569 | cell aging | 9/137 | 1.97E-07 | 8.62E-04 |
| GO:0033559 | unsaturated fatty acid metabolic process | 9/137 | 1.97E-07 | 8.62E-04 |
| GO:0010976 | positive regulation of neuron projection development | 13/137 | 2.00E-07 | 8.72E-04 |
| GO:1900408 | negative regulation of cellular response to oxidative stress | 7/137 | 2.02E-07 | 8.81E-04 |
| GO:1903202 | negative regulation of oxidative stress-induced cell death | 7/137 | 2.02E-07 | 8.81E-04 |
| GO:0034637 | cellular carbohydrate biosynthetic process | 8/137 | 2.07E-07 | 9.07E-04 |
| GO:0097756 | negative regulation of blood vessel diameter | 8/137 | 2.07E-07 | 9.07E-04 |
| GO:0030902 | hindbrain development | 10/137 | 2.09E-07 | 9.12E-04 |
| GO:0106056 | regulation of calcineurin-mediated signaling | 6/137 | 2.14E-07 | 9.35E-04 |
| GO:1901099 | negative regulation of signal transduction in absence of ligand | 6/137 | 2.14E-07 | 9.35E-04 |
| GO:2001240 | negative regulation of extrinsic apoptotic signaling pathway in absence of ligand | 6/137 | 2.14E-07 | 9.35E-04 |
| GO:0048013 | ephrin receptor signaling pathway | 8/137 | 2.27E-07 | 9.92E-04 |
| GO:1904705 | regulation of vascular associated smooth muscle cell proliferation | 8/137 | 2.27E-07 | 9.92E-04 |
| GO:1990874 | vascular associated smooth muscle cell proliferation | 8/137 | 2.27E-07 | 9.92E-04 |
| GO:0006636 | unsaturated fatty acid biosynthetic process | 7/137 | 2.27E-07 | 9.93E-04 |
| GO:0034976 | response to endoplasmic reticulum stress | 13/137 | 2.34E-07 | 1.02E-03 |
| GO:0044262 | cellular carbohydrate metabolic process | 13/137 | 2.34E-07 | 1.02E-03 |
| GO:1901888 | regulation of cell junction assembly | 11/137 | 2.39E-07 | 1.04E-03 |
| GO:0060969 | negative regulation of gene silencing | 6/137 | 2.54E-07 | 1.11E-03 |
| GO:0032653 | regulation of interleukin-10 production | 7/137 | 2.56E-07 | 1.12E-03 |
| GO:0032732 | positive regulation of interleukin-1 production | 7/137 | 2.56E-07 | 1.12E-03 |
| GO:0061098 | positive regulation of protein tyrosine kinase activity | 7/137 | 2.56E-07 | 1.12E-03 |
| GO:0097755 | positive regulation of blood vessel diameter | 7/137 | 2.56E-07 | 1.12E-03 |
| GO:1902883 | negative regulation of response to oxidative stress | 7/137 | 2.56E-07 | 1.12E-03 |
| GO:1903428 | positive regulation of reactive oxygen species biosynthetic process | 7/137 | 2.56E-07 | 1.12E-03 |
| GO:0006690 | icosanoid metabolic process | 9/137 | 2.63E-07 | 1.15E-03 |
| GO:0010042 | response to manganese ion | 5/137 | 2.67E-07 | 1.16E-03 |
| GO:1904375 | regulation of protein localization to cell periphery | 9/137 | 2.82E-07 | 1.23E-03 |
| GO:0043954 | cellular component maintenance | 7/137 | 2.87E-07 | 1.26E-03 |
| GO:0002698 | negative regulation of immune effector process | 9/137 | 3.02E-07 | 1.32E-03 |
| GO:0090068 | positive regulation of cell cycle process | 13/137 | 3.17E-07 | 1.39E-03 |
| GO:0034103 | regulation of tissue remodeling | 8/137 | 3.22E-07 | 1.41E-03 |
| GO:0034605 | cellular response to heat | 9/137 | 3.23E-07 | 1.41E-03 |

| | | | | |
|---|---|---|---|---|
| GO:0043500 | muscle adaptation | 9/137 | 3.23E-07 | 1.41E-03 |
| GO:0046165 | alcohol biosynthetic process | 10/137 | 3.33E-07 | 1.45E-03 |
| GO:0018210 | peptidyl-threonine modification | 9/137 | 3.46E-07 | 1.51E-03 |
| GO:0032386 | regulation of intracellular transport | 14/137 | 3.51E-07 | 1.53E-03 |
| GO:1903321 | negative regulation of protein modification by small protein conjugation or removal | 8/137 | 3.51E-07 | 1.53E-03 |
| GO:0038179 | neurotrophin signaling pathway | 6/137 | 3.52E-07 | 1.54E-03 |
| GO:0055090 | acylglycerol homeostasis | 6/137 | 3.52E-07 | 1.54E-03 |
| GO:0070328 | triglyceride homeostasis | 6/137 | 3.52E-07 | 1.54E-03 |
| GO:0045766 | positive regulation of angiogenesis | 11/137 | 3.54E-07 | 1.55E-03 |
| GO:0032613 | interleukin-10 production | 7/137 | 3.60E-07 | 1.57E-03 |
| GO:0009612 | response to mechanical stimulus | 11/137 | 3.71E-07 | 1.62E-03 |
| GO:1903531 | negative regulation of secretion by cell | 10/137 | 3.72E-07 | 1.63E-03 |
| GO:0048872 | homeostasis of number of cells | 12/137 | 3.73E-07 | 1.63E-03 |
| GO:0090257 | regulation of muscle system process | 12/137 | 3.73E-07 | 1.63E-03 |
| GO:0070555 | response to interleukin-1 | 11/137 | 4.08E-07 | 1.78E-03 |
| GO:1902895 | positive regulation of pri-miRNA transcription by RNA polymerase II | 6/137 | 4.12E-07 | 1.80E-03 |
| GO:0010660 | regulation of muscle cell apoptotic process | 8/137 | 4.15E-07 | 1.81E-03 |
| GO:0002576 | platelet degranulation | 9/137 | 4.23E-07 | 1.85E-03 |
| GO:0021537 | telencephalon development | 12/137 | 4.23E-07 | 1.85E-03 |
| GO:0001667 | ameboidal-type cell migration | 16/137 | 4.33E-07 | 1.89E-03 |
| GO:0051100 | negative regulation of binding | 10/137 | 4.39E-07 | 1.92E-03 |
| GO:0042180 | cellular ketone metabolic process | 12/137 | 4.41E-07 | 1.93E-03 |
| GO:0038083 | peptidyl-tyrosine autophosphorylation | 6/137 | 4.79E-07 | 2.09E-03 |
| GO:0002690 | positive regulation of leukocyte chemotaxis | 8/137 | 4.88E-07 | 2.13E-03 |
| GO:0035249 | synaptic transmission; glutamatergic | 8/137 | 4.88E-07 | 2.13E-03 |
| GO:0007584 | response to nutrient | 10/137 | 4.90E-07 | 2.14E-03 |
| GO:0046824 | positive regulation of nucleocytoplasmic transport | 7/137 | 4.97E-07 | 2.17E-03 |
| GO:2000378 | negative regulation of reactive oxygen species metabolic process | 7/137 | 4.97E-07 | 2.17E-03 |
| GO:0010595 | positive regulation of endothelial cell migration | 9/137 | 5.14E-07 | 2.25E-03 |
| GO:0042100 | B cell proliferation | 8/137 | 5.29E-07 | 2.31E-03 |
| GO:0031330 | negative regulation of cellular catabolic process | 12/137 | 5.40E-07 | 2.36E-03 |
| GO:0001660 | fever generation | 4/137 | 5.40E-07 | 2.36E-03 |
| GO:0002676 | regulation of chronic inflammatory response | 4/137 | 5.40E-07 | 2.36E-03 |
| GO:0045348 | positive regulation of MHC class II biosynthetic process | 4/137 | 5.40E-07 | 2.36E-03 |
| GO:1990535 | neuron projection maintenance | 4/137 | 5.40E-07 | 2.36E-03 |
| GO:2000425 | regulation of apoptotic cell clearance | 4/137 | 5.40E-07 | 2.36E-03 |
| GO:0099173 | postsynapse organization | 10/137 | 5.45E-07 | 2.38E-03 |
| GO:0010907 | positive regulation of glucose metabolic process | 6/137 | 5.56E-07 | 2.43E-03 |
| GO:0021795 | cerebral cortex cell migration | 6/137 | 5.56E-07 | 2.43E-03 |
| GO:0046006 | regulation of activated T cell proliferation | 6/137 | 5.56E-07 | 2.43E-03 |
| GO:0090279 | regulation of calcium ion import | 6/137 | 5.56E-07 | 2.43E-03 |
| GO:0071404 | cellular response to low-density lipoprotein particle stimulus | 5/137 | 5.68E-07 | 2.48E-03 |
| GO:1900273 | positive regulation of long-term synaptic potentiation | 5/137 | 5.68E-07 | 2.48E-03 |
| GO:1903798 | regulation of production of miRNAs involved in gene silencing by miRNA | 5/137 | 5.68E-07 | 2.48E-03 |
| GO:0010657 | muscle cell apoptotic process | 8/137 | 5.72E-07 | 2.50E-03 |
| GO:0045833 | negative regulation of lipid metabolic process | 8/137 | 5.72E-07 | 2.50E-03 |
| GO:0043280 | positive regulation of cysteine-type endopeptidase activity involved in apoptotic process | 9/137 | 5.84E-07 | 2.55E-03 |
| GO:0046718 | viral entry into host cell | 9/137 | 5.84E-07 | 2.55E-03 |
| GO:0009416 | response to light stimulus | 13/137 | 5.91E-07 | 2.58E-03 |
| GO:0002637 | regulation of immunoglobulin production | 7/137 | 6.12E-07 | 2.67E-03 |
| GO:0034381 | plasma lipoprotein particle clearance | 7/137 | 6.12E-07 | 2.67E-03 |
| GO:0010634 | positive regulation of epithelial cell migration | 10/137 | 6.38E-07 | 2.79E-03 |
| GO:0002369 | T cell cytokine production | 6/137 | 6.42E-07 | 2.81E-03 |
| GO:0006959 | humoral immune response | 14/137 | 6.52E-07 | 2.85E-03 |

| | | | | |
|---|---|---|---|---|
| GO:0002702 | positive regulation of production of molecular mediator of immune response | 8/137 | 6.68E-07 | 2.92E-03 |
| GO:0042698 | ovulation cycle | 7/137 | 6.77E-07 | 2.96E-03 |
| GO:0002719 | negative regulation of cytokine production involved in immune response | 5/137 | 7.14E-07 | 3.12E-03 |
| GO:0045821 | positive regulation of glycolytic process | 5/137 | 7.14E-07 | 3.12E-03 |
| GO:0050995 | negative regulation of lipid catabolic process | 5/137 | 7.14E-07 | 3.12E-03 |
| GO:0045666 | positive regulation of neuron differentiation | 14/137 | 7.16E-07 | 3.13E-03 |
| GO:0007215 | glutamate receptor signaling pathway | 8/137 | 7.21E-07 | 3.15E-03 |
| GO:0010522 | regulation of calcium ion transport into cytosol | 8/137 | 7.21E-07 | 3.15E-03 |
| GO:0032637 | interleukin-8 production | 8/137 | 7.21E-07 | 3.15E-03 |
| GO:0033173 | calcineurin-NFAT signaling cascade | 6/137 | 7.39E-07 | 3.23E-03 |
| GO:0002705 | positive regulation of leukocyte mediated immunity | 9/137 | 7.48E-07 | 3.27E-03 |
| GO:0046330 | positive regulation of JNK cascade | 9/137 | 7.48E-07 | 3.27E-03 |
| GO:1903076 | regulation of protein localization to plasma membrane | 8/137 | 7.78E-07 | 3.40E-03 |
| GO:0042113 | B cell activation | 13/137 | 8.08E-07 | 3.53E-03 |
| GO:0033692 | cellular polysaccharide biosynthetic process | 7/137 | 8.24E-07 | 3.60E-03 |
| GO:0031349 | positive regulation of defense response | 14/137 | 8.37E-07 | 3.66E-03 |
| GO:0034372 | very-low-density lipoprotein particle remodeling | 4/137 | 8.44E-07 | 3.69E-03 |
| GO:0010828 | positive regulation of glucose transmembrane transport | 6/137 | 8.48E-07 | 3.70E-03 |
| GO:0034504 | protein localization to nucleus | 12/137 | 8.63E-07 | 3.77E-03 |
| GO:0070920 | regulation of production of small RNA involved in gene silencing by RNA | 5/137 | 8.87E-07 | 3.88E-03 |
| GO:0002456 | T cell mediated immunity | 8/137 | 9.03E-07 | 3.94E-03 |
| GO:0045600 | positive regulation of fat cell differentiation | 7/137 | 9.08E-07 | 3.97E-03 |
| GO:0061028 | establishment of endothelial barrier | 6/137 | 9.69E-07 | 4.24E-03 |
| GO:0002824 | positive regulation of adaptive immune response based on somatic recombination of immune receptors built from immunoglobulin superfamily domains | 8/137 | 9.71E-07 | 4.24E-03 |
| GO:0044070 | regulation of anion transport | 8/137 | 9.71E-07 | 4.24E-03 |
| GO:0005977 | glycogen metabolic process | 7/137 | 9.98E-07 | 4.36E-03 |
| GO:0010517 | regulation of phospholipase activity | 7/137 | 9.98E-07 | 4.36E-03 |
| GO:0008286 | insulin receptor signaling pathway | 9/137 | 1.01E-06 | 4.41E-03 |
| GO:0002708 | positive regulation of lymphocyte mediated immunity | 8/137 | 1.04E-06 | 4.56E-03 |
| GO:0009743 | response to carbohydrate | 11/137 | 1.09E-06 | 4.76E-03 |
| GO:0006073 | cellular glucan metabolic process | 7/137 | 1.10E-06 | 4.79E-03 |
| GO:0044042 | glucan metabolic process | 7/137 | 1.10E-06 | 4.79E-03 |
| GO:0031346 | positive regulation of cell projection organization | 14/137 | 1.10E-06 | 4.81E-03 |
| GO:0010559 | regulation of glycoprotein biosynthetic process | 6/137 | 1.10E-06 | 4.83E-03 |
| GO:0030225 | macrophage differentiation | 6/137 | 1.10E-06 | 4.83E-03 |
| GO:0061138 | morphogenesis of a branching epithelium | 10/137 | 1.11E-06 | 4.86E-03 |
| GO:0007050 | cell cycle arrest | 11/137 | 1.14E-06 | 4.96E-03 |
| GO:0043123 | positive regulation of I-kappaB kinase/NF-kappaB signaling | 10/137 | 1.17E-06 | 5.10E-03 |
| GO:0031100 | animal organ regeneration | 7/137 | 1.20E-06 | 5.25E-03 |
| GO:0042542 | response to hydrogen peroxide | 9/137 | 1.20E-06 | 5.25E-03 |
| GO:0046631 | alpha-beta T cell activation | 9/137 | 1.20E-06 | 5.25E-03 |
| GO:1990778 | protein localization to cell periphery | 13/137 | 1.21E-06 | 5.28E-03 |
| GO:2001239 | regulation of extrinsic apoptotic signaling pathway in absence of ligand | 6/137 | 1.25E-06 | 5.48E-03 |
| GO:0021819 | layer formation in cerebral cortex | 4/137 | 1.26E-06 | 5.50E-03 |
| GO:0033700 | phospholipid efflux | 4/137 | 1.26E-06 | 5.50E-03 |
| GO:1902337 | regulation of apoptotic process involved in morphogenesis | 4/137 | 1.26E-06 | 5.50E-03 |
| GO:0051048 | negative regulation of secretion | 10/137 | 1.28E-06 | 5.61E-03 |
| GO:0002286 | T cell activation involved in immune response | 8/137 | 1.29E-06 | 5.64E-03 |
| GO:0033077 | T cell differentiation in thymus | 7/137 | 1.31E-06 | 5.75E-03 |
| GO:0033555 | multicellular organismal response to stress | 7/137 | 1.31E-06 | 5.75E-03 |
| GO:1904659 | glucose transmembrane transport | 8/137 | 1.38E-06 | 6.05E-03 |
| GO:0006692 | prostanoid metabolic process | 6/137 | 1.42E-06 | 6.21E-03 |
| GO:0006693 | prostaglandin metabolic process | 6/137 | 1.42E-06 | 6.21E-03 |

| | | | | |
|---|---|---|---|---|
| GO:0097720 | calcineurin-mediated signaling | 6/137 | 1.42E-06 | 6.21E-03 |
| GO:0007422 | peripheral nervous system development | 7/137 | 1.44E-06 | 6.28E-03 |
| GO:0061387 | regulation of extent of cell growth | 8/137 | 1.48E-06 | 6.48E-03 |
| GO:0051604 | protein maturation | 12/137 | 1.55E-06 | 6.79E-03 |
| GO:0042310 | vasoconstriction | 7/137 | 1.57E-06 | 6.86E-03 |
| GO:0060998 | regulation of dendritic spine development | 7/137 | 1.57E-06 | 6.86E-03 |
| GO:0003012 | muscle system process | 15/137 | 1.57E-06 | 6.87E-03 |
| GO:0005976 | polysaccharide metabolic process | 8/137 | 1.59E-06 | 6.94E-03 |
| GO:0046822 | regulation of nucleocytoplasmic transport | 8/137 | 1.59E-06 | 6.94E-03 |
| GO:0061041 | regulation of wound healing | 9/137 | 1.59E-06 | 6.95E-03 |
| GO:0097300 | programmed necrotic cell death | 6/137 | 1.61E-06 | 7.02E-03 |
| GO:1902932 | positive regulation of alcohol biosynthetic process | 5/137 | 1.61E-06 | 7.05E-03 |
| GO:0030336 | negative regulation of cell migration | 13/137 | 1.67E-06 | 7.29E-03 |
| GO:1903169 | regulation of calcium ion transmembrane transport | 9/137 | 1.68E-06 | 7.35E-03 |
| GO:0001938 | positive regulation of endothelial cell proliferation | 8/137 | 1.70E-06 | 7.42E-03 |
| GO:0002674 | negative regulation of acute inflammatory response | 4/137 | 1.81E-06 | 7.90E-03 |
| GO:0034370 | triglyceride-rich lipoprotein particle remodeling | 4/137 | 1.81E-06 | 7.90E-03 |
| GO:0071287 | cellular response to manganese ion | 4/137 | 1.81E-06 | 7.90E-03 |
| GO:1900272 | negative regulation of long-term synaptic potentiation | 4/137 | 1.81E-06 | 7.90E-03 |
| GO:0006984 | ER-nucleus signaling pathway | 6/137 | 1.81E-06 | 7.90E-03 |
| GO:1902893 | regulation of pri-miRNA transcription by RNA polymerase II | 6/137 | 1.81E-06 | 7.90E-03 |
| GO:0008645 | hexose transmembrane transport | 8/137 | 1.81E-06 | 7.93E-03 |
| GO:0022037 | metencephalon development | 8/137 | 1.81E-06 | 7.93E-03 |
| GO:0043200 | response to amino acid | 8/137 | 1.81E-06 | 7.93E-03 |
| GO:0072330 | monocarboxylic acid biosynthetic process | 11/137 | 1.85E-06 | 8.08E-03 |
| GO:0001503 | ossification | 14/137 | 1.86E-06 | 8.13E-03 |
| GO:0031397 | negative regulation of protein ubiquitination | 7/137 | 1.86E-06 | 8.15E-03 |
| GO:0048678 | response to axon injury | 7/137 | 1.86E-06 | 8.15E-03 |
| GO:0016358 | dendrite development | 11/137 | 1.92E-06 | 8.41E-03 |
| GO:2000108 | positive regulation of leukocyte apoptotic process | 5/137 | 1.94E-06 | 8.47E-03 |
| GO:0001937 | negative regulation of endothelial cell proliferation | 7/137 | 2.03E-06 | 8.87E-03 |
| GO:0034644 | cellular response to UV | 7/137 | 2.03E-06 | 8.87E-03 |
| GO:0007162 | negative regulation of cell adhesion | 12/137 | 2.06E-06 | 8.99E-03 |
| GO:0015749 | monosaccharide transmembrane transport | 8/137 | 2.07E-06 | 9.05E-03 |
| GO:0032640 | tumor necrosis factor production | 8/137 | 2.07E-06 | 9.05E-03 |
| GO:0046620 | regulation of organ growth | 8/137 | 2.07E-06 | 9.05E-03 |
| GO:0001890 | placenta development | 9/137 | 2.09E-06 | 9.11E-03 |
| GO:0001763 | morphogenesis of a branching structure | 10/137 | 2.14E-06 | 9.33E-03 |
| GO:0031099 | regeneration | 10/137 | 2.14E-06 | 9.33E-03 |
| GO:0001818 | negative regulation of cytokine production | 13/137 | 2.28E-06 | 9.94E-03 |
| GO:0061614 | pri-miRNA transcription by RNA polymerase II | 6/137 | 2.28E-06 | 9.96E-03 |
| GO:0043457 | regulation of cellular respiration | 5/137 | 2.31E-06 | 1.01E-02 |
| GO:0090314 | positive regulation of protein targeting to membrane | 5/137 | 2.31E-06 | 1.01E-02 |
| GO:0034219 | carbohydrate transmembrane transport | 8/137 | 2.35E-06 | 1.03E-02 |
| GO:1902107 | positive regulation of leukocyte differentiation | 9/137 | 2.44E-06 | 1.07E-02 |
| GO:0071706 | tumor necrosis factor superfamily cytokine production | 8/137 | 2.51E-06 | 1.10E-02 |
| GO:1901222 | regulation of NIK/NF-kappaB signaling | 8/137 | 2.51E-06 | 1.10E-02 |
| GO:0050930 | induction of positive chemotaxis | 4/137 | 2.52E-06 | 1.10E-02 |
| GO:0050966 | detection of mechanical stimulus involved in sensory perception of pain | 4/137 | 2.52E-06 | 1.10E-02 |
| GO:1902430 | negative regulation of amyloid-beta formation | 4/137 | 2.52E-06 | 1.10E-02 |
| GO:1904748 | regulation of apoptotic process involved in development | 4/137 | 2.52E-06 | 1.10E-02 |
| GO:2000027 | regulation of animal organ morphogenesis | 11/137 | 2.52E-06 | 1.10E-02 |
| GO:1903018 | regulation of glycoprotein metabolic process | 6/137 | 2.55E-06 | 1.11E-02 |
| GO:2000146 | negative regulation of cell motility | 13/137 | 2.65E-06 | 1.16E-02 |
| GO:0042594 | response to starvation | 10/137 | 2.66E-06 | 1.16E-02 |
| GO:0034767 | positive regulation of ion transmembrane transport | 9/137 | 2.71E-06 | 1.18E-02 |
| GO:0002675 | positive regulation of acute inflammatory response | 5/137 | 2.74E-06 | 1.20E-02 |

| | | | | |
|---|---|---|---|---|
| GO:0034368 | protein-lipid complex remodeling | 5/137 | 2.74E-06 | 1.20E-02 |
| GO:0034369 | plasma lipoprotein particle remodeling | 5/137 | 2.74E-06 | 1.20E-02 |
| GO:0034377 | plasma lipoprotein particle assembly | 5/137 | 2.74E-06 | 1.20E-02 |
| GO:0034284 | response to monosaccharide | 10/137 | 2.78E-06 | 1.22E-02 |
| GO:0090287 | regulation of cellular response to growth factor stimulus | 12/137 | 2.79E-06 | 1.22E-02 |
| GO:0045844 | positive regulation of striated muscle tissue development | 7/137 | 2.81E-06 | 1.23E-02 |
| GO:0048636 | positive regulation of muscle organ development | 7/137 | 2.81E-06 | 1.23E-02 |
| GO:0070509 | calcium ion import | 7/137 | 2.81E-06 | 1.23E-02 |
| GO:0022029 | telencephalon cell migration | 6/137 | 2.84E-06 | 1.24E-02 |
| GO:0046034 | ATP metabolic process | 12/137 | 2.88E-06 | 1.26E-02 |
| GO:0071456 | cellular response to hypoxia | 10/137 | 2.90E-06 | 1.27E-02 |
| GO:0009267 | cellular response to starvation | 9/137 | 3.00E-06 | 1.31E-02 |
| GO:0001776 | leukocyte homeostasis | 7/137 | 3.04E-06 | 1.33E-02 |
| GO:0006112 | energy reserve metabolic process | 7/137 | 3.04E-06 | 1.33E-02 |
| GO:0006919 | activation of cysteine-type endopeptidase activity involved in apoptotic process | 7/137 | 3.04E-06 | 1.33E-02 |
| GO:1901863 | positive regulation of muscle tissue development | 7/137 | 3.04E-06 | 1.33E-02 |
| GO:0001764 | neuron migration | 9/137 | 3.15E-06 | 1.38E-02 |
| GO:0046661 | male sex differentiation | 9/137 | 3.15E-06 | 1.38E-02 |
| GO:0002712 | regulation of B cell mediated immunity | 6/137 | 3.17E-06 | 1.38E-02 |
| GO:0002889 | regulation of immunoglobulin mediated immune response | 6/137 | 3.17E-06 | 1.38E-02 |
| GO:0002761 | regulation of myeloid leukocyte differentiation | 8/137 | 3.22E-06 | 1.41E-02 |
| GO:0048675 | axon extension | 8/137 | 3.22E-06 | 1.41E-02 |
| GO:0002724 | regulation of T cell cytokine production | 5/137 | 3.23E-06 | 1.41E-02 |
| GO:0010743 | regulation of macrophage derived foam cell differentiation | 5/137 | 3.23E-06 | 1.41E-02 |
| GO:0034367 | protein-containing complex remodeling | 5/137 | 3.23E-06 | 1.41E-02 |
| GO:0050849 | negative regulation of calcium-mediated signaling | 5/137 | 3.23E-06 | 1.41E-02 |
| GO:0001894 | tissue homeostasis | 11/137 | 3.28E-06 | 1.43E-02 |
| GO:0034329 | cell junction assembly | 14/137 | 3.40E-06 | 1.49E-02 |
| GO:0046486 | glycerolipid metabolic process | 14/137 | 3.40E-06 | 1.49E-02 |
| GO:0034349 | glial cell apoptotic process | 4/137 | 3.41E-06 | 1.49E-02 |
| GO:0045346 | regulation of MHC class II biosynthetic process | 4/137 | 3.41E-06 | 1.49E-02 |
| GO:0051044 | positive regulation of membrane protein ectodomain proteolysis | 4/137 | 3.41E-06 | 1.49E-02 |
| GO:0070885 | negative regulation of calcineurin-NFAT signaling cascade | 4/137 | 3.41E-06 | 1.49E-02 |
| GO:0106057 | negative regulation of calcineurin-mediated signaling | 4/137 | 3.41E-06 | 1.49E-02 |
| GO:1901550 | regulation of endothelial cell development | 4/137 | 3.41E-06 | 1.49E-02 |
| GO:1903140 | regulation of establishment of endothelial barrier | 4/137 | 3.41E-06 | 1.49E-02 |
| GO:0051101 | regulation of DNA binding | 8/137 | 3.42E-06 | 1.49E-02 |
| GO:0030278 | regulation of ossification | 10/137 | 3.44E-06 | 1.50E-02 |
| GO:0048016 | inositol phosphate-mediated signaling | 6/137 | 3.52E-06 | 1.54E-02 |
| GO:0090276 | regulation of peptide hormone secretion | 10/137 | 3.59E-06 | 1.57E-02 |
| GO:0008637 | apoptotic mitochondrial changes | 8/137 | 3.63E-06 | 1.59E-02 |
| GO:0033028 | myeloid cell apoptotic process | 5/137 | 3.78E-06 | 1.65E-02 |
| GO:0032412 | regulation of ion transmembrane transporter activity | 11/137 | 3.79E-06 | 1.66E-02 |
| GO:0043506 | regulation of JUN kinase activity | 7/137 | 3.83E-06 | 1.67E-02 |
| GO:0045931 | positive regulation of mitotic cell cycle | 9/137 | 3.84E-06 | 1.68E-02 |
| GO:0007566 | embryo implantation | 6/137 | 3.90E-06 | 1.70E-02 |
| GO:0021885 | forebrain cell migration | 6/137 | 3.90E-06 | 1.70E-02 |
| GO:0046883 | regulation of hormone secretion | 11/137 | 4.08E-06 | 1.78E-02 |
| GO:0043030 | regulation of macrophage activation | 6/137 | 4.32E-06 | 1.89E-02 |
| GO:0046456 | icosanoid biosynthetic process | 6/137 | 4.32E-06 | 1.89E-02 |
| GO:0050707 | regulation of cytokine secretion | 6/137 | 4.32E-06 | 1.89E-02 |
| GO:0034763 | negative regulation of transmembrane transport | 8/137 | 4.33E-06 | 1.89E-02 |
| GO:0007616 | long-term memory | 5/137 | 4.41E-06 | 1.93E-02 |
| GO:0016242 | negative regulation of macroautophagy | 5/137 | 4.41E-06 | 1.93E-02 |
| GO:0042311 | vasodilation | 5/137 | 4.41E-06 | 1.93E-02 |
| GO:0042759 | long-chain fatty acid biosynthetic process | 5/137 | 4.41E-06 | 1.93E-02 |
| GO:0043276 | anoikis | 5/137 | 4.41E-06 | 1.93E-02 |

| | | | | |
|---|---|---|---|---|
| GO:0045907 | positive regulation of vasoconstriction | 5/137 | 4.41E-06 | 1.93E-02 |
| GO:0050715 | positive regulation of cytokine secretion | 5/137 | 4.41E-06 | 1.93E-02 |
| GO:0036294 | cellular response to decreased oxygen levels | 10/137 | 4.41E-06 | 1.93E-02 |
| GO:0032535 | regulation of cellular component size | 13/137 | 4.48E-06 | 1.96E-02 |
| GO:0017014 | protein nitrosylation | 4/137 | 4.53E-06 | 1.98E-02 |
| GO:0018119 | peptidyl-cysteine S-nitrosylation | 4/137 | 4.53E-06 | 1.98E-02 |
| GO:0045342 | MHC class II biosynthetic process | 4/137 | 4.53E-06 | 1.98E-02 |
| GO:1990138 | neuron projection extension | 9/137 | 4.66E-06 | 2.03E-02 |
| GO:0001836 | release of cytochrome c from mitochondria | 6/137 | 4.77E-06 | 2.08E-02 |
| GO:0019229 | regulation of vasoconstriction | 6/137 | 4.77E-06 | 2.08E-02 |
| GO:0033209 | tumor necrosis factor-mediated signaling pathway | 9/137 | 4.88E-06 | 2.13E-02 |
| GO:0002064 | epithelial cell development | 10/137 | 4.98E-06 | 2.18E-02 |
| GO:0065005 | protein-lipid complex assembly | 5/137 | 5.11E-06 | 2.23E-02 |
| GO:1901030 | positive regulation of mitochondrial outer membrane permeabilization involved in apoptotic signaling pathway | 5/137 | 5.11E-06 | 2.23E-02 |
| GO:0010469 | regulation of signaling receptor activity | 9/137 | 5.12E-06 | 2.24E-02 |
| GO:0032677 | regulation of interleukin-8 production | 7/137 | 5.14E-06 | 2.25E-02 |
| GO:0097529 | myeloid leukocyte migration | 10/137 | 5.19E-06 | 2.27E-02 |
| GO:0010574 | regulation of vascular endothelial growth factor production | 6/137 | 5.25E-06 | 2.30E-02 |
| GO:0014706 | striated muscle tissue development | 13/137 | 5.30E-06 | 2.32E-02 |
| GO:0050680 | negative regulation of epithelial cell proliferation | 9/137 | 5.36E-06 | 2.34E-02 |
| GO:1901568 | fatty acid derivative metabolic process | 9/137 | 5.36E-06 | 2.34E-02 |
| GO:0060562 | epithelial tube morphogenesis | 12/137 | 5.46E-06 | 2.39E-02 |
| GO:0030888 | regulation of B cell proliferation | 6/137 | 5.78E-06 | 2.53E-02 |
| GO:0034113 | heterotypic cell-cell adhesion | 6/137 | 5.78E-06 | 2.53E-02 |
| GO:0055081 | anion homeostasis | 6/137 | 5.78E-06 | 2.53E-02 |
| GO:2001244 | positive regulation of intrinsic apoptotic signaling pathway | 6/137 | 5.78E-06 | 2.53E-02 |
| GO:0045665 | negative regulation of neuron differentiation | 10/137 | 5.84E-06 | 2.55E-02 |
| GO:0030540 | female genitalia development | 4/137 | 5.89E-06 | 2.57E-02 |
| GO:0030730 | sequestering of triglyceride | 4/137 | 5.89E-06 | 2.57E-02 |
| GO:0031649 | heat generation | 4/137 | 5.89E-06 | 2.57E-02 |
| GO:1902992 | negative regulation of amyloid precursor protein catabolic process | 4/137 | 5.89E-06 | 2.57E-02 |
| GO:0002701 | negative regulation of production of molecular mediator of immune response | 5/137 | 5.90E-06 | 2.58E-02 |
| GO:0090313 | regulation of protein targeting to membrane | 5/137 | 5.90E-06 | 2.58E-02 |
| GO:1903725 | regulation of phospholipid metabolic process | 7/137 | 5.93E-06 | 2.59E-02 |
| GO:0030198 | extracellular matrix organization | 13/137 | 6.25E-06 | 2.73E-02 |
| GO:0043062 | extracellular structure organization | 13/137 | 6.42E-06 | 2.81E-02 |
| GO:0040013 | negative regulation of locomotion | 13/137 | 6.60E-06 | 2.88E-02 |
| GO:0072659 | protein localization to plasma membrane | 11/137 | 6.63E-06 | 2.90E-02 |
| GO:0090322 | regulation of superoxide metabolic process | 5/137 | 6.79E-06 | 2.97E-02 |
| GO:0030516 | regulation of axon extension | 7/137 | 6.81E-06 | 2.97E-02 |
| GO:0071674 | mononuclear cell migration | 7/137 | 6.81E-06 | 2.97E-02 |
| GO:0001885 | endothelial cell development | 6/137 | 6.97E-06 | 3.05E-02 |
| GO:0046686 | response to cadmium ion | 6/137 | 6.97E-06 | 3.05E-02 |
| GO:0060135 | maternal process involved in female pregnancy | 6/137 | 6.97E-06 | 3.05E-02 |
| GO:0070265 | necrotic cell death | 6/137 | 6.97E-06 | 3.05E-02 |
| GO:1900449 | regulation of glutamate receptor signaling pathway | 6/137 | 6.97E-06 | 3.05E-02 |
| GO:2000401 | regulation of lymphocyte migration | 6/137 | 6.97E-06 | 3.05E-02 |
| GO:0001822 | kidney development | 11/137 | 7.09E-06 | 3.10E-02 |
| GO:0051271 | negative regulation of cellular component movement | 13/137 | 7.16E-06 | 3.13E-02 |
| GO:0030316 | osteoclast differentiation | 7/137 | 7.29E-06 | 3.18E-02 |
| GO:0050810 | regulation of steroid biosynthetic process | 7/137 | 7.29E-06 | 3.18E-02 |
| GO:2001243 | negative regulation of intrinsic apoptotic signaling pathway | 7/137 | 7.29E-06 | 3.18E-02 |
| GO:0043903 | regulation of symbiotic process | 10/137 | 7.36E-06 | 3.22E-02 |
| GO:0034375 | high-density lipoprotein particle remodeling | 4/137 | 7.52E-06 | 3.29E-02 |
| GO:0034433 | steroid esterification | 4/137 | 7.52E-06 | 3.29E-02 |
| GO:0034434 | sterol esterification | 4/137 | 7.52E-06 | 3.29E-02 |

| | | | | |
|---|---|---|---|---|
| GO:0034435 | cholesterol esterification | 4/137 | 7.52E-06 | 3.29E-02 |
| GO:1900221 | regulation of amyloid-beta clearance | 4/137 | 7.52E-06 | 3.29E-02 |
| GO:0032729 | positive regulation of interferon-gamma production | 6/137 | 7.63E-06 | 3.34E-02 |
| GO:1905953 | negative regulation of lipid localization | 6/137 | 7.63E-06 | 3.34E-02 |
| GO:0071347 | cellular response to interleukin-1 | 9/137 | 7.71E-06 | 3.37E-02 |
| GO:0032733 | positive regulation of interleukin-10 production | 5/137 | 7.77E-06 | 3.40E-02 |
| GO:0043029 | T cell homeostasis | 5/137 | 7.77E-06 | 3.40E-02 |
| GO:0045923 | positive regulation of fatty acid metabolic process | 5/137 | 7.77E-06 | 3.40E-02 |
| GO:0048009 | insulin-like growth factor receptor signaling pathway | 5/137 | 7.77E-06 | 3.40E-02 |
| GO:0035304 | regulation of protein dephosphorylation | 8/137 | 7.98E-06 | 3.49E-02 |
| GO:0038061 | NIK/NF-kappaB signaling | 9/137 | 8.05E-06 | 3.52E-02 |
| GO:0021700 | developmental maturation | 11/137 | 8.10E-06 | 3.54E-02 |
| GO:2001257 | regulation of cation channel activity | 9/137 | 8.41E-06 | 3.68E-02 |
| GO:0001505 | regulation of neurotransmitter levels | 10/137 | 8.56E-06 | 3.74E-02 |
| GO:0071695 | anatomical structure maturation | 10/137 | 8.56E-06 | 3.74E-02 |
| GO:0007517 | muscle organ development | 13/137 | 8.62E-06 | 3.77E-02 |
| GO:0014037 | Schwann cell differentiation | 5/137 | 8.86E-06 | 3.87E-02 |
| GO:0032660 | regulation of interleukin-17 production | 5/137 | 8.86E-06 | 3.87E-02 |
| GO:1904706 | negative regulation of vascular associated smooth muscle cell proliferation | 5/137 | 8.86E-06 | 3.87E-02 |
| GO:0008584 | male gonad development | 8/137 | 8.86E-06 | 3.87E-02 |
| GO:0044264 | cellular polysaccharide metabolic process | 7/137 | 8.90E-06 | 3.89E-02 |
| GO:0060537 | muscle tissue development | 13/137 | 9.08E-06 | 3.97E-02 |
| GO:0045670 | regulation of osteoclast differentiation | 6/137 | 9.11E-06 | 3.98E-02 |
| GO:0050918 | positive chemotaxis | 6/137 | 9.11E-06 | 3.98E-02 |
| GO:0072678 | T cell migration | 6/137 | 9.11E-06 | 3.98E-02 |
| GO:0030879 | mammary gland development | 8/137 | 9.33E-06 | 4.08E-02 |
| GO:0046546 | development of primary male sexual characteristics | 8/137 | 9.33E-06 | 4.08E-02 |
| GO:0032930 | positive regulation of superoxide anion generation | 4/137 | 9.48E-06 | 4.14E-02 |
| GO:0060252 | positive regulation of glial cell proliferation | 4/137 | 9.48E-06 | 4.14E-02 |
| GO:0150079 | negative regulation of neuroinflammatory response | 4/137 | 9.48E-06 | 4.14E-02 |
| GO:1902176 | negative regulation of oxidative stress-induced intrinsic apoptotic signaling pathway | 4/137 | 9.48E-06 | 4.14E-02 |
| GO:0043502 | regulation of muscle adaptation | 7/137 | 9.50E-06 | 4.15E-02 |
| GO:0060996 | dendritic spine development | 7/137 | 9.50E-06 | 4.15E-02 |
| GO:0072001 | renal system development | 11/137 | 9.53E-06 | 4.16E-02 |
| GO:0031345 | negative regulation of cell projection organization | 9/137 | 9.58E-06 | 4.19E-02 |
| GO:0040014 | regulation of multicellular organism growth | 6/137 | 9.94E-06 | 4.34E-02 |
| GO:0072577 | endothelial cell apoptotic process | 6/137 | 9.94E-06 | 4.34E-02 |
| GO:0048713 | regulation of oligodendrocyte differentiation | 5/137 | 1.01E-05 | 4.40E-02 |
| GO:0010212 | response to ionizing radiation | 8/137 | 1.03E-05 | 4.52E-02 |
| GO:0043279 | response to alkaloid | 7/137 | 1.08E-05 | 4.72E-02 |
| GO:0048662 | negative regulation of smooth muscle cell proliferation | 6/137 | 1.08E-05 | 4.73E-02 |
| GO:0010821 | regulation of mitochondrion organization | 9/137 | 1.09E-05 | 4.75E-02 |
| GO:0022408 | negative regulation of cell-cell adhesion | 9/137 | 1.09E-05 | 4.75E-02 |
| GO:0043267 | negative regulation of potassium ion transport | 5/137 | 1.14E-05 | 4.98E-02 |
| GO:1902042 | negative regulation of extrinsic apoptotic signaling pathway via death domain receptors | 5/137 | 1.14E-05 | 4.98E-02 |

Table 4: Novel gene ontology terms (n=43)

| ID | Description | Gene ratio | p-value | Adjusted p-value |
|---|---|---|---|---|
| GO:0007265 | Ras protein signal transduction | 10/24 | 5.91E-12 | 9.72E-09 |
| GO:0051056 | regulation of small GTPase mediated signal transduction | 8/24 | 3.92E-09 | 6.45E-06 |
| GO:0051222 | positive regulation of protein transport | 8/24 | 8.04E-09 | 1.32E-05 |
| GO:1904951 | positive regulation of establishment of protein localization | 8/24 | 1.13E-08 | 1.87E-05 |
| GO:0034599 | cellular response to oxidative stress | 7/24 | 8.23E-08 | 1.35E-04 |

| | | | | |
|---|---|---|---|---|
| GO:0046578 | regulation of Ras protein signal transduction | 6/24 | 1.26E-07 | 2.08E-04 |
| GO:1903829 | positive regulation of cellular protein localization | 7/24 | 1.48E-07 | 2.44E-04 |
| GO:0062197 | cellular response to chemical stress | 7/24 | 2.27E-07 | 3.74E-04 |
| GO:1905477 | positive regulation of protein localization to membrane | 5/24 | 4.90E-07 | 8.06E-04 |
| GO:2001233 | regulation of apoptotic signaling pathway | 7/24 | 5.74E-07 | 9.45E-04 |
| GO:0038127 | ERBB signaling pathway | 5/24 | 9.46E-07 | 1.56E-03 |
| GO:0006979 | response to oxidative stress | 7/24 | 1.15E-06 | 1.89E-03 |
| GO:0051205 | protein insertion into membrane | 4/24 | 1.29E-06 | 2.13E-03 |
| GO:0034614 | cellular response to reactive oxygen species | 5/24 | 2.07E-06 | 3.41E-03 |
| GO:0090316 | positive regulation of intracellular protein transport | 5/24 | 2.90E-06 | 4.76E-03 |
| GO:2001235 | positive regulation of apoptotic signaling pathway | 5/24 | 2.98E-06 | 4.89E-03 |
| GO:1905475 | regulation of protein localization to membrane | 5/24 | 4.06E-06 | 6.68E-03 |
| GO:1900739 | regulation of protein insertion into mitochondrial membrane involved in apoptotic signaling pathway | 3/24 | 4.61E-06 | 7.59E-03 |
| GO:1900740 | positive regulation of protein insertion into mitochondrial membrane involved in apoptotic signaling pathway | 3/24 | 4.61E-06 | 7.59E-03 |
| GO:0001844 | protein insertion into mitochondrial membrane involved in apoptotic signaling pathway | 3/24 | 7.18E-06 | 1.18E-02 |
| GO:0032388 | positive regulation of intracellular transport | 5/24 | 7.64E-06 | 1.26E-02 |
| GO:0045787 | positive regulation of cell cycle | 6/24 | 8.05E-06 | 1.32E-02 |
| GO:0007050 | cell cycle arrest | 5/24 | 9.86E-06 | 1.62E-02 |
| GO:0007229 | integrin-mediated signaling pathway | 4/24 | 9.88E-06 | 1.63E-02 |
| GO:0000302 | response to reactive oxygen species | 5/24 | 1.01E-05 | 1.66E-02 |
| GO:0001558 | regulation of cell growth | 6/24 | 1.13E-05 | 1.85E-02 |
| GO:0038093 | Fc receptor signaling pathway | 5/24 | 1.14E-05 | 1.87E-02 |
| GO:1901030 | positive regulation of mitochondrial outer membrane permeabilization involved in apoptotic signaling pathway | 3/24 | 1.15E-05 | 1.90E-02 |
| GO:0007173 | epidermal growth factor receptor signaling pathway | 4/24 | 1.65E-05 | 2.72E-02 |
| GO:0033157 | regulation of intracellular protein transport | 5/24 | 1.77E-05 | 2.91E-02 |
| GO:0014066 | regulation of phosphatidylinositol 3-kinase signaling | 4/24 | 1.87E-05 | 3.08E-02 |
| GO:0042770 | signal transduction in response to DNA damage | 4/24 | 2.25E-05 | 3.70E-02 |
| GO:0002429 | immune response-activating cell surface receptor signaling pathway | 6/24 | 2.42E-05 | 3.99E-02 |
| GO:0002757 | immune response-activating signal transduction | 6/24 | 2.42E-05 | 3.99E-02 |
| GO:0051204 | protein insertion into mitochondrial membrane | 3/24 | 2.48E-05 | 4.08E-02 |
| GO:1901028 | regulation of mitochondrial outer membrane permeabilization involved in apoptotic signaling pathway | 3/24 | 2.48E-05 | 4.08E-02 |
| GO:0000082 | G1/S transition of mitotic cell cycle | 5/24 | 2.64E-05 | 4.34E-02 |
| GO:0002433 | immune response-regulating cell surface receptor signaling pathway involved in phagocytosis | 4/24 | 2.67E-05 | 4.40E-02 |
| GO:0038096 | Fc-gamma receptor signaling pathway involved in phagocytosis | 4/24 | 2.67E-05 | 4.40E-02 |
| GO:0016049 | cell growth | 6/24 | 2.69E-05 | 4.43E-02 |
| GO:0097193 | intrinsic apoptotic signaling pathway | 5/24 | 2.77E-05 | 4.56E-02 |
| GO:0038094 | Fc-gamma receptor signaling pathway | 4/24 | 2.91E-05 | 4.78E-02 |

### 3 Pathways

Table 5: Pathways (n=45)

| Pathway | KEGG ID | p-value | Adjusted p-value | Genes |
|---|---|---|---|---|
| Alzheimer disease | path:hsa05010 | 7.67E-12 | 7.21E-10 | [APP, GSK3B, LRP1, PSEN2, LPL, PSEN1, IDE, TNF, APH1A, NCSTN, CASP3, APOE, NOS1, SNCA, MME, ADAM10, GRIN2B, TNFRSF1A, BACE1, BACE2, ADAM17, CDK5, IL1B, FAS, MAPT, GAPDH, CDK5R1] |
| Malaria | path:hsa05144 | 3.09E-10 | 2.90E-08 | [IL10, CXCL8, CR1, TGFB1, LRP1, IL18, TNF, ICAM1, IL6, CD40LG, IL1B, TLR9, CCL2, TLR4] |
| Chagas disease (American trypanosomiasis) | path:hsa05142 | 3.00E-08 | 2.82E-06 | [IL10, ACE, TGFB1, CXCL8, NOS2, TNF, TNFRSF1A, C3, IL6, PPP2R2B, IL1B, TLR9, CCL3, AKT1, CCL2, FAS, TLR4] |
| Tuberculosis | path:hsa05152 | 7.75E-08 | 7.28E-06 | [IL10, TGFB1, CR1, NOS2, VDR, IL18, TNF, TNFRSF1A, C3, IL1A, IL6, CASP3, IL1B, BCL2, TLR9, AKT1, HLA-DRA, BAX, CTSD, TLR4, HLA-DRB1, RAB7A] |
| Fluid shear stress and atherosclerosis | path:hsa05418 | 1.25E-07 | 1.17E-05 | [GSTO2, GSTO1, NOS3, CAV1, GSTP1, GSTT1, TNF, ICAM1, TNFRSF1A, VEGFA, IL1A, IL1B, BCL2, AKT1, HMOX1, CCL2, CTNNB1, TP53, SQSTM1] |
| HIF-1 signaling pathway | path:hsa04066 | 1.40E-07 | 1.32E-05 | [NOS2, NOS3, IGF1, MTOR, IGF1R, INS, VEGFA, IL6, TF, BCL2, AKT1, HMOX1, TIMP1, IL6R, TLR4, GAPDH] |
| Rheumatoid arthritis | path:hsa05323 | 1.72E-07 | 1.61E-05 | [CXCL8, TGFB1, MMP3, IL18, TNF, ICAM1, VEGFA, IL1A, IL6, IL1B, CCL3, CCL2, HLA-DRA, TLR4, HLA-DRB1] |
| Human cytomegalovirus infection | path:hsa05163 | 3.27E-07 | 3.08E-05 | [GSK3B, CXCL8, CDKN2A, PTGER2, HLA-A, PTGS2, TNF, MTOR, VEGFA, TNFRSF1A, IL6, SP1, IL1B, CASP3, CCL3, AKT1, CTNNB1, FAS, GNB3, CCL2, BAX, TP53, IL6R, SOS2] |
| Cholesterol metabolism | path:hsa04979 | 4.91E-07 | 4.61E-05 | [ABCA1, CETP, SOAT1, NPC1, LRP1, APOC1, APOA1, LPL, LRP2, APOE, LDLR] |
| Leishmaniasis | path:hsa05140 | 6.03E-07 | 5.67E-05 | [IL10, CR1, TGFB1, NOS2, PTGS2, TNF, C3, IL4, IL1A, IL1B, HLA-DRA, TLR4, HLA-DRB1] |
| Amyotrophic lateral sclerosis (ALS) | path:hsa05014 | 6.07E-07 | 5.71E-05 | [TOMM40, CASP3, BCL2, BAX, NOS1, TNFRSF1B, TNF, TP53, GRIN2B, TNFRSF1A, SOD1] |
| AGE-RAGE signaling pathway in diabetic complications | path:hsa04933 | 7.21E-07 | 6.78E-05 | [TGFB1, CXCL8, NOS3, TNF, AGER, ICAM1, VEGFA, IL1A, IL6, CASP3, IL1B, BCL2, AKT1, CCL2, BAX] |

| | | | | |
|---|---|---|---|---|
| African trypanosomiasis | path:hsa05143 | 1.04E-06 | 9.73E-05 | [IL10, IL6, IL1B, IL18, TLR9, APOA1, FAS, TNF, ICAM1] |
| Prion diseases | path:hsa05020 | 1.35E-06 | 1.27E-04 | [PRNP, IL1A, IL6, HSPA5, IL1B, BAX, FYN, NCAM2, SOD1] |
| PI3K-Akt signaling pathway | path:hsa04151 | 4.55E-06 | 4.28E-04 | [GSK3B, IRS1, FGF1, IGF1R, INS, RXRA, RELN, NTF3, AKT1, PCK1, IL6R, NTRK1, NGFR, NTRK2, BDNF, NOS3, IGF2, IGF1, MTOR, VEGFA, IL4, GH1, IL6, PPP2R2B, BCL2, GNB3, TP53, TLR4, SOS2] |
| Non-alcoholic fatty liver disease (NAFLD) | path:hsa04932 | 7.45E-06 | 7.01E-04 | [GSK3B, TGFB1, CXCL8, IRS1, TNF, EIF2S1, TNFRSF1A, INS, IL1A, IL6, RXRA, CASP3, IL1B, AKT1, BAX, FAS, IL6R] |
| Inflammatory bowel disease (IBD) | path:hsa05321 | 7.58E-06 | 7.13E-04 | [IL10, IL4, IL1A, IL6, TGFB1, IL1B, IL18, HLA-DRA, TNF, TLR4, HLA-DRB1] |
| Neurotrophin signaling pathway | path:hsa04722 | 7.68E-06 | 7.22E-04 | [NTRK1, NGFR, GSK3B, NTRK2, IRS1, BDNF, PSEN2, PSEN1, NTF3, BCL2, AKT1, BAX, TP53, SOS2, TP73] |
| Kaposi sarcoma-associated herpesvirus infection | path:hsa05167 | 1.10E-05 | 1.03E-03 | [GSK3B, CXCL8, EIF2AK2, HLA-A, PTGS2, MTOR, ICAM1, TNFRSF1A, VEGFA, C3, RCAN1, IL6, CASP3, AKT1, GNB3, BAX, CTNNB1, FAS, TP53] |
| Influenza A | path:hsa05164 | 1.25E-05 | 1.17E-03 | [GSK3B, CXCL8, IL18, EIF2AK2, TNF, EIF2S1, ICAM1, TNFRSF1A, IL1A, IL6, DDX39B, IL1B, AKT1, HLA-DRA, CCL2, FAS, TLR4, HLA-DRB1] |
| Prostate cancer | path:hsa05215 | 1.61E-05 | 1.51E-03 | [GSK3B, GSTP1, MMP3, IGF1, MTOR, IGF1R, INS, PLAU, BCL2, AKT1, CTNNB1, TP53, SOS2] |
| Serotonergic synapse | path:hsa04726 | 1.90E-05 | 1.78E-03 | [APP, MAOB, MAOA, ALOX15, HTR2C, HTR2A, PTGS2, SLC6A4, PTGS1, HTR6, CYP2D6, CASP3, ALOX5, GNB3] |
| Toxoplasmosis | path:hsa05145 | 1.90E-05 | 1.78E-03 | [IL10, TGFB1, NOS2, TNF, TNFRSF1A, CD40LG, CASP3, ALOX5, BCL2, AKT1, HLA-DRA, TLR4, LDLR, HLA-DRB1] |
| Allograft rejection | path:hsa05330 | 2.14E-05 | 2.01E-03 | [IL10, IL4, CD40LG, HLA-DRA, FAS, HLA-A, TNF, HLA-DRB1] |
| Ovarian steroidogenesis | path:hsa04913 | 2.63E-05 | 2.47E-03 | [LHCGR, FSHR, ALOX5, IGF1, PTGS2, CYP19A1, LDLR, IGF1R, INS] |
| Proteoglycans in cancer | path:hsa05205 | 3.31E-05 | 3.12E-03 | [TGFB1, CAV1, IGF2, ANK3, IGF1, HSPG2, TNF, ESR1, MTOR, VEGFA, IGF1R, PLAU, CASP3, AKT1, CTNNB1, FAS, TP53, TLR4, SOS2] |
| Pertussis | path:hsa05133 | 3.52E-05 | 3.31E-03 | [C4B, IL10, C3, IL1A, IL6, CXCL8, NOS2, CASP3, IL1B, TNF, TLR4] |

| | | | | |
|---|---|---|---|---|
| Hepatocellular carcinoma | path:hsa05225 | 3.65E-05 | 3.43E-03 | [GSK3B, TGFB1, GSTO2, GSTO1, CDKN2A, GSTP1, IGF2, GSTT1, MTOR, IGF1R, LRP6, AKT1, HMOX1, BAX, CTNNB1, TP53, SOS2] |
| Graft-versus-host disease | path:hsa05332 | 4.72E-05 | 4.43E-03 | [IL1A, IL6, IL1B, HLA-DRA, FAS, HLA-A, TNF, HLA-DRB1] |
| Type I diabetes mellitus | path:hsa04940 | 6.76E-05 | 6.36E-03 | [IL1A, IL1B, HLA-DRA, FAS, HLA-A, TNF, HLA-DRB1, INS] |
| Legionellosis | path:hsa05134 | 6.84E-05 | 6.43E-03 | [C3, IL6, CXCL8, CR1, CASP3, IL1B, IL18, TNF, TLR4] |
| Measles | path:hsa05162 | 9.97E-05 | 9.37E-03 | [GSK3B, EIF2AK2, EIF2S1, IL4, IL1A, IL6, IL1B, TLR9, AKT1, FAS, FYN, TP53, TLR4, TP73] |
| MAPK signaling pathway | path:hsa04010 | 1.05E-04 | 9.91E-03 | [NTRK1, NGFR, NTRK2, TGFB1, BDNF, IGF2, IGF1, FGF1, TNF, MAPK8IP1, VEGFA, IGF1R, TNFRSF1A, INS, IL1A, IL1B, CASP3, NTF3, AKT1, FAS, MAPT, TP53, SOS2] |
| Herpes simplex infection | path:hsa05168 | 1.23E-04 | 1.16E-02 | [TBP, EIF2AK2, HLA-A, TNF, EIF2S1, TNFRSF1A, C3, IL6, CASP3, IL1B, CDK1, TLR9, HLA-DRA, CCL2, FAS, TP53, HLA-DRB1] |
| Longevity regulating pathway | path:hsa04211 | 1.54E-04 | 1.44E-02 | [IRS1, AKT1, BAX, PPARG, IGF1, SOD2, SIRT1, TP53, MTOR, IGF1R, INS] |
| Hepatitis C | path:hsa05160 | 1.75E-04 | 1.64E-02 | [GSK3B, EIF2AK2, TNF, EIF2S1, TNFRSF1A, RXRA, PPP2R2B, CASP3, AKT1, CTNNB1, BAX, FAS, TP53, SOS2, LDLR] |
| NF-kappa B signaling pathway | path:hsa04064 | 2.51E-04 | 2.36E-02 | [CXCL8, CD40LG, PARP1, PLAU, IL1B, BCL2, PTGS2, TNF, TLR4, ICAM1, TN-FRSF1A] |
| TNF signaling pathway | path:hsa04668 | 2.58E-04 | 2.42E-02 | [IL6, CASP3, IL1B, MMP3, AKT1, CCL2, FAS, TNFRSF1B, PTGS2, TNF, ICAM1, TN-FRSF1A] |
| Hematopoietic cell lineage | path:hsa04640 | 3.03E-04 | 2.84E-02 | [IL4, IL1A, IL6, CR1, MME, IL1B, HLA-DRA, TNF, IL6R, HLA-DRB1, CD33] |
| Breast cancer | path:hsa05224 | 3.39E-04 | 3.19E-02 | [GSK3B, IGF1, FGF1, ESR1, ESR2, MTOR, IGF1R, LRP6, SP1, AKT1, BAX, CTNNB1, TP53, SOS2] |
| Regulation of lipolysis in adipocytes | path:hsa04923 | 3.54E-04 | 3.33E-02 | [IRS1, NPY, AKT1, ADRB1, ADRB2, PTGS2, PTGS1, INS] |
| Gastric cancer | path:hsa05226 | 3.90E-04 | 3.66E-02 | [GSK3B, ABCB1, TGFB1, FGF1, MTOR, LRP6, RXRA, BCL2, AKT1, BAX, CTNNB1, CTNNA3, TP53, SOS2] |
| Adipocytokine signaling pathway | path:hsa04920 | 4.08E-04 | 3.83E-02 | [RXRA, IRS1, NPY, AKT1, TN-FRSF1B, PCK1, TNF, MTOR, TNFRSF1A] |
| Salmonella infection | path:hsa05132 | 5.04E-04 | 4.74E-02 | [IL1A, IL6, CXCL8, NOS2, IL1B, IL18, CCL3, KLC1, TLR4, RAB7A] |

| | | | | |
|---|---|---|---|---|
| Colorectal cancer | path:hsa05210 | 5.04E-04 | 4.74E-02 | [GSK3B, TGFB1, CASP3, BCL2, AKT1, BAX, CTNNB1, TP53, SOS2, MTOR] |

### 4 Genes

Table 6: The novel discovered genes (n=24)

| Gene symbol | Gene name |
|---|---|
| RB1 | RB transcriptional corepressor 1 |
| SHC3 | SHC adaptor protein 3 |
| EP300 | E1A binding protein p300 |
| JAK2 | Janus kinase 2 |
| VAV1 | vav guanine nucleotide exchange factor 1 |
| YWHAB | tyrosine 3-monooxygenase/tryptophan 5-monooxygenase activation protein beta |
| CUL3 | cullin 3 |
| MAPK1 | mitogen-activated protein kinase 1 |
| GRB2 | growth factor receptor bound protein 2 |
| CDK2 | cyclin dependent kinase 2 |
| YWHAQ | tyrosine 3-monooxygenase/tryptophan 5-monooxygenase activation protein theta |
| EGFR | epidermal growth factor receptor |
| ABL1 | ABL proto-oncogene 1, non-receptor tyrosine kinase |
| YWHAZ | tyrosine 3-monooxygenase/tryptophan 5-monooxygenase activation protein zeta |
| CSNK2A1 | casein kinase 2 alpha 1 |
| TRIO | trio Rho guanine nucleotide exchange factor |
| SGSM3 | small G protein signaling modulator 3 |
| PPARD | peroxisome proliferator activated receptor delta |
| ARHGEF12 | Rho guanine nucleotide exchange factor 12 |
| RAB1A | RAB1A, member RAS oncogene family |
| FUT8 | fucosyltransferase 8 |
| LCP2 | lymphocyte cytosolic protein 2 |
| SPATA13 | spermatogenesis associated 13 |
| RACK1 | receptor for activated C kinase 1 |

Table 7: The whole list of AD-related genes (n=261)

| Gene symbol | Gene name |
|---|---|
| RB1 | RB transcriptional corepressor 1 |
| SHC3 | SHC adaptor protein 3 |
| CTNNB1 | catenin beta 1 |
| EP300 | E1A binding protein p300 |
| TP53 | tumor protein p53 |
| JAK2 | Janus kinase 2 |
| VAV1 | vav guanine nucleotide exchange factor 1 |
| YWHAB | tyrosine 3-monooxygenase/tryptophan 5-monooxygenase activation protein beta |
| AKT1 | AKT serine/threonine kinase 1 |
| CUL3 | cullin 3 |
| MAPK1 | mitogen-activated protein kinase 1 |
| SIRT1 | sirtuin 1 |
| GRB2 | growth factor receptor bound protein 2 |
| GSK3B | glycogen synthase kinase 3 beta |
| CDK2 | cyclin dependent kinase 2 |
| YWHAQ | tyrosine 3-monooxygenase/tryptophan 5-monooxygenase activation protein theta |
| EGFR | epidermal growth factor receptor |

| | |
|---|---|
| ABL1 | ABL proto-oncogene 1, non-receptor tyrosine kinase |
| CASP3 | caspase 3 |
| CDK1 | cyclin dependent kinase 1 |
| YWHAZ | tyrosine 3-monooxygenase/tryptophan 5-monooxygenase activation protein zeta |
| FYN | FYN proto-oncogene, Src family tyrosine kinase |
| CSNK2A1 | casein kinase 2 alpha 1 |
| ESR1 | estrogen receptor 1 |
| MTOR | mechanistic target of rapamycin kinase |
| APP | amyloid beta precursor protein |
| BCL2 | BCL2 apoptosis regulator |
| PPARG | peroxisome proliferator activated receptor gamma |
| PARP1 | poly(ADP-ribose) polymerase 1 |
| RXRA | retinoid X receptor alpha |
| TP73 | tumor protein p73 |
| HSPA5 | heat shock protein family A (Hsp70) member 5 |
| PSEN1 | presenilin 1 |
| CDKN2A | cyclin dependent kinase inhibitor 2A |
| VDR | vitamin D receptor |
| CD2AP | CD2 associated protein |
| CDK5 | cyclin dependent kinase 5 |
| NTRK1 | neurotrophic receptor tyrosine kinase 1 |
| SP1 | Sp1 transcription factor |
| ADRB2 | adrenoceptor beta 2 |
| TGFB1 | transforming growth factor beta 1 |
| IRS1 | insulin receptor substrate 1 |
| NR1H2 | nuclear receptor subfamily 1 group H member 2 |
| ESR2 | estrogen receptor 2 |
| LRRK2 | leucine rich repeat kinase 2 |
| TNF | tumor necrosis factor |
| EIF2AK2 | eukaryotic translation initiation factor 2 alpha kinase 2 |
| CHRNB2 | cholinergic receptor nicotinic beta 2 subunit |
| TNFRSF1A | TNF receptor superfamily member 1A |
| RAB7A | RAB7A, member RAS oncogene family |
| TRIO | trio Rho guanine nucleotide exchange factor |
| SGSM3 | small G protein signaling modulator 3 |
| LRP1 | LDL receptor related protein 1 |
| MAPK8IP1 | mitogen-activated protein kinase 8 interacting protein 1 |
| TLR4 | toll like receptor 4 |
| DYRK1A | dual specificity tyrosine phosphorylation regulated kinase 1A |
| SOS2 | SOS Ras/Rho guanine nucleotide exchange factor 2 |
| CAV1 | caveolin 1 |
| IL6 | interleukin 6 |
| PSEN2 | presenilin 2 |
| IGF1 | insulin like growth factor 1 |
| IL1B | interleukin 1 beta |
| PPARD | peroxisome proliferator activated receptor delta |
| IL6R | interleukin 6 receptor |
| DAPK1 | death associated protein kinase 1 |
| GAB2 | GRB2 associated binding protein 2 |
| VEGFA | vascular endothelial growth factor A |
| CHRNA3 | cholinergic receptor nicotinic alpha 3 subunit |
| NEDD9 | neural precursor cell expressed, developmentally down-regulated 9 |
| ARHGEF12 | Rho guanine nucleotide exchange factor 12 |
| ADAM10 | ADAM metallopeptidase domain 10 |
| APBA1 | amyloid beta precursor protein binding family A member 1 |
| CHRNA4 | cholinergic receptor nicotinic alpha 4 subunit |
| PICALM | phosphatidylinositol binding clathrin assembly protein |
| RAB1A | RAB1A, member RAS oncogene family |
| DNM2 | dynamin 2 |

| | |
|---|---|
| TBP | TATA-box binding protein |
| ADAM17 | ADAM metallopeptidase domain 17 |
| CST3 | cystatin C |
| FUT8 | fucosyltransferase 8 |
| ICAM1 | intercellular adhesion molecule 1 |
| APBB2 | amyloid beta precursor protein binding family B member 2 |
| NOS1 | nitric oxide synthase 1 |
| IL10 | interleukin 10 |
| APOE | apolipoprotein E |
| BIN1 | bridging integrator 1 |
| SNCA | synuclein alpha |
| CCL2 | C-C motif chemokine ligand 2 |
| IGF1R | insulin like growth factor 1 receptor |
| NGFR | nerve growth factor receptor |
| NRG1 | neuregulin 1 |
| CHRNA7 | cholinergic receptor nicotinic alpha 7 subunit |
| DNMBP | dynamin binding protein |
| CHRFAM7A | CHRNA7 (exons 5-10) and FAM7A (exons A-E) fusion |
| LCP2 | lymphocyte cytosolic protein 2 |
| DRD4 | dopamine receptor D4 |
| A2M | alpha-2-macroglobulin |
| BDNF | brain derived neurotrophic factor |
| PIN1 | peptidylprolyl cis/trans isomerase, NIMA-interacting 1 |
| BAX | BCL2 associated X, apoptosis regulator |
| PLAU | plasminogen activator, urokinase |
| UBQLN1 | ubiquilin 1 |
| INS | insulin |
| TNK1 | tyrosine kinase non receptor 1 |
| NOS3 | nitric oxide synthase 3 |
| CDK5R1 | cyclin dependent kinase 5 regulatory subunit 1 |
| C3 | complement C3 |
| SPATA13 | spermatogenesis associated 13 |
| NOS2 | nitric oxide synthase 2 |
| EIF2S1 | eukaryotic translation initiation factor 2 subunit alpha |
| EPHA1 | EPH receptor A1 |
| DDX39B | DExD-box helicase 39B |
| PTGS2 | prostaglandin-endoperoxide synthase 2 |
| CFTR | CF transmembrane conductance regulator |
| TLR9 | toll like receptor 9 |
| APOA1 | apolipoprotein A1 |
| HFE | homeostatic iron regulator |
| CXCL8 | C-X-C motif chemokine ligand 8 |
| ACHE | acetylcholinesterase (Cartwright blood group) |
| TNFRSF1B | TNF receptor superfamily member 1B |
| TFAM | transcription factor A, mitochondrial |
| TNFAIP1 | TNF alpha induced protein 1 |
| GRIN2B | glutamate ionotropic receptor NMDA type subunit 2B |
| CARD8 | caspase recruitment domain family member 8 |
| WWC1 | WW and C2 domain containing 1 |
| NTRK2 | neurotrophic receptor tyrosine kinase 2 |
| HTR2A | 5-hydroxytryptamine receptor 2A |
| AGER | advanced glycosylation end-product specific receptor |
| LRP6 | LDL receptor related protein 6 |
| NCSTN | nicastrin |
| RCAN1 | regulator of calcineurin 1 |
| TARDBP | TAR DNA binding protein |
| ABCA1 | ATP binding cassette subfamily A member 1 |
| RACK1 | receptor for activated C kinase 1 |
| SLC6A3 | solute carrier family 6 member 3 |

| | |
|---|---|
| SOD1 | superoxide dismutase 1 |
| IL18 | interleukin 18 |
| F2 | coagulation factor II, thrombin |
| CD40LG | CD40 ligand |
| HMOX1 | heme oxygenase 1 |
| KLK6 | kallikrein related peptidase 6 |
| GAPDH | glyceraldehyde-3-phosphate dehydrogenase |
| LDLR | low density lipoprotein receptor |
| COMT | catechol-O-methyltransferase |
| BCHE | butyrylcholinesterase |
| ADRB1 | adrenoceptor beta 1 |
| NPC1 | NPC intracellular cholesterol transporter 1 |
| RELN | reelin |
| HTR6 | 5-hydroxytryptamine receptor 6 |
| IL4 | interleukin 4 |
| IGF2R | insulin like growth factor 2 receptor |
| CLU | clusterin |
| IL1A | interleukin 1 alpha |
| GH1 | growth hormone 1 |
| MEOX2 | mesenchyme homeobox 2 |
| HLA-DRA | major histocompatibility complex, class II, DR alpha |
| NTF3 | neurotrophin 3 |
| GRN | granulin precursor |
| IDE | insulin degrading enzyme |
| FAS | Fas cell surface death receptor |
| SOD2 | superoxide dismutase 2 |
| ABCG1 | ATP binding cassette subfamily G member 1 |
| UCHL1 | ubiquitin C-terminal hydrolase L1 |
| HLA-A | major histocompatibility complex, class I, A |
| KLC1 | kinesin light chain 1 |
| CCL3 | C-C motif chemokine ligand 3 |
| TREM2 | triggering receptor expressed on myeloid cells 2 |
| PTGS1 | prostaglandin-endoperoxide synthase 1 |
| LRP2 | LDL receptor related protein 2 |
| GNB3 | G protein subunit beta 3 |
| S100B | S100 calcium binding protein B |
| LHCGR | luteinizing hormone/choriogonadotropin receptor |
| ANK3 | ankyrin 3 |
| FGF1 | fibroblast growth factor 1 |
| LPL | lipoprotein lipase |
| SERPINA3 | serpin family A member 3 |
| ECE1 | endothelin converting enzyme 1 |
| DHCR24 | 24-dehydrocholesterol reductase |
| MAPT | microtubule associated protein tau |
| APH1A | aph-1 homolog A, gamma-secretase subunit |
| TOMM40 | translocase of outer mitochondrial membrane 40 |
| HTR2C | 5-hydroxytryptamine receptor 2C |
| CTSD | cathepsin D |
| SORL1 | sortilin related receptor 1 |
| CHAT | choline O-acetyltransferase |
| CETP | cholesteryl ester transfer protein |
| CFH | complement factor H |
| BACE1 | beta-secretase 1 |
| TIMP1 | TIMP metallopeptidase inhibitor 1 |
| BACE2 | beta-secretase 2 |
| CRP | C-reactive protein |
| TIMP2 | TIMP metallopeptidase inhibitor 2 |
| SQSTM1 | sequestosome 1 |
| PCDH11X | protocadherin 11 X-linked |

| | |
|---|---|
| PRNP | prion protein |
| CR1 | complement C3b/C4b receptor 1 (Knops blood group) |
| HSPG2 | heparan sulfate proteoglycan 2 |
| PPP2R2B | protein phosphatase 2 regulatory subunit Bbeta |
| ABCA7 | ATP binding cassette subfamily A member 7 |
| ABCA2 | ATP binding cassette subfamily A member 2 |
| TF | transferrin |
| SOAT1 | sterol O-acyltransferase 1 |
| IREB2 | iron responsive element binding protein 2 |
| SERPINA1 | serpin family A member 1 |
| GSTP1 | glutathione S-transferase pi 1 |
| CYP19A1 | cytochrome P450 family 19 subfamily A member 1 |
| OGG1 | 8-oxoguanine DNA glycosylase |
| FSHR | follicle stimulating hormone receptor |
| PCK1 | phosphoenolpyruvate carboxykinase 1 |
| ALDH2 | aldehyde dehydrogenase 2 family member |
| CBS | cystathionine beta-synthase |
| CD33 | CD33 molecule |
| MTHFD1L | methylenetetrahydrofolate dehydrogenase (NADP+ dependent) 1 like |
| OLR1 | oxidized low density lipoprotein receptor 1 |
| DLST | dihydrolipoamide S-succinyltransferase |
| SST | somatostatin |
| HLA-DRB1 | major histocompatibility complex, class II, DR beta 1 |
| PON1 | paraoxonase 1 |
| NPY | neuropeptide Y |
| COX15 | cytochrome c oxidase assembly homolog COX15 |
| DKK1 | dickkopf WNT signaling pathway inhibitor 1 |
| MTR | 5-methyltetrahydrofolate-homocysteine methyltransferase |
| APOC1 | apolipoprotein C1 |
| IGF2 | insulin like growth factor 2 |
| ACE | angiotensin I converting enzyme |
| MAOB | monoamine oxidase B |
| GSTO1 | glutathione S-transferase omega 1 |
| MMP3 | matrix metallopeptidase 3 |
| NCAM2 | neural cell adhesion molecule 2 |
| MTHFR | methylenetetrahydrofolate reductase |
| CYP2D6 | cytochrome P450 family 2 subfamily D member 6 |
| BLMH | bleomycin hydrolase |
| TFCP2 | transcription factor CP2 |
| CP | ceruloplasmin |
| SLC30A6 | solute carrier family 30 member 6 |
| ALOX5 | arachidonate 5-lipoxygenase |
| COX10 | cytochrome c oxidase assembly factor heme A:farnesyltransferase COX10 |
| GSTT1 | glutathione S-transferase theta 1 |
| ABCB1 | ATP binding cassette subfamily B member 1 |
| TTR | transthyretin |
| MAOA | monoamine oxidase A |
| SIGMAR1 | sigma non-opioid intracellular receptor 1 |
| MME | membrane metalloendopeptidase |
| PON3 | paraoxonase 3 |
| SLC6A4 | solute carrier family 6 member 4 |
| GSTO2 | glutathione S-transferase omega 2 |
| APOD | apolipoprotein D |
| PTGER2 | prostaglandin E receptor 2 |
| CTNNA3 | catenin alpha 3 |
| MPO | myeloperoxidase |
| ALOX15 | arachidonate 15-lipoxygenase |
| LRRTM3 | leucine rich repeat transmembrane neuronal 3 |
| SORCS1 | sortilin related VPS10 domain containing receptor 1 |

| | |
|---|---|
| CYP46A1 | cytochrome P450 family 46 subfamily A member 1 |
| GOLM1 | golgi membrane protein 1 |
| PRND | prion like protein doppel |
| EXOC3L2 | exocyst complex component 3 like 2 |
| VSNL1 | visinin like 1 |
| CALHM1 | calcium homeostasis modulator 1 |
| MS4A4A | membrane spanning 4-domains A4A |
| COL25A1 | collagen type XXV alpha 1 chain |

### 5 Pathways significances

Table 8: Pathways significance for the entire network (n=41)

| Pathway | Centrality |
|---|---|
| Non-alcoholic fatty liver disease (NAFLD) | 7.61E-06 |
| Tuberculosis | 7.58E-06 |
| Human cytomegalovirus infection | 7.48E-06 |
| GE-RAGE signaling pathway in diabetic complications | 6.84E-06 |
| Influenza A | 6.60E-06 |
| Chagas disease (American trypanosomiasis) | 6.22E-06 |
| Rheumatoid arthritis | 5.94E-06 |
| MAPK signaling pathway | 5.80E-06 |
| Herpes simplex infection | 5.75E-06 |
| Pertussis | 5.54E-06 |
| TNF signaling pathway | 5.36E-06 |
| Kaposi sarcoma-associated herpesvirus infection | 5.32E-06 |
| Malaria | 5.04E-06 |
| Measles | 4.97E-06 |
| Inflammatory bowel disease (IBD) | 4.81E-06 |
| Fluid shear stress and atherosclerosis | 4.80E-06 |
| Legionellosis | 4.74E-06 |
| Proteoglycans in cancer | 4.64E-06 |
| Leishmaniasis | 4.61E-06 |
| Hepatitis C | 4.61E-06 |
| Toxoplasmosis | 4.59E-06 |
| Graft-versus-host disease | 4.50E-06 |
| African trypanosomiasis | 4.41E-06 |
| NF-kappa B signaling pathway | 4.16E-06 |
| Hematopoietic cell lineage | 4.02E-06 |
| PI3K-Akt signaling pathway | 3.98E-06 |
| Salmonella infection | 3.90E-06 |
| Colorectal cancer | 3.68E-06 |
| Gastric cancer | 3.43E-06 |
| Amyotrophic lateral sclerosis (ALS) | 3.28E-06 |
| Type I diabetes mellitus | 3.12E-06 |
| Hepatocellular carcinoma | 3.09E-06 |
| HIF-1 signaling pathway | 3.08E-06 |
| Alzheimer disease | 2.94E-06 |
| Prostate cancer | 2.78E-06 |
| Neurotrophin signaling pathway | 2.67E-06 |
| Longevity regulating pathway | 2.64E-06 |
| Breast cancer | 2.64E-06 |
| Allograft rejection | 2.25E-06 |
| Adipocytokine signaling pathway | 1.43E-06 |
| Prion diseases | 1.31E-06 |

Table 9: Pathways significance for cluster 1 (n=5)

| Pathway | Centrality |
| --- | --- |
| Non-alcoholic fatty liver disease (NAFLD) | 1.53E-06 |
| TNF signaling pathway | 1.47E-06 |
| Alzheimer disease | 1.15E-06 |
| Amyotrophic lateral sclerosis (ALS) | 1.12E-06 |
| Adipocytokine signaling pathway | 8.69E-07 |

Table 10: Pathways significance for cluster 2 (n=21)

| Pathway | Centrality |
| --- | --- |
| Tuberculosis | 3.74E-06 |
| Influenza A | 3.50E-06 |
| Rheumatoid arthritis | 3.32E-06 |
| Inflammatory bowel disease (IBD) | 3.26E-06 |
| Pertussis | 3.23E-06 |
| AGE-RAGE signaling pathway in diabetic complications | 3.17E-06 |
| Leishmaniasis | 3.09E-06 |
| Chagas disease (American trypanosomiasis) | 3.06E-06 |
| Malaria | 3.00E-06 |
| Herpes simplex infection | 2.89E-06 |
| Legionellosis | 2.79E-06 |
| Hematopoietic cell lineage | 2.60E-06 |
| Graft-versus-host disease | 2.60E-06 |
| African trypanosomiasis | 2.53E-06 |
| Salmonella infection | 2.42E-06 |
| Measles | 2.31E-06 |
| NF-kappa B signaling pathway | 2.06E-06 |
| Toxoplasmosis | 2.02E-06 |
| Type I diabetes mellitus | 1.81E-06 |
| Allograft rejection | 1.74E-06 |
| Prion diseases | 5.98E-07 |

Table 11: Pathways significance for cluster 3 (n=15)

| Pathway | Centrality |
| --- | --- |
| Human cytomegalovirus infection | 2.19E-06 |
| Hepatocellular carcinoma | 2.08E-06 |
| Prostate cancer | 2.06E-06 |
| Kaposi sarcoma-associated herpesvirus infection | 2.06E-06 |
| Gastric cancer | 2.02E-06 |
| PI3K-Akt signaling pathway | 2.01E-06 |
| Colorectal cancer | 2.01E-06 |
| Breast cancer | 2.01E-06 |
| MAPK signaling pathway | 1.92E-06 |
| Proteoglycans in cancer | 1.86E-06 |
| Longevity regulating pathway | 1.83E-06 |
| Neurotrophin signaling pathway | 1.74E-06 |
| Hepatitis C | 1.73E-06 |
| Fluid shear stress and atherosclerosis | 1.40E-06 |
| HIF-1 signaling pathway | 1.32E-06 |
